# Potent reaction hijacking inhibitors of *Plasmodium falciparum* asparagine tRNA synthetase

**DOI:** 10.64898/2026.05.15.725388

**Authors:** Xi Ye, Lisl Y. Esherick, Nutpakal Ketprasit, Sunil K. Narwal, Luiz C. Godoy, Nonlawat Boonyalai, Con Dogovski, Craig J. Morton, Tayla Rabie, Mufuliat Toyin Famodimu, Chia-Wei Tai, Tomas Yeo, Lena H. M. Le, Michael G. Leeming, Mariana Laureano De Souza, Elodie Chenu, Darren J. Creek, Michael J Delves, Lyn-Marie Birkholtz, James Duffy, Karen Lobb, Greg Durst, Marcus C.S. Lee, David A. Fidock, Jacquin C. Niles, Miles G. Siegel, Leann Tilley, Stanley C. Xie

## Abstract

Malaria remains one of the major threats to human health. Breakthrough drugs with high potency and low resistance risk are needed to combat the ever-increasing resistance to currently deployed antimalarials. Here, we explore a series of 4-amino-quinazoline-based sulfonamides, with drug-like physicochemical parameters and a synthetically accessible scaffold. Exemplars exhibit nanomolar potency against blood stage *Plasmodium* cultures, with up to 300-fold selectivity compared with a mammalian cell line. The compounds are also active against transmissible stages of *P. falciparum* and are refractory to resistance development. Targeted mass spectrometry reveals that the compounds act as reaction hijacking inhibitors targeting *P. falciparum* aminoacyl tRNA synthetases (aaRSs). Subtle changes to the chemical structure switch the main target from cytoplasmic tRNA threonine synthetase (*Pf*ThrRS) to cytoplasmic asparagine synthetase (*Pf*AsnRS), a change that is associated with increased potency and selectivity. The target preference was confirmed by selective knock-down of different *P. falciparum* aaRSs and by tolerance selection in a mutator line. Consistent with aaRS targets, exemplar compounds activate the amino acid starvation response. Recombinant enzyme inhibition and thermal stabilisation assays confirm the susceptibility of *Pf*AsnRS to reaction hijacking and show that human AsnRS is less susceptible. A molecular model of Asn-tRNA-bound *Pf*AsnRS reveals that a potent hijacker adopts a pose similar to adenosine 5’-monophosphate (AMP). An AlphaFold model of the native *Pf*AsnRS dimer helps explain the tolerance-conferring effect of a mutation at the dimer interface.

## Introduction

Malaria remains a major cause of illness and mortality. For example, in 2024, over 260 million malaria cases and 600,000 deaths were reported, with numbers remaining high following COVID19-related disruptions (1). Current antimalarial treatments are rapidly losing efficacy (2), with ∼50% treatment failure of first-line artemisinin combination therapies reported in some regions of South East Asia (2). Worryingly, partial resistance to artemisinin and its derivatives has been clinically validated in Africa (3, 4), where most malaria deaths occur. With the current malaria vaccine achieving only 30% efficacy in reducing severe cases (5), new drug candidates are needed that have novel modes of action and are refractory to the development of resistance (6).

Aminoacyl-tRNA synthetases (aaRSs) are responsible for conjugating the correct amino acid onto its cognate tRNA, thus supplying charged tRNA for protein translation. As *Plasmodium falciparum* is a fast-growing parasite that relies on protein synthesis, aaRSs are considered important drug targets. Our recent work has revealed that some *P. falciparum* aaRSs are susceptible to inhibition via an unusual process called reaction hijacking (7–9). A series of sulfamoyl adenosine 5’-monophosphate (AMP) mimics have been shown to attack the ester bond of the enzyme-bound charged tRNA, leading to the formation of inhibitory sulfamoyl-amino acid adducts (7, 8). The generation of a tight binding adduct within the active site inactivates the enzyme.

A series of pyrazolopyrimidine sulfamates, with a 7-position substituent, showed potent activity against *P. falciparum* blood stage parasites and good selectivity compared with mammalian cell lines (7, 8). These compounds were shown to hijack *P. falciparum* tyrosyl-tRNA synthetase (*Pf*TyrRS), generating the relevant tyrosine adduct. Importantly, human TyrRS does not catalyse the hijacking reaction (7, 8). Exemplar ML471 exhibits single dose oral efficacy in a humanised mouse model of *P. falciparum* malaria (7), the gold standard for further development of antimalarials. Two concerns with ML471 were a propensity for resistance development, and a highly polar scaffold that limited oral bioavailability (10).

A second series of sulfamoyl AMP-mimics, including the 4-aminothieno-pyrimidine benzene sulfonamide, OSM-S-106, exhibits activity against parasite cultures and low mammalian cell toxicity (11). While OSM-S-106 is structurally divergent from AMP and ML471, targeted mass spectrometry of OSM-S-106-treated *P. falciparum* cultures confirmed that it acts as a reaction hijacking inhibitor. In this case, asparagine-tRNA synthetase (*Pf*AsnRS) was identified as the major hijacking target, with *P. falciparum* GlyRS and *P. falciparum* AlaRS as minor targets (11). OSM-S-106 exhibited a low propensity for resistance development (11). However, exploration of several substitutions of the aminothienopyrimidine core, the pendant aromatic ring and the primary sulfonamide did not identify compounds more potent than OSM-S-106, suggesting exploration of other scaffolds was warranted.

Here we continue our studies of sulfamoyl AMP-mimics with a scaffold hop from the aminothieno-pyrimidine system to an aminoquinazoline-based system. Structure-activity relationship (SAR) analysis of different substitutions identified 4-aminoquinazoline sulfonamides (4AQS) that exhibit very potent (nanomolar) activity against *P. falciparum* in cellular assays. The compounds exhibit drug-like physicochemical parameters, low resistance propensity and a synthetically accessible scaffold. Targeted mass spectrometry and conditional knock-down studies identified aaRS targets of the 4AQS series in *P. falciparum*. Interestingly, while the initial less active compound targeted *P. falciparum* threonyl-tRNA synthetase (*Pf*ThrRS) and *Pf*AsnRS with similar efficiency, more potent compounds preferentially targeted *Pf*AsnRS, plus additional minor targets.

## Results

### Synthesis of the 4AQS series and SAR analysis against P. falciparum and HepG2 mammalian cells

We explored a series of 4AQS compounds (Table 1; Supplementary Figure 1) as potential reaction hijacking inhibitors. Synthesis was achieved using the 4-aminoquinazoline synthesis protocol, as delineated in Scheme 1 and the Supplementary Information (Chemistry Methods). In a representative example, compound **3** (LG-0020420) (hereafter referred to as LG420) was synthesized by the following procedure: A mixture of 3-(4,4,5,5-tetramethyl-1,3,2-dioxaborolan-2-yl)benzenesulfonamide (120 mg, 424 μmol), 7-bromoquinazolin-4-amine (100 mg, 446 μmol), Pd(dppf)Cl_2_ (30.0 mg, 41.0 μmol) and K_2_CO_3_ (180 mg, 1.30 mmol) in dioxane (1.5 mL) and H_2_O (0.4 mL) was degassed and purged with N_2_ 3 times, then the mixture was stirred at 80°C for 12 h under N_2_ atmosphere. The mixture was filtered and the solid was collected, the filter cake was washed with water (4 mL) and MeOH (3 mL), then triturated with DMF (added 2 drops TFA) to give 3-(4-aminoquinazolin-7-yl)benzenesulfonamide (38.8 mg, 129 μmol, 28.9% yield) as a pale yellow solid. All final products were synthesized in sufficient quantities to allow for characterization by 1H NMR and LC/MS, typically a minimum of 10 mg final product, and appeared stable in the solid state with respect to degradation under ambient conditions (Supplementary Information). Haloaryl amino acids that are not commercially available, as well as authentic samples of amino acid adducts, were generated for sulfonamides LG-0020957 (Asn-420); LG-0020445 (Thr-420); and LG-0021200 (Asn-866), were prepared as described in Supplementary Information.

**Table 1.** Activities of 4AQS compounds as inhibitors of asexual parasite growth and toxicity in a mammalian cell line. n = Number of biological repeats. Data values represent mean ± SEM. Full LG designations are provided. Medicines for Malaria Venture (MMV) designations are in brackets.

| Structure |  |  |  |  |  |  |  |  |
| --- | --- | --- | --- | --- | --- | --- | --- | --- |
| Compound ID | **LG-0020420**<br>(MMV2323511) | **LG-0020866**<br>(MMV2480572) | **LG-0021041**<br>(MMV2480986) | **LG-0021685**<br>(MMV2482269) | **LG-0021877**<br>(MMV2482899) | **LG-0022521**<br>(MMV2503069) | **LG-0022563**<br>(MMV2503071) | **LG-0022677**<br>(MMV2503074) |
| 3D7 IC<sub>50</sub> (nM) | 265 ± 5 (n=3) | 37 ± 8 (n=3) | 100 ± 17 (n=3) | 12 ± 2 (n=3) | 257 ± 33 (n=3) | 73 ± 5 (n=3) | 9 ± 1 (n=3) | 3.1 ± 0.6 (n=4) |
| Dd2 IC<sub>50</sub> (nM) | 295 ± 25 (n=3) | 40 ± 9 (n=3) | 158 ± 26 (n=3) | 10 ± 1 (n=3) | 249 ± 26 (n=3) | 79 ± 4 (n=3) | 11 ± 1 (n=3) | 4.9 ± 1.3 (n=4) |
| HepG2 IC<sub>50</sub> (nM) | 11,900 ± 1,700 (n=3) | 1,210 ± 410 (n=3) | 1,830 ± 250 (n=3) | 300 ± 46 (n=3) | >25,000 (n=3) | 8,340 ± 1,380 (n=3) | 2,710 ± 460 (n=3) | 428 ± 24 (n=4) |
| Ratio HepG2/3D7 | 45 | 33 | 18 | 25 | >97 | 114 | 301 | 138 |
| SONH<sub>2</sub> pKa | 10.14 | 8.89 | 8.66 | 7.49 | 10.14 | 10.11 | 8.89 | 7.49 |

**Scheme 1:**
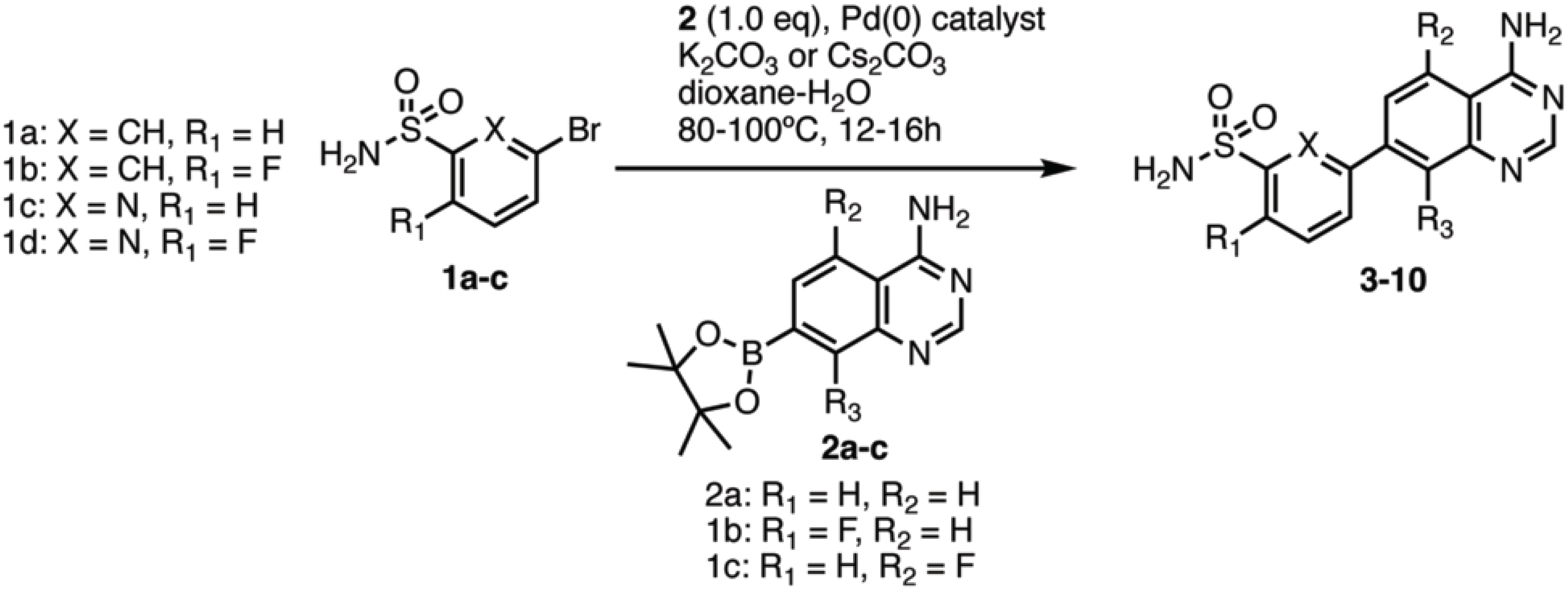
General 4AQS synthesis.

For the initial derivative, LG420, the aminothienopyrimidine core of OSM-S-106 (11) is replaced with a 4-aminoquinazoline core. LG420 exhibits reasonable activity in cellular assays against the *P. falciparum* laboratory strains, 3D7 (IC_50_ = 265 nM) and Dd2 (IC_50_ = 295 nM) (Table 1). LG420 shows lower activity against the mammalian cell line, HepG2, with an IC_50_ value of 11,900 nM, representing 45-fold selectivity (Table 1).

In an attempt to increase the nucleophilicity of the primary sulfonamide warhead, we substituted the phenyl sulfonamide for the sterically equivalent pyridine sulfonamide LG-0020866 (LG866, Table 1). This modification lowers the pKa from a calculated value of 10.14 to 8.89 (Table 1). LG866 exhibits 7-fold enhanced activity against 3D7 (IC_50_ = 37 nM) and Dd2 (IC_50_ = 40 nM) compared with LG420 (Table 1). LG866 also shows increased toxicity to HepG2 (33-fold selectivity) (Table 1).

We next explored fluorination of the phenyl ring of LG420 in the 6’ position as an alternative strategy to increase nucleophilicity (LG-0021041; LG041; Table 1). Compared with LG420, LG041 exhibits increased activity against 3D7 (IC_50_ = 100 nM) and Dd2 (IC_50_ = 158 nM) (Table 1), however this modification also increases toxicity and slightly reduces the cellular selectivity (HepG2/3D7 = 18-fold) (Table 1).

LG-0021685 (LG685; Table 1) combines the 2’ pyridine sulfonamide with the 6’ fluorination. Combining these modifications further reduces the sulfonamide pKa and shows substantially increased activity against 3D7 (IC_50_ = 12 nM) and Dd2 (IC_50_ = 10 nM; Table 1), with similar cellular selectivity as LG041 (HepG2/3D7 = 25-fold).

We next turned our attention to substitutions on the 4-aminoquinazoline ring (adenine binding site) at the 5- or 8-position (LG-0021877; LG877 and LG-0022521; LG521; Table 1), as these substitutions were recently shown to increase potency and selectivity in dual site sulfamoyl AMP mimic-amino acid adducts targeting a bacterial ProRS (12). Compared with LG420, LG521 exhibits increased activity against 3D7 (IC_50_ = 73 nM) and Dd2 (IC_50_ = 79 nM) (Table 1), with a more marked improvement in cellular selectivity (HepG2/3D7 = 114-fold) (Table 1). LG877 exhibits potency similar to LG420 against *P. falciparum* (IC_50_ = 257 nM) and Dd2 (IC_50_ = 249 nM) but does show much improved HepG2 toxicity (IC_50_ >25,000 nM).

Building on the lower HepG2 toxicity of LG877, and on information from a study of dual site inhibitors of ProRS (12), we combined the 2’ position pyridine sulfonamide with 5-position fluorination of the quinazoline ring (LG-0022563; LG563; Supplementary Figure 1). LG563 exhibits very potent activity against 3D7 (IC_50_ = 9 nM) and Dd2 (IC_50_ = 11 nM) (Table 1), and maintains cellular selectivity (HepG2/3D7 = 301-fold).

Finally, we explored the triple combination of fluorination at both the 5 and 6’ positions combined with the 2’-pyridyl sulfonamide (LG-0022677; LG677; Table 1). LG677 exhibits low nanomolar activity against 3D7 (IC_50_ = 3.1 nM) and Dd2 (IC_50_ = 4.9 nM) (Table 1), while still maintaining >100-fold cellular selectivity (HepG2/3D7 = 138-fold).

### LG677 and LG563 exhibit activity against transmissible forms of P. falciparum

LG677 and LG563 also exhibit potency against immature gametocytes (IC_50_48h_ = 199 nM and 224 nM, respectively), with lower activity against mature stage gametocytes (Supplementary Table 1). The assay was validated with known inhibitors, Methylene Blue and the *Plasmodium* phosphatidylinositol 4-kinase (PI4K) inhibitor, MMV390048. LG677 and LG563 also inhibit the fertility of male gametes (IC_50_48h_ = 373 nM and 658 nM, respectively) with weaker activity against female gametes (Supplementary Table 1). The positive control, Cabamiquine (also known as M5717 or DDD107498), exhibited potent activity against both gametocyte sexes, consistent with a previous report (13).

### LG866 exhibits a low propensity for resistance development

*In vitro* evolution under inhibitor pressure, followed by whole-genome sequencing is a standardised method to explore *P. falciparum*’s propensity for developing resistance and to identify or validate the antimalarial drug target(s) (14, 15). For these studies, a single-step selection was performed, with an initial inoculum of 10^9^ parasites (Dd2-B2 strain) across three flasks, at a concentration of 3 x IC_90_ (285 nM). No recrudescent parasites were observed over a 60-day selection period. Based on these studies, we conclude that the Minimum Inoculum for Resistance (MIR) value for LG866 is >10^9^ (Supplementary Table 2).

Following the MIR studies, selections were initiated using the Dd2-Polδ parasite line, which has a mutation rate that is ∼8-10 fold higher than Dd2-B2 (16). Two flasks, each with an inoculum of 10^9^ parasites, were exposed to a pulsing protocol. Recrudescence was observed on day 42 in both flasks, representing a predicted MIR of ∼4×10^9^ (Supplementary Table 2). The recrudescent lines exhibited a low level of resistance, showing a 6-8-fold increase in IC_50_ compared to the parental Dd2-B2 line (Supplementary Figure 2A and Supplementary Table 3). The recrudescent lines were subjected to whole-genome sequencing, which identified a missense mutation in both flasks 1 and 2 (A210T) within PF3D7_0211800 on chromosome 2, encoding the cytoplasmic *P. falciparum* AsnRS (Supplementary Table 4).

### 4AQS series compounds do not enrich lines from a resistant parasite set

The AReBar (Antimalarial Barcode Sequencing) assay comprises a pool of 53 barcoded parasites that have mutations in genes recognized as antimalarial targets or resistance determinants (Supplementary Table 5) (17, 18). Outgrowth of specific barcode lines offers insight into the susceptibility to resistance of the compound of interest. LG420 and LG866 were applied at 3 x IC_50_ values to the AReBar library, leading to decreased parasite growth (Supplementary Figure 2B). No enrichment was observed above the threshold for any mutant line (Supplementary Figure 2C), even though the library includes the *Pf*AsnRS^R487S^ mutant identified upon selection against the known *Pf*AsnRS hijacker, OSM-S-106 (11). One mutant harbouring an R2180P single nucleotide polymorphism in ATP-binding cassette (ABC) transporter ABCI3 (PF3D7_0319700) was present at relatively high abundance and gave a signal close to the threshold (Supplementary Figure 2C-D) but on testing, the IC_50_ value for this line was equivalent to wildtype (Supplementary Figure 2E), indicting it does not confer resistance. As a positive control, the prolyl tRNA synthetase inhibitor halofuginone highly enriched the *Pf*ProRS^R482H^ mutant (Supplementary Figure 2D).

We also directly tested the response of a transfectant harbouring the *Pf*AsnRS^R487S^ mutation. This line exhibited two-fold decreased sensitivity to OSM-S-106 in agreement with our previous report (11). The mutant line does not show reduced sensitivity to LG420 but does exhibit 2.5-fold reduced sensitivity to LG866 (Supplementary Figure 2F). The low level resistance explains why the *Pf*AsnRS^R487S^ mutant was not selected when the pool was pressured at 3 x IC_50_.

### LG420 and LG866 induce the amino acid starvation response

Inhibition of aaRSs leads to a build-up of uncharged tRNA, which in turn leads to eIF2α phosphorylation (7, 19, 20). Exposure to different concentrations of LG420, LG866, LG563 and LG677 triggers eIF2α phosphorylation, to an extent that closely reflects their potency as inhibitors of the growth of *P. falciparum* (Figure 1, Supplementary Figure 3). The known ThrRS inhibitor, borrelidin (21), was used as a positive control.

**Figure 1.**
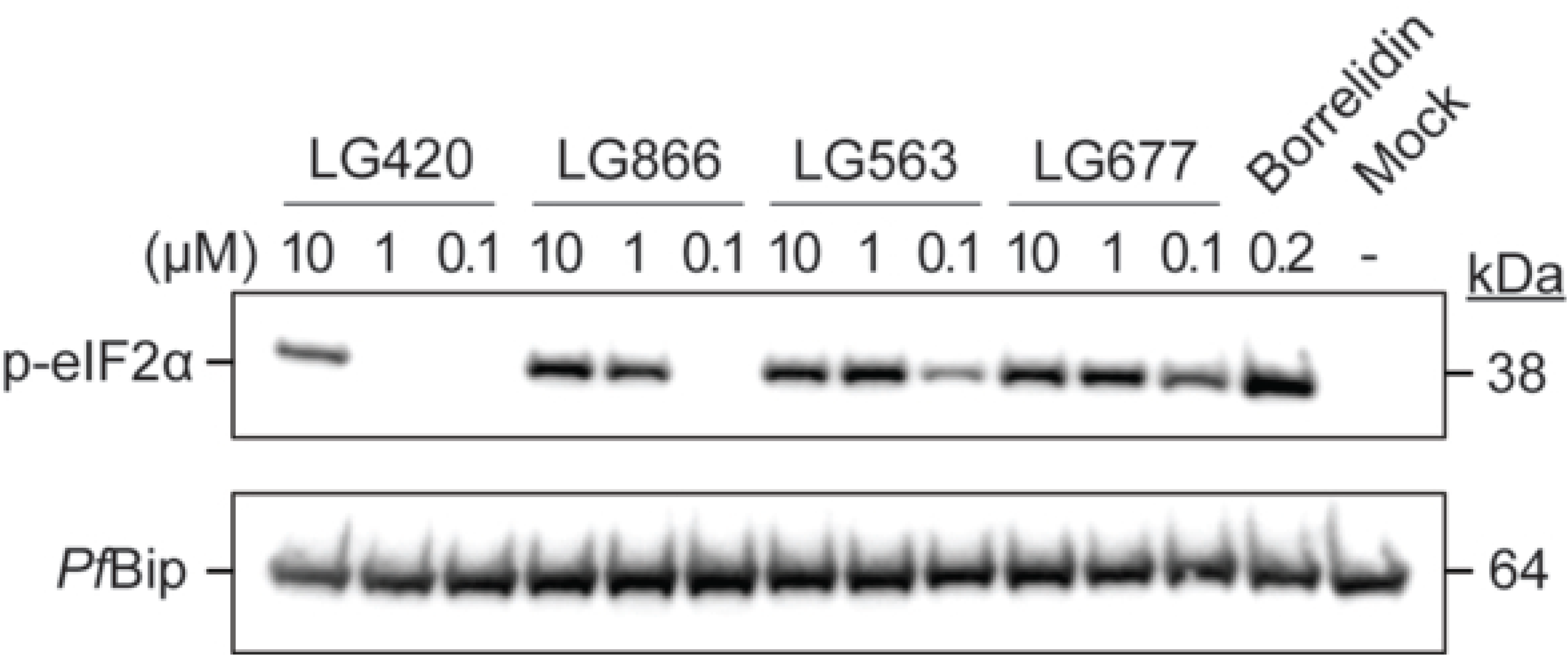
aaRS inhibition induces the amino acid starvation response via eIF2α phosphorylation. Trophozoite-stage *P. falciparum* 3D7 cultures were exposed to 0.2 µM borrelidin and different concentrations of LG420, LG866, LG563, and LG677 for 3 h. Western blot analysis was performed on parasite extracts to detect phosphorylated eIF2α. *Pf*BiP served as the loading control. See additional blots from independent experiments in Supplementary Figure 3.

### 4AQS series compounds hijack P. falciparum aaRSs to form amino acid adducts

Reaction-hijacking sulfamoyl AMP mimics attack the activated ester bonds of the enzyme-bound aminoacyl tRNA to form amino acid adducts that remain bound to the enzyme (8). We used targeted mass spectrometry to search for potential sulfonamide-amino acid conjugates formed in *P. falciparum* cultures that had been treated with 4AQS series compounds (10 μM, for 3 h). Following Folch extraction of lysates, the aqueous phase was subjected to liquid chromatography-coupled with mass spectrometry (LC-MS) and the anticipated masses were interrogated. For LG420, the extract yielded strong signals for both Thr-LG420 and Asn-LG420, with precursor ion and fragmentation spectra consistent with those of the synthetic Asn-LG420 and Thr-LG420 conjugates (Figure 2A, B; Table 2). By contrast, for the more potent derivatives LG866, LG685, LG521, LG041, LG563 and LG677, the Asn-conjugate was the predominant species, with other adducts, such as the Thr adduct, giving signals about 10x lower (Figure 2B,C; Table 2, Supplementary Figure 4-6). These data indicate that *Pf*AsnRS is an important target for 4AQS compounds and that subtle changes in chemical structure can broaden or change the target preference.

**Figure 2.**
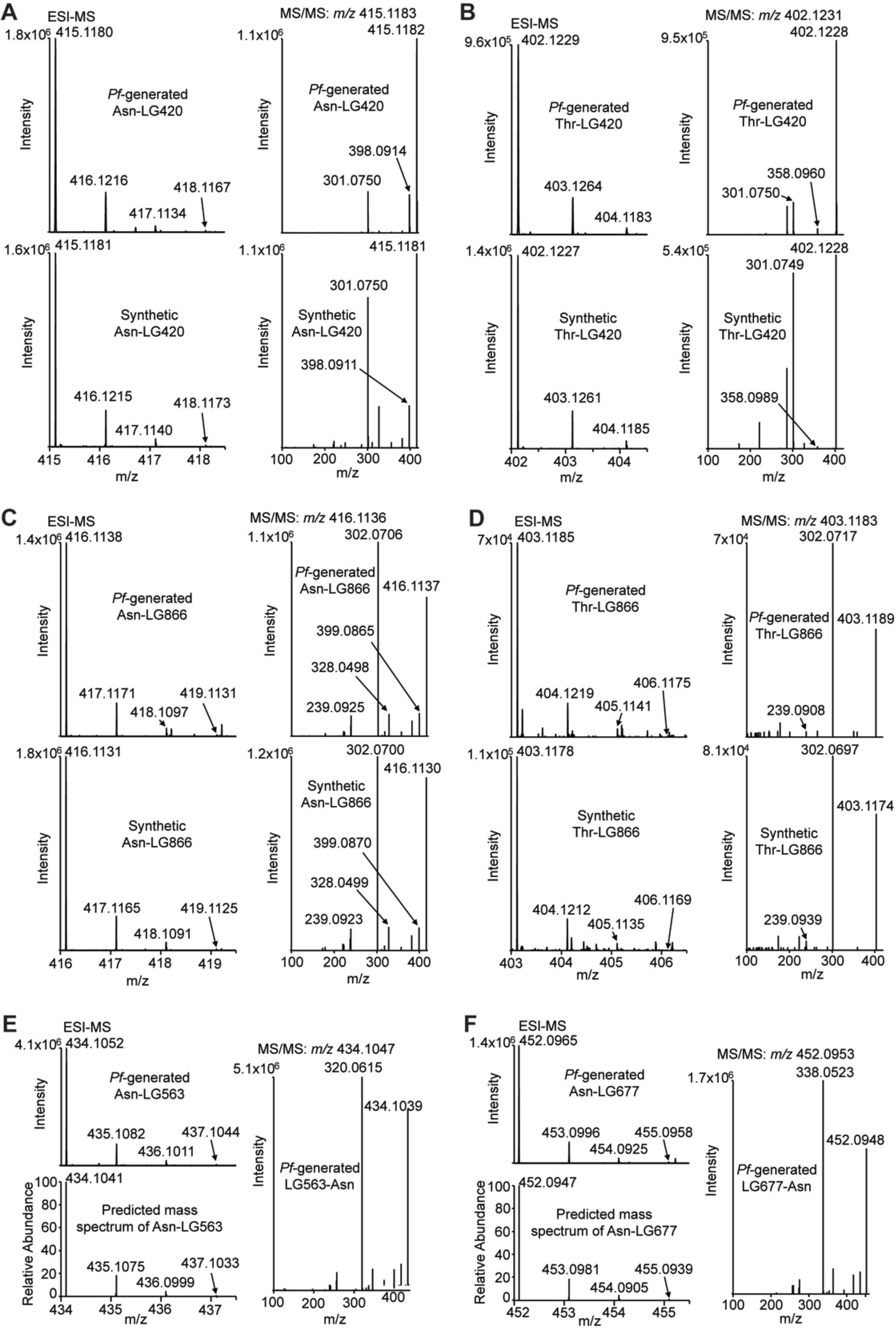
Targeted mass spectrometry identifies amino acid adducts in *P. falciparum.* Late trophozoite stage *P. falciparum* 3D7 cultures were exposed to 10 μM LG420, LG866, LG563 and LG677 for 3 h. Extracts containing amino acid adducts were analysed by LC-MS. Mass spectra of Asn-LG420 (*m/z* = 415.1183) (A), Thr-LG420 (*m/z* = 403.1231) (B), Asn-LG866 (*m/z* = 416.1136) (C), Thr-LG866 (*m/z* = 403.1183) (D), Asn-LG563 (*m/z* = 434.1047) (E) and Asn-LG677 (*m/z* = 452.0953) (F) are shown in the left panels, and their MS/MS fragmentation spectra are shown in the right panels. For LG420 and LG866 (A,B), the top panels show adducts generated by *P. falciparum*, and bottom panels show synthetic standards spiked into untreated parasite lysate (1 μM Asn-LG420, Thr-LG420 and Asn-LG866; 0.3 μM Thr-LG866). For LG563 and LG677 (C,D), the detected (top left panels) and predicted (bottom left panels) mass spectra, and MS/MS fragmentation spectra (right panels) are shown.

**Table 2.** Targets of pyrazolopyrimidine sulfamates as detected by targeted mass spectrometry and activities in biochemical assays. Adducts detected by mass spectrometry and relative signal strengths are denoted. ATP consumption by *Pf*AsnRS and *Hs*AsnRS (25 nM) was measured following incubation with 100 μM 4AQS compound in the presence of 200 μM L-asparagine, 10 μM ATP, 1 unit/mL inorganic pyrophosphatase, and 2.5 mg/mL *E. coli* tRNA at 37°C for 1 h. Percentage inhibition was calculated relative to the AsnRS plus *Ec*tRNA with no compound (positive control). For DSF experiments, *Pf*AsnRS and *Hs*AsnRS were incubated at 37°C for 3 h in the apo form or with 100 μM compound (3 μM AsnRS, 40 μM ATP, 80 μM L-asparagine, 60 μM *Ec*tRNA). Tm_app_ values were measured in the presence of 10X SYPRO Orange with (*Pf*AsnRS) or without (*Hs*AsnRS) 0.001% SDS. ΔTm_app_ was calculated as Tm_app_ (AsnRS, with ATP, Asn, *Ec*tRNA and pro-inhibitor) − Tm_app_ (AsnRS with ATP, Asn and *Ec*tRNA). n = Number of biological repeats.

| LG ID | LG420 | LG866 | LG041 | LG685 | LG521 | LG563 | LG677 |
| --- | --- | --- | --- | --- | --- | --- | --- |
| **Adducts detected by MS ID** | Asn, Thr | Asn (strong), Thr, Lys, Ala, Gly (minor) | Asn (strong), Thr (minor) | Asn (strong), Thr, Gly, Lys (minor) | Asn (strong), Thr, Lys (minor) | Asn (strong), Ala, Thr, Gly, Lys (minor) | Asn (strong) Gly, Thr, Ala, Lys (minor) |
| **% Inhibition of *Pf*AsnRS (100 μM compound)** | 7.4 ± 2.8 (n=6) | 12.0 ± 2.5 (n=3) | 10.2 ± 2.3 (n=3) | 14.1 ± 1.5 (n=3) | 21.6 ± 3.1 (n=3) | 29.9 ± 0.6 (n=3) | 47.6 ± 2.9 (n=4) |
| **% Inhibition of *Hs*AsnRS (100 μM compound)** | -2.2 ± 0.7 (n=5) | -0.6 ± 0.4 (n=4) | -3.7 ± 1.2 (n=4) | 1.3 ± 2.5 (n=3) | 5.0 ± 0.8 (n=3) | -3.4 ± 2.1 (n=5) | -1.4 ± 1.8 (n=5) |
| **DSF $\Delta T_{m_{app}}$ *Pf*AsnRS (100 μM compound)** | 0.8 ± 0.2 (n=4) | 1.5 ± 0.1 (n=3) | 1.6 ± 0.1 (n=3) | 1.8 ± 0.1 (n=4) | 1.8 ± 0.1 (n=4) | 1.2 ± 0.1 (n=3) | 2.1 ± 0.1 (n=3) |
| **DSF $\Delta T_{m_{app}}$ *Hs*AsnRS (100 μM compound)** | -0.3 ± 0.0 (n=3) | -0.2 ± 0.0 (n=3) | -0.3 ± 0.0 (n=3) | 0.0 ± 0.1 (n=3) | 0.0 ± 0.1 (n=3) | -0.2 ± 0.0 (n=3) | -0.2 ± 0.0 (n=3) |

### Conditional knock-down confirms PfAsnRS and PfThrRS as important targets

The TetR/DOZI-RNA aptamer module can be used to conditionally regulate the expression of *P. falciparum* aaRSs (22). We used the system to knock-down the expression of the two main targets that were identified by mass spectrometry, namely *Pf*AsnRS (PF3D7_0211800) and *Pf*ThrRS (PF3D7_1126000). The oligonucleotides used for gene knockdown donor vector construction are listed in Supplementary Table 6. As anticipated, parasite viability assays revealed that knockdown of *Pf*AsnRS or *Pf*ThrRS affects parasite growth (Supplementary Figure 7), confirming that these enzymes are essential for blood stage development. Upon knockdown of *Pf*AsnRS, the parasites exhibited a 3.4-fold and 12-fold enhancement in susceptibility to LG420 and LG866, respectively, compared to the control (Figure 3A, Supplementary Figure 8), confirming *Pf*AsnRS as an important target of these compounds. Upon knockdown of *Pf*ThrRS, the parasites exhibited a 41-fold enhancement in susceptibility to LG420, but no change in susceptibility to LG866, consistent with the suggestion that *Pf*ThrRS is an important target for LG420. Interestingly, for LG041, knockdown of either *Pf*ThrRS or *Pf*AsnRS enhances susceptibility, while, in agreement with the mass spectrometry-based Target ID data, *Pf*AsnRS is the main susceptibility determinant for the more potent compounds, LG685, LG563 and LG677. Similarly, for OSM-S-106, *Pf*AsnRS appears to be the major target (Figure 3B), as expected from previous studies (11). These data are consistent with the suggestion that both *Pf*AsnRS and *Pf*ThrRS are potential targets of this series but that selective targeting of *Pf*AsnRS is associated with higher anti-plasmodial potency.

**Figure 3.**
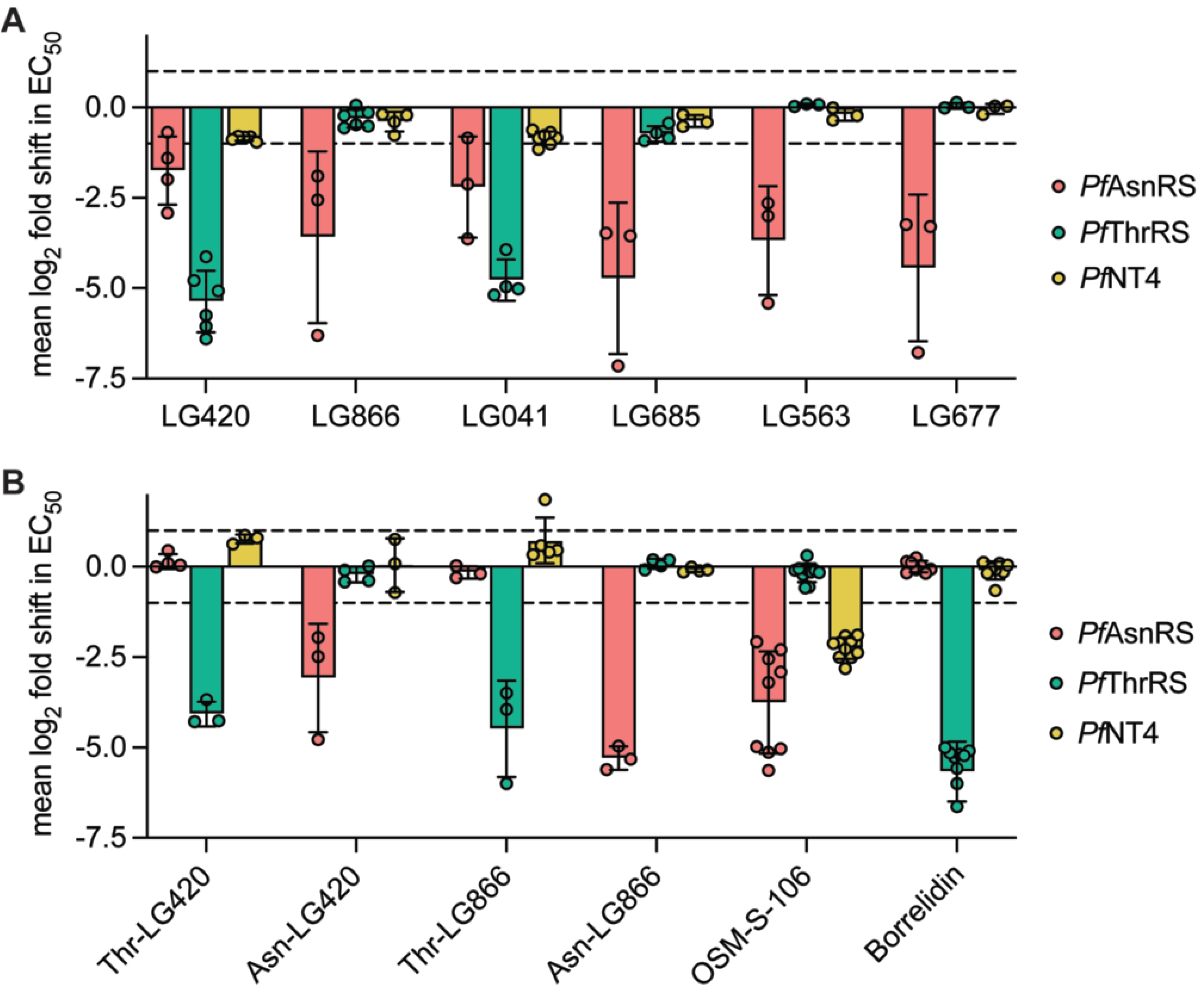
Inducible knock-down analysis confirms *P. falciparum* aaRS targets. Changes in sensitivity to exposure (72-h) to 4AQS series compounds (A) and adducts and controls (B) for aptamer-regulatable *Pf*AsnRS (red), *Pf*ThrRS (green) and *Pf*NT4 (yellow) knock-down lines upon reduction of target protein expression. Bars represent the mean of three to ten replicates and circles represent the values for each replicate. See Supplementary Figure 8 for individual drug inhibition curves.

As controls, we examined the effect of the *Pf*AsnRS and *Pf*ThrRS knockdowns on the potency of synthetic Asn- and Thr- adducts of LG420 and LG866. As anticipated, the potency of the Thr adducts was enhanced in the *Pf*ThrRS knockdown, while the potency of the Asn adducts was enhanced in the *Pf*AsnRS knockdown (Figure 3B, Supplementary Figure 8).

Of interest, previous work showed that down-regulation of *P. falciparum* nucleoside transporter 4 (*Pf*NT4) enhanced sensitivity to OSM-S-106 (11). In agreement with that report (11), knockdown of *Pf*NT4 had no effect on the growth of parasites (Supplementary Figure 7) but was associated with increased susceptibility to OSM-S-106 (Figure 3B). By contrast, down-regulation of *Pf*NT4 had no effect on susceptibility to 4AQS series compounds (Figure 3A, Supplementary Figure 8).

### 4AQS hijackers inhibit ATP consumption by PfAsnRS but not Homo sapiens (Hs)AsnRS

We sought to examine the direct effects of 4AQS compounds on the activity of aaRSs. Expression and characterisation of aminoacylation-competent recombinant *Pf*AsnRS and *Hs*AsnRS in *E. coli* have been described (11), and we used the published methods to generate active enzymes. By contrast, our attempts to construct a competent *Pf*ThrRS aminoacylation system *in vitro* were not successful. Therefore, for this work, we focused on the *in vitro* inhibition of AsnRS.

We have previously used the Kinase GLO system (7, 11) to assess the ability of recombinant aaRSs to consume ATP in the initial phase of the aminoacylation reaction, *i.e.*, via the production and release of AMP. In the absence of tRNA, *Pf*AsnRS and *Hs*AsnRS consume low levels of ATP (Figure 4A,B, left hand bars). Addition of *E. coli* tRNA increases the level of ATP consumption by about 5-fold (Figure 4A,B, second bars), consistent with productive aminoacylation.

**Figure 4.**
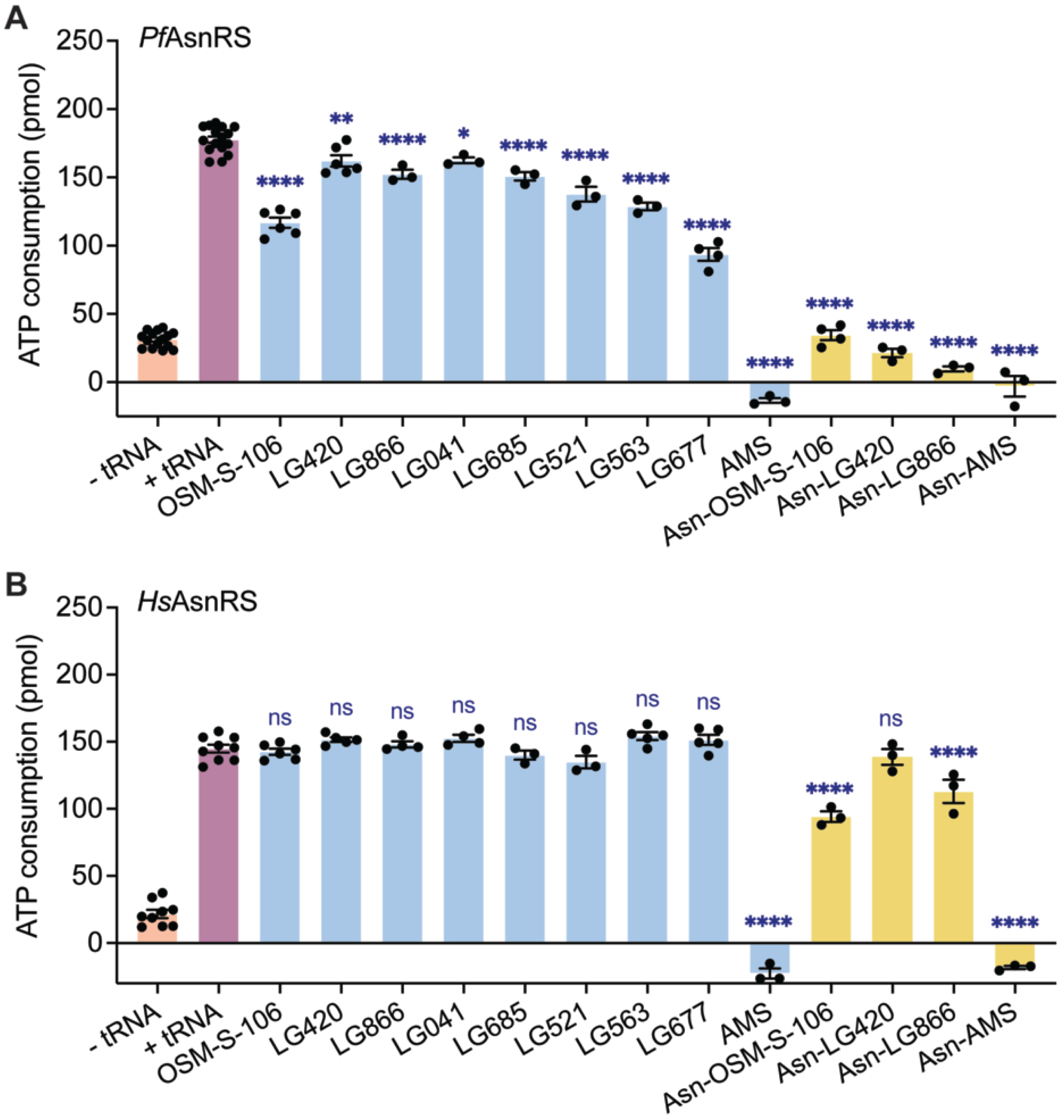
4AQS compounds inhibit ATP consumption by *Pf*AsnRS but not *Hs*AsnRS. Effects of hijacking inhibitors and synthetic amino acid adducts on ATP consumption by *Pf*AsnRS (A) and *Hs*AsnRS (B). Reactions were incubated at 37°C for 1 h. Controls consist of AsnRS in the absence (pink bars) and presence (purple bars) of 100 μM *E. coli* tRNA. Inhibitors were tested at 100 μM (free base compounds, blue bars) or 1 μM (Asn adducts, yellow bars) in the presence of AsnRS and *Ec*tRNA. Data are the average of 3-16 independent experiments. Error bars represent SEM. Asterisks (*) denote the statistical significance of differences in ATP consumption relative to the AsnRS plus tRNA control, as determined by ordinary one-way ANOVA with Dunnett’s multiple comparisons test (GraphPad Prism, version 11.0.0; ns, not significant; *p < 0.05, **p < 0.01, ***p < 0.001, ****p < 0.0001). See Table 2 and Supplementary Table 7 for individual percentage inhibition values.

OSM-S-106 has previously been reported as a hijacking inhibitor of recombinant *Pf*AsnRS, when it is added in the presence of all substrates, *i.e*., Asn, ATP and tRNA (11). Here we confirmed that activity (Figure 4A). Incubation with each of the 4AQS series results in a significant decrease in ATP consumption by *Pf*AsnRS (Figure 4A), with the level of inhibition increasing with the potency of the compounds in the 3D7 growth assay (Table 1; Supplementary Figure 9A). For example, LG677, the most potent anti-plasmodial compound, is also the most active inhibitor *in vitro* (Figure 4A). The synthetic adducts, Asn-OSM-S-106, Asn-LG420 and Asn-LG866 are also very efficient inhibitors of ATP consumption (Figure 4A).

The 4AQS compounds show no activity as hijacking inhibitors of *Hs*AsnRS ATP consumption in this *in vitro* assay (Figure 4B), consistent with our previous result for OSM-S-106 (11). These data suggest that *Hs*AsnRS is intrinsically less susceptible to hijacking by this class of sulfonamides. By contrast, adenosine 5’-sulfamate (AMS), a broadly reactive and highly potent hijacker (7), inhibits ATP consumption by both *Pf*AsnRS and *Hs*AsnRS (Figure 4A,B). Of interest, the synthetic adducts, Asn-OSM-S-106, Asn-LG420 and Asn-LG866, are also less efficient inhibitors of ATP consumption by recombinant *Hs*AsnRS (Figure 4B; Supplementary Table 7).

### 4AQS hijackers induce thermal stabilisation of PfAsnRS but have no effect on HsAsnRS

Differential Scanning Fluorimetry (DSF) has previously been used to monitor hijacking of *P. falciparum* TyrRS (7, 8). Interestingly, apo *Pf*AsnRS is highly stable, exhibiting a melting temperature (T_m_) of 60°C (Supplementary Table 8; Supplementary Figure 9B). By contrast, recombinant apo *Hs*AsnRS exhibits a T_m_ of 51.5°C (Supplementary Table 9). We found that the high basal stability of *Pf*AsnRS made it difficult to use DSF to monitor subtle increases in T_m_ upon hijacking reactions. However, in work-up experiments, we found that inclusion of a low concentration of a chaotropic agent (0.001% SDS) during the final thermal melting step enhanced the ability to discriminate changes induced by ligand binding. Under these conditions, incubation with hijackers stabilised *Pf*AsnRS to an extent that, while modest, correlates the increase in potency (Figure 5A, Table 2). For comparison, the known high potency hijacker, AMS, increased the T_m_ by 4.5°C (Figure 5A, Supplementary Table 9). Similarly, pre-formed synthetic Asn-hijacker adducts caused substantial stabilisation (Figure 5B, Supplementary Table 9).

**Figure 5.**
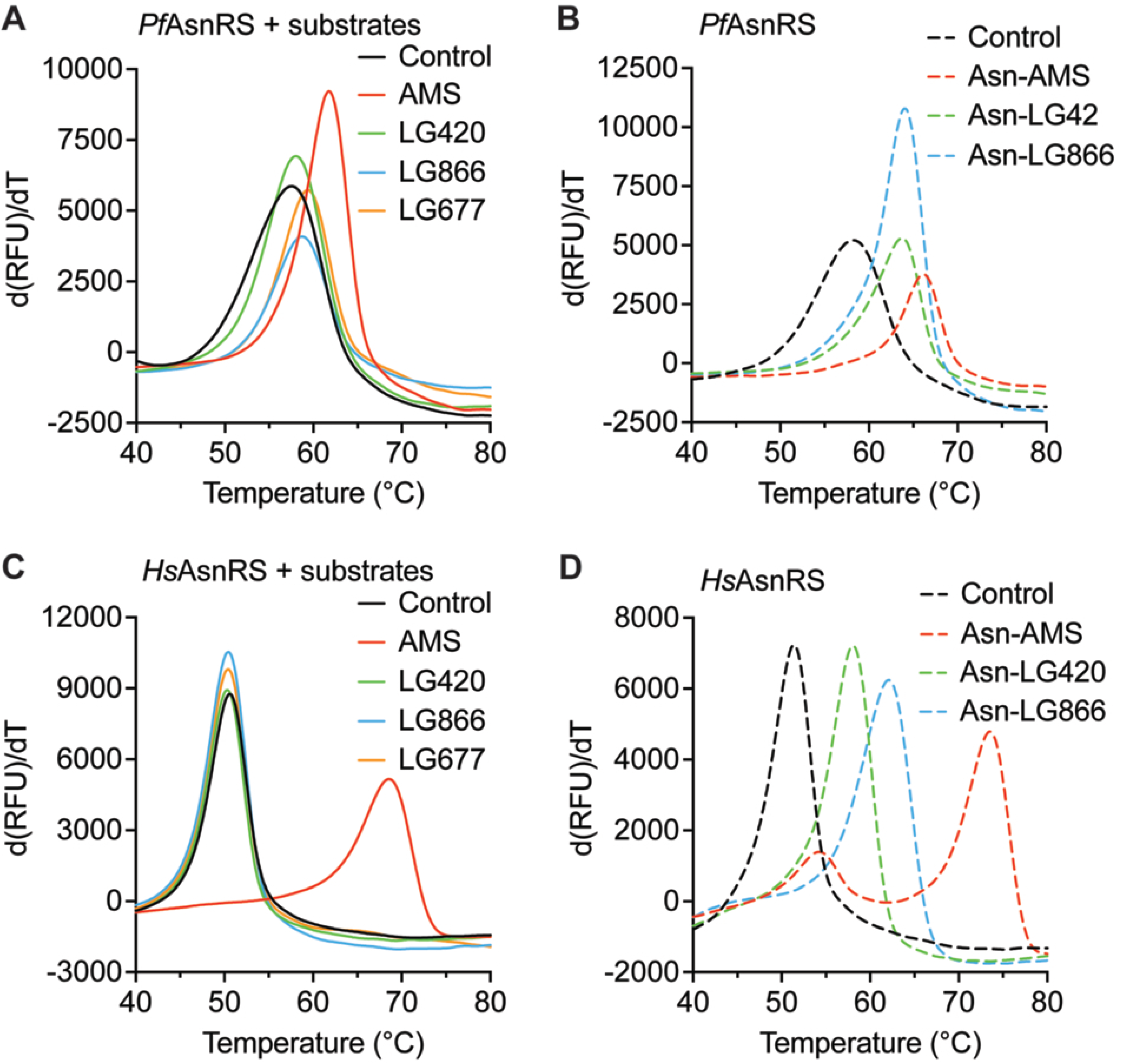
Thermal stabilization of *Pf*AsnRS and *Hs*AsnRS by 4AQS compounds and adducts. (A-D) First derivatives of melting curves for *Pf*AsnRS (A,B) and *Hs*AsnRS (C,D) (1.5 μM) after incubation at 37°C for 3 h with 100 μM LG420, LG866, LG677 and AMS in the presence of 20 μM ATP, 40 μM Asn, 80 μM *Ec*tRNA (A, C) or 100 μM Asn adducts of LG420, LG866 and AMS (B, D). Controls consist of AsnRS in the presence of ATP, Asn and *Ec*tRNA (A, C), or AsnRS alone (B, D). For *Pf*AsnRS samples, the mixture was destabilised by the addition of 0.001% SDS, prior to the thermal melting step. Data are representative of 3-7 independent experiments. See Supplementary Table 9 for Tm values for all 4AQS compounds examined.

Recombinant human AsnRS showed no stabilisation upon incubation with any of the 4AQS compounds (Figure 5C). It is difficult to make a direct comparison given the difference in protocol for the two enzymes. Nonetheless, the data are consistent with the suggestion that *Hs*AsnRS is intrinsically less susceptible to hijacking by 4AQS. By contrast, *Hs*AsnRS is susceptible to hijacking by AMS, which led to a substantive (17.8°C) increase in the T_m_ (Figure 5C, Supplementary Table 9). *Hs*AsnRS is also highly stabilised upon incubation with pre-formed synthetic Asn-hijacker adducts (Figure 5D, Supplementary Table 9).

### PfAsnRS models provide insights into hijacker pose and the resistance-conferring mutation

The susceptibility of *Pf*AsnRS to reaction hijacking depends on its ability to generate the Asn-4AQS adduct, which in turn depends on positioning the sulfonamide in a suitable orientation to attack the relevant carbonyl carbon of Asn-tRNA. For this work, we built on a previously generated model of *Pf*AsnRS and included a conserved structural water molecule seen in experimental aaRS structures mediating interactions between the backbone of active site residues (in *Pf*AsnRS Glu583 and Arg584) and the nitrogen at position 3 of the AMP purine. Here the water molecule provides the same binding interaction with the quinazoline nitrogen of the hijacking ligand (Figure 6A). *Pf*AsnRS exhibits typical characteristics of a Type II aaRS, comprising an N-terminal β-barrel anticodon-binding domain connected via a hinge region to a larger C-terminal catalytic domain that adopts an α-β fold (Figure 6A). Three motifs (I-III) that are involved in ATP binding and dimerization (23) are highlighted (Figure 6A). An alignment of sequences of AsnRSs from different species, showing the conservation of these motifs is presented in Supplementary Figure 10.

**Figure 6.**
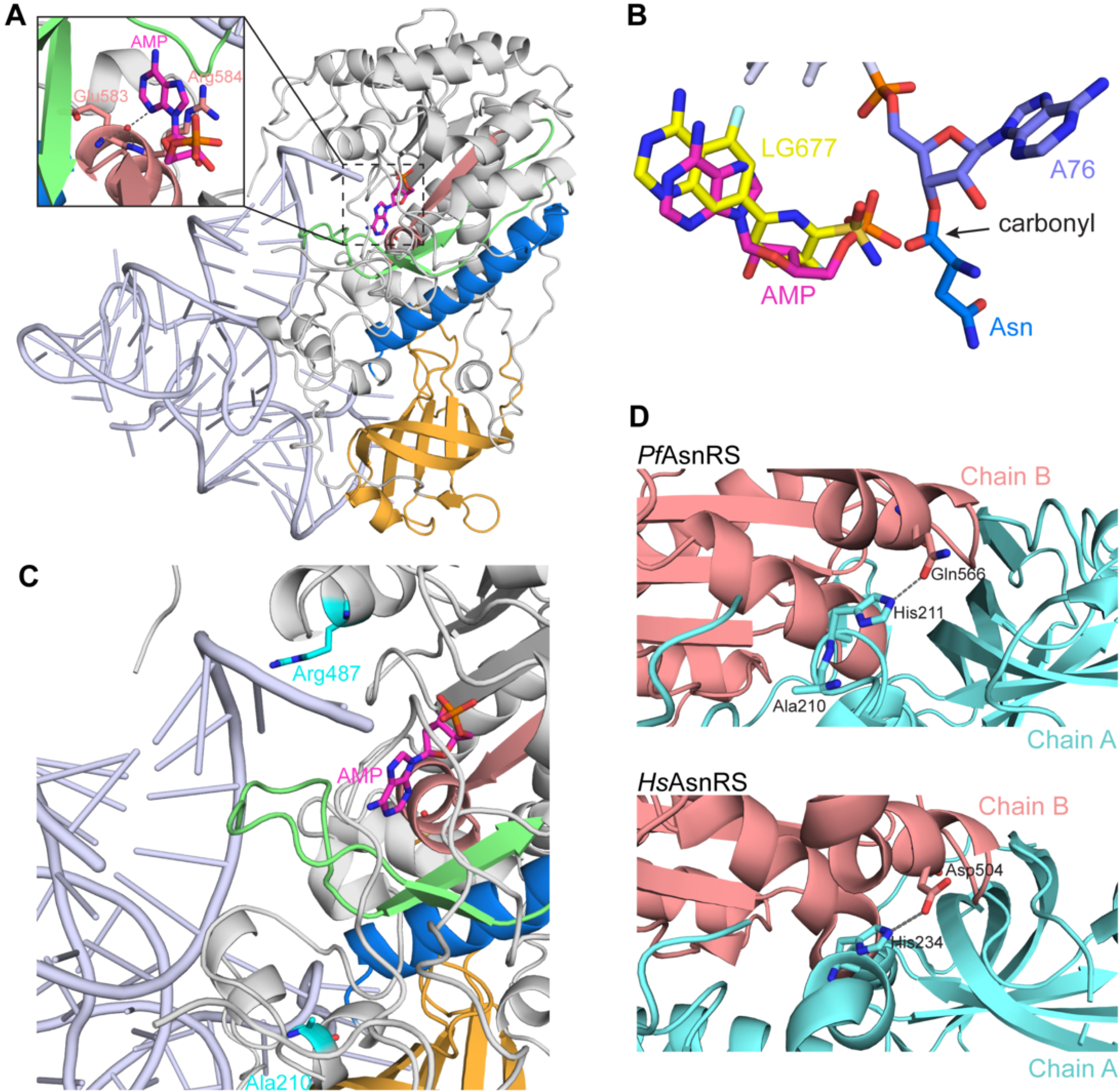
A *Pf*AsnRS-Asn-tRNA complex and *Pf*AsnRS dimer models. (A) Ribbon diagram representation of *Pf*AsnRS in complex with Asn-tRNA and AMP, generated by overlay of an AlphaFold model of *Pf*AsnRS with the *E. coli* AspRS/tRNA(Asp) complex (PDB ID 1C0A) (59). The anticodon-binding domain is shown in yellow, motif I in blue, motif II in green and motif III in pink. The position of the bound ligand (magenta) is highlighted with a dotted black square. Inset shows a zoom of the active site, with a solvent water (red sphere) that likely mediates interactions between the active site residues (Glu583 and Arg584), and the ligand. (B) Close-up view of the active site with 3’ terminal A76 of the tRNA and representative in silico docks of AMP and LG677 to the *Pf*AsnRS-Asn-tRNA model. The sulfonamide is correctly positioned to mount an attack on the Asn-tRNA carbonyl carbon. (C) The *Pf*AsnRS-Asn-tRNA model showing that Ala210 is distal to the active site and the Arg487Ser mutation site. (D) Upper panel. Close-up view of the dimer interface of an AlphaFold Multimer model of *Pf*AsnRS, showing Ala210 and the adjacent His211 which interacts with Gln566 in the other monomer. Lower panel. Close-up view of the dimer interface of the *Hs*AsnRS dimer (PDB ID 8TC7) showing the equivalent His234 which interacts with Asp504 in the other monomer.

The most potent compound, LG677, the natural ligand AMP, and the known *Pf*AsnRS hijacker, OSM-S-106, were docked into the catalytic site, in the context of the bound Asn-tRNA product, using Surflex in SybylX2.1. AMP docks into the site as anticipated with the phosphate positioned close to the amino acid tRNA ester bond and the ribose twisted relative to the adenine (Figure 6B). The quinazoline moiety of LG677 docks into the adenine-binding site and appears to be stabilised by a cation-π interaction with the guanidine moiety of Arg584 (Supplementary Figure 11A). Interestingly, the pyridine ring, which is located in the ribose-binding site, is twisted toward planarity with the quinazoline ring (Figure 6B). This co-planarity means that the sulfonamide nitrogen of LG677 overlaps with the AMP phosphate despite the shorter overall length of LG677. Similarly, the amino groups on the quinazoline ring of LG677 and the adenine of AMP occupy similar positions. Docked OSM-S-106 adopts a very similar pose to LG677 (Supplementary Figure 11B), consistent with our observation that these molecules both preferentially target *Pf*AsnRS. The fluorine on the pyridine moiety of LG677 faces a hydrophobic environment lined by Val534, Ile535 and Leu580 (Supplementary Figure 11A).

As described above, pulsed drug selection of a more highly mutating parasite line led to recrudescent lines in which *Pf*AsnRS harbours a Ala210Thr mutation. Examination of the Asn-tRNA-bound *Pf*AsnRS shows that Ala210 mutation lies well away from the catalytic site and the tRNA binding regions (Figure 6C). It is also distal to the Arg487Ser mutation that is associated with low level resistance to OSM-S-106 and LG866. *Pf*AsnRS forms a native dimer (11). We generated a model of the dimer using AlphaFold Multimer (24, 25). Interestingly, Ala210 sits close to the dimer interface, directly adjacent to a highly conserved histidine residue (His211; Supplementary Figure 10). His211 immediately precedes motif I, which mediates dimerisation of class II aaRS homodimers (26) (Figure 6D, top panel). In *Hs*AsnRS, the equivalent residue, His234, also contributes to stabilisation of the dimer interface (Figure 6D, bottom panel).

## Discussion

The genomes of many organisms encode multiple genes for multiple tRNA isoacceptors, *i.e.,* tRNAs that are charged with the same amino acid but have different anticodons (27). By contrast, *Plasmodium* encodes a minimal set of 45 tRNA genes in its nuclear genome, covering both cytoplasmic and apicoplast tRNAs, with only one gene copy per tRNA isoacceptor (28). This makes *P. falciparum* particularly susceptible to aaRS inhibitors (29). Recently, AMP-mimicking sulfonamide and sulfamates have been shown to act as reaction-hijacking inhibitors of either class I or class II aaRSs, attacking the enzyme-bound charged tRNA and forming an inhibitory adduct with the amino acid. For example, ML471 targets *Pf*TyrRS with high potency and selectivity, both *in vitro* and *in vivo* (7, 8); however, the ML471 scaffold exhibits low lipophilicity, which limits its oral bioavailability (10). Moreover, this class of hijacker showed susceptibility to the development of resistance. Another hijacker, OSM-S-106, targets *Pf*AsnRS, plus additional minor aaRS targets (11). OSM-S-106 is refractory to resistance development, and has improved physicochemical characteristics, but exhibits lower potency than ML471. We therefore sought an additional compound class with hijacking capability.

While reaction-hijacking AMP-mimicking sulfonamides/ sulfamates have only recently been studied as “pro-inhibitors” that lead to the generation of tight-binding adducts *in situ*, “dual site” sulfomyl AMP mimic-amino acid adducts have long been explored as inhibitors of aaRSs and as stabilisers for structural studies of aaRSs (30–32). These adducts can exhibit potent activity in biochemical assays; however, they generally lack species-selectivity and are expected to be difficult to develop as anti-infectives due to poor cellular permeability and oral bioavailability (33–36).

Nonetheless, the potencies of amino acid conjugates of sulfamoyl adenylates can provide inspiration for the development of new hijacking precursors. For this work, we built on work examining amino acid adducts of a series of AMP mimicking benzenesulfonamides, which were explored as dual site inhibitors of bacterial aaRSs. Teng *et al.* explored threonine (Thr) conjugated benzenesulfonamides as inhibitors of ThrRS (36), demonstrating that, at least in the context of the amino acid adduct, the adenylate phosphate of AMP can be substituted with a benzenesulfonamide group, while a phenyl group can be used to occupy the ribose binding site (36). Similarly, proline conjugates of a 4-amino quinazoline benzene sulfonamide series have been explored as inhibitors of bacterial prolyl-tRNA synthetase (ProRS). This study used a fluorine scanning strategy and demonstrated that fluorine substitutions at specific positions selectively enhanced activity against *Pseudomonas aeruginosa* ProRS. (12). Leucine conjugates of benzenesulfonamide-based AMP mimics have also been explored as inhibitors of bacterial and trypanosomal LeuRS (37, 38).

Compound **10a** from the Teng *et al.* series (36) is a Thr adduct of an AMP-mimicking 4-aminoquinazoline benzene sulfonamide core. The “pro-inhibitor” core of **10a** represents a direct swap of the aminothienopyrimidine core of OSM-S-106 (11), with a 4-aminoquinazoline core. We sought to determine whether the 4-aminoquinazoline form of OSM-S-106 retained hijacking activity against susceptible *P. falciparum* aaRSs.

The first compound synthesised, LG420, showed moderate activity against *P. falciparum* cultures (IC_50_ = 265 nM). Interestingly, targeted mass spectrometry revealed the formation of apparently high levels of Thr-LG420 and Asn-LG420. Similarly, analysis of *Pf*ThrRS and *Pf*AsnRS conditional knock-down lines confirmed that the potency of LG420 is enhanced when the abundance of either of these targets is decreased, indicating that LG420 is a bona fide dual targeting hijacker. Both *Pf*AsnRS (39) and *Pf*ThrRS have been previously validated as targets in *P. falciparum* (21, 40).

To improve the potency of the series, the LG420 phenyl sulfonamide was modified to the equivalent pyridine sulfonamide, with the N in the ortho (2’) position (LG866). This modification is designed to enhance the nucleophilicity of the usually poorly reactive sulfonamide group, so as to promote attack on the ester bond of the charged tRNA. As anticipated, this change gave a 7-fold increase in potency. Of interest, targeted mass spectrometry and conditional knockdown studies revealed an apparent switch in enzyme susceptibility, such that *Pf*AsnRS is now the major target, with *Pf*ThrRS, *Pf*LysRS and *Pf*AlaRS and *Pf*GlyRS as more minor targets. This result suggests that efficient targeting of *Pf*AsnRS is associated with potent antimalarial activity. LG420 and LG866 were shown to activate the amino acid starvation response, further confirming involvement of aminoacyl tRNA synthetase targets.

Fluorine substituents exhibit electronegativity which can alter the electronics of aromatic rings and enhance hydrogen bonding interaction with residues in the surrounding pocket (41, 42). Fluorine also exhibits lipophilicity which can enhance permeability and cellular activity (43). Recently Luo *et al.* successfully used fluorination of the phenyl and 4-aminoquinazoline groups of Pro adducts to enhance activity against *Pseudomonas aeruginosa* ProRS (12). A crystal structure of *Pa*ProRS with the optimised inhibitor, PAA-38, revealed that fluorine substitutions at the C8 position of the quinazoline and the C6’ position of the phenyl group were oriented toward hydrophobic environments and contributed new hydrophobic and van der Waals interactions (12). We therefore explored fluorination of the phenyl ring of LG420 and the pyrimidine of LG866 in the 6’ position (LG041 and LG685), to explore potential fluorophilic pockets and, additionally, to increase the nucleophilicity of the sulfonamide warhead. These substitutions provided an additional 3-fold increase in potency. Again, targeted mass spectrometry and conditional knockdown studies revealed *Pf*AsnRS as the major target, with similar secondary targets to LG866.

Fluorination of the 4-aminoquinazoline ring in the 5- or 8-position (LG877 and LG521), gave further potency increases, and a combination of these modifications produced compounds LG563 and LG677, with low nanomolar activity against *P. falciparum* cultures. Again, the main *P. falciparum* aaRS is *Pf*AsnRS, with other minor targets identified by targeted mass spectrometry. Upon conditional knockdown of *Pf*AsnRS, the IC_50_ value for inhibition of parasite cultures by LG677 was 0.22 nM, confirming the target and indicating the marked potency of this compound.

The switch of target towards *Pf*AsnRS for more potent 4AQS compounds may reflect the fact that this aaRS is particularly susceptible to hijacking. For example, the broadly active hijacker dealanylascamycin (DACM) produces the Asn adduct as the most abundant product (44). In addition, a single tRNA(Asn) is responsible for all the Asn-tRNA(Asn) needed to produce a proteome rich in asparagine repeats. Indeed, the ratio of the tRNA(Asn) gene copy number to amino acid frequency is lowest of the *P. falciparum* tRNAs, potentially limiting translation efficiency (39). This may make *P. falciparum* AsnRS a particularly susceptible target.

One important characteristic of new antimalarial candidates is that they should exhibit a low propensity for resistance. We found that no resistant parasites emerged from an inoculum of 10^9^ exposed parasites, which compares well with other compounds selected for development (45). Similarly, no mutant lines were enriched from a resistome panel, when selected with 4AQS compounds at 3 x IC_50_ value. Nonetheless, we found that the known *Pf*AsnRS^R487S^ mutant provides low level resistance (∼2-fold) to LG866, but not LG420. The dual targets of LG420 and the minor secondary targets of other 4AQS compounds may protect these compounds against the ready development of resistance.

In previous work (11), ramp-up exposure pressure to OSM-S-106 also selected for parasite clones harbouring mutations in *Pf*NT4. *Pf*NT4 is a putative purine transporter (46–48), and it has been suggested that mutations in *Pf*NT4 may enhance the transport of OSM-S-106 away from its site of action. In this work, no mutations in *Pf*NT4 or gene amplification were observed, suggesting that the 4AQS series is not a substrate for this transporter.

The fast mutator line, Dd2-Polδ, harbours DNA polymerase-δ mutations that impair 3ʹ−5ʹ proof-reading activity (16). When a large inoculum of this parasite line was exposed to 3 x IC_90_ of LG866, a mutant line emerged harbouring a *Pf*AsnRS^A210T^ mutation. Ala210 is distal from the active site and tRNA binding interface. The Ala210Thr mutation lies adjacent to residue His211, which is conserved in human, yeast and other eukaryote AsnRSs. In *Hs*AsnRS, the equivalent His interacts with the neighbouring monomer. In our *Pf*AsnRS dimer model, His211 are predicted to be involved in dimer stabilisation. The Ala210Thr mutation introduces a larger polar residue that may alter the conformation of His211 and affect its ability to mediate dimer stability. In class II aaRSs, the tRNA bound to one monomer makes some interactions with the other monomer (49). Thus, the stability of the dimer interface may indirectly affect the longevity of the Asn-tRNA product in the active site, which may in turn affect hijacking susceptibility. The fact that no resistance-conferring mutations were observed involving residues close to the active site suggests that such mutations might affect enzyme function.

While it has not been possible to obtain X-ray crystallographic data for *Pf*AsnRS, our modelling studies reveal that LG677 adopts a pose in the active site that positions the sulfonamide ready for attack on the carbonyl carbon of the charged tRNA. Moreover, our analysis reveals that the fluorine substituent on the pyrimidine ring of LG677 appears to access a hydrophobic environment within the active site pocket of *Pf*AsnRS, which may contribute to improved potency.

We generated recombinant *Pf*AsnRS and showed that 4AQS compounds inhibit enzyme activity with a SAR that correlates with cellular activity. We note a discrepancy between the cellular potency (nM range) and the *in vitro* enzyme inhibition (μM range). This difference is likely due to the fact that, in the biochemical assay, the enzyme first needs to generate the charged tRNA product, which is then attacked by the pro-inhibitor to generate the Asn-4AQS adduct. Several rounds of ATP consumption and aminoacylation may be needed before the hijacker-amino acid adduct is formed. In the cellular assay, growth inhibition is monitored over a period of 72 h, which is likely sufficient time for the formation of adducts, and thus loss of aaRS function.

Similarly, the 4AQS compounds stabilise *Pf*AsnRS against thermal denaturation. Importantly, under the conditions of our *in vitro* assays, there is no evidence for stabilisation of human AsnRS, or for inhibition of human AsnRS enzyme activity. Accordingly, the series exhibited up to 300-fold selectivity compared with the mammalian cell line, HepG2. Nonetheless, further improvements in selectivity would be needed for further development of this compound series.

In summary, the 4AQS series of reaction hijacking inhibitors reported here offer very potent activity against blood stage *P. falciparum* and reasonable activity against transmissible stages. The ability to target multiple aaRSs appears to reduce the susceptibility to development of resistance, a critical feature for next-generation antimalarials. Thus, the 4AQS series offers an interesting starting point for the development of new antimalarial compounds to combat this lethal pathogen.

## Methods

### Activity against P. falciparum cultures

Analysis of activity against *P. falciparum* laboratory strains, 3D7 and Dd2, was tested by TCG Lifesciences, Kolkata, India using a *P. falciparum* lactate dehydrogenase growth inhibition assay as described previously (50). Following exposure of cultures, an aliquot of freshly prepared reaction mix (70 μL) containing 100 mM Tris-HCl pH 8, 143 mM sodium L-lactate, 143 μM 3-acetyl pyridine adenine dinucleotide, 179 μM Nitro Blue tetrazolium chloride, diaphorase (2.8U/mL) and 0.7% Tween 20 was added to each well of the assay plate. After mixing, plates were placed at 21⁰C for 20 min, in the dark. The percentage of growth inhibition was normalized relative to positive (0.2% DMSO, 0% inhibition) and negative (mixture of 100 μM chloroquine and 100 μM atovaquone, 100% inhibition) controls. *P. falciparum* strains (3D7 and Dd2) were obtained from BEI Resources. Three independent experiments were performed for each assay, each in technical duplicate.

### Antimalarial resistome barcode sequencing (AReBar) cross-resistance assay and activity against a PfAsnRS^R487S^ transfectant cell line

Susceptibility of a pool of 53 barcoded lines (Supplementary Table 5), covering 33 modes of antimalarial action, was assessed using the AReBar assay, as described previously (17). Each uniquely barcoded line possesses distinct resistance mutations in either drug targets or resistance genes, predominantly in the Dd2 background. Compounds were applied to the pool (1 mL × triplicate cultures) at 3 × IC_50_ over the 14-day assay period. Parasitemia was monitored by flow cytometry (CytoFlex 5, Beckman) every 2–3 days, with cells stained with 1 × SYBR Green I, 0.2 µM Mitotracker Deep Red, and parasitemia maintained between 0.3%–5% parasitemia. Samples were harvested at days 0 and 14, lysed with 0.05% saponin, washed twice with PBS, and resuspended in 30 µL PBS prior to barcode amplification. Sequencing of barcode amplicons was performed using a minION Mk1C (Oxford Nanopore), and read counts were analyzed using DESeq2 (51) to quantify differentially represented barcodes relative to the no-drug control. Significance testing was performed by Wald test. CRISPR/Cas9 editing was previously used to generate parasites encoding the R487S mutation in *Pf*AsnRS in a Dd2 parasite background (11). Dose response assays were performed for 72 h, with compounds plated using a Tecan D300e dispenser. Parasites were added at 1% parasitemia, 1% hematocrit. After 72 h, assays were lysed and stained with 1×SYBR Green, incubated >1 h, and measured using a FLUOstar Omega plate reader.

### Activity against gametocyte stages of P. falciparum

Gametocytogenesis was induced on a tightly synchronised (>97% rings) asexual parasite culture (Pf3D7*elo1*-*pfs16*-CBG9; kind gift from Pietro Alano), 0.5% parasitemia and 6% hematocrit) with a combination of nutrient starvation and a decrease in hematocrit, as previously described (52). For immature gametocytes (>90% stage II/III), cultures were exposed to 50 mM N-acetylglucosamine (NAG) on days 1–4 to eliminate residual asexual parasites and harvested on days 5–6. For mature (>95% stage V) gametocytes, NAG treatment occurred on days 3–7, and the parasites were harvested at day 13. Immature and mature gametocyte cultures (2% gametocytaemia 1.5 % haematocrit, 150 μL/well) were exposed to compounds and incubated stationary at 37°C for 48 h under hypoxic conditions (53), after which luciferase activity was determined with a non-lysing D-luciferin substrate (1 mM in 0.1 M citrate buffer, pH 5.5) and bioluminescence was detected with a 2 s integration time with a GloMax^®^ Explorer Multimode Microplate Reader (Promega).

### Activity against P. falciparum gametes

Inhibition of activation of stage V male and female gametocytes to form gametes was assessed, broadly as described previously (54). Briefly, *P. falciparum* NF54 strain stage V gametocytes were incubated in 384 well plates with test compounds at 37°C for 48 hr. Then, gametogenesis was stimulated by decreasing plate temperature and addition of xanthurenic acid. Twenty minutes later, the emergence of male gametes was recorded microscopically using phase contrast and ×4 objective lens using automated microscopy. The plate was incubated at 26°C for a further 24 h to allow emerged female gametes to maximally express *Pf*s25 on their cell surface. An anti-Pfs25 antibody (Mab 4B7) conjugated to the Cy3 fluorophore was added to the plates and female gametes detected by live fluorescence microscopy at ×4 objective. Data was processed using custom image analysis algorithms and percent inhibition relative to positive Cabamiquine (DDD498) and negative (DMSO) controls was calculated. Data values are the mean of independent biological repeats.

### Activity against HepG2

Viability of the HepG2 (Human Caucasian hepatocyte carcinoma) cell line from ATCC (American Type Culture Collection, Manassas, USA; HB-8065) was assessed using the Cell Titer-Glo luminescence cell viability kit (Promega). For the assay, 2,000 cells/well were plated in 384-well plates 24 h prior to the experiment and incubated in a CO_2_ incubator at 37°C. Following removal of the media, cells were treated with fresh medium containing either vehicle (0.5% DMSO) or serially diluted compounds or doxorubicin (1.3 nM to 25 μM) in a final volume of 50 μL/well and further incubated for 72 h at 37°C in a CO_2_ incubator. Positive control wells (100% inhibition) were treated with 5 μL of 1% Triton X-100. Following incubation, 25 μL of medium was discarded and 25 μl of CellTiter-Glo reagent was added to each well and the plate was kept on a plate shaker for 15 min at 25°C with shaking at 300 rpm. Luminescence signals were measured in a PHERAstar FSX reader (BMG LABTECH). IC_50_ values were calculated using the 4-parametric logistic curve fitting program of Graphpad prism (version 5). Three independent experiments were performed for each assay, each in technical duplicate.

### Minimum inoculum of resistance

Minimum Inoculum of Resistance (MIR) studies were conducted for LG866 as described previously (45). The in-house mean IC_50_ of LG866 against Dd2-B2 and Dd2-Polδ was 48 nM, with a corresponding mean IC_90_ value of 95 nM (N, n = 2, 2). A total inoculum of 10^9^ parasites in three flasks were treated with 3 x IC_90_ of compound and cleared from the culture rapidly. Wells were monitored daily by smear during the first seven days to ensure parasite clearance, during which media was changed daily. Thereafter, cultures were screened three times weekly by flow cytometry and smearing, and the selection maintained under consistent drug pressure over 60 days. Giemsa-stained blood smears examined by light microscope showed no parasite recrudescence in the Dd2-B2 line, while parasites were observed in both Dd2-Polδ flasks on day 42.

### Whole-genome sequencing and analysis of LG866-resistant parasites

Select recrudescent wells were scaled up to at least 4-5% parasitemia at 3% hematocrit before parasite extraction from the red blood cells (RBCs). Briefly, infected RBCs were washed in phosphate buffered saline (PBS, pH 7.4) and the pellet lysed in 0.15% saponin in PBS for 5 min at 37°C followed by high-speed centrifugation. Pellets were washed twice with PBS and stored at −80°C until genomic DNA was isolated using a Blood DNA Mini Extraction Kit and sent for whole genome sequencing (WGS). Whole-genome sequencing was performed using a Nextera Flex DNA library kit and multiplexed on a MiSeq flow cell to generate 300 bp paired-end reads. Sequences were aligned to the *P. falciparum* 3D7 reference genome (PlasmoDB-48_Pfalciparum3D7) using the Burrow-Wheeler Alignment (BWA version 0.7.17). PCR duplicates and unmapped reads were filtered out using Samtools (version 1.13) and Picard MarkDuplicates (GATK version 4.2.2). Base quality scores were recalibrated using GATK BaseRecalibrator (GATK version 4.2.2). GATK HaplotypeCaller (GATK version 4.2.2) was used to identify all possible single nucleotide variants (SNVs) in test parasite lines filtered based on quality scores (variant quality as function of depth QD > 1.5, mapping quality > 40, min base quality score > 20, read depth > 10) to obtain high-quality single nucleotide polymorphisms (SNPs) that were annotated using SnpEff version 4.3t. BIC-Seq version 1.1.2 was used to test for copy number variants (CNVs) against the parental line using the Bayesian statistical model. SNPs and CNVs were visually inspected and confirmed using Integrative Genome Viewer (IGV). All gene annotations in the analysis were based on PlasmoDB-48_Pfalciparum3D7. The list of variants from the resistant clones were compared against the Dd2-B2 parent to obtain homozygous SNPs present exclusively in the resistant clones.

### Generation of conditional knockdown parasite lines

Conditional knockdown (cKD) *P. falciparum* lines for cytosolic AsnRS (PF3D7_0211800) and PfNT4 have been described previously (11). To generate a cKD line for ThrRS (PF3D7_1126000), the ThrRS native locus was modified using CRISPR/Cas9 gene editing at the 3’ end to introduce a V5-2x-HA epitope tag, an aptamer array, and the machinery required for TetR-DOZI-mediated translational repression (22, 55). To create the donor plasmid for homology-directed repair, left and right homology regions (LHR and RHR) were amplified from parasite gDNA using PCR, then cloned into the pSN054_V5 plasmid backbone, along with a 100 bp segment prepared using a Klenow reaction containing the sgRNA sequence, and a synthesized DNA fragment that recodonizes the sgRNA recognition site. The pSN054_V5 plasmid also integrates a *Renilla luciferase* reporter and the selection marker *blasticidin-S-deaminase*. The final plasmid was validated using restriction enzyme digestion and Sanger sequencing, then transfected into Cas9- and T7-expressing NF54 parasites by preloading of RBCs (56). Parasite cultures were maintained in 500 nM anhydrotetracyline (aTc, Sigma-Aldrich); transfectants were selected using 2.5 µg/ml blasticidin (RPI Corp) starting 4 days post-transfection. Recovery of parasites was monitored using the Renilla-Glo(R) Luciferase Assay System (Promega) and Giemsa staining (Sigma-Aldrich). The primers used for donor plasmid assembly and the sequences of synthesized DNA fragments are in Supplementary Table 6.

### Growth assay for knockdown parasite lines

To determine the effect of protein knockdown on parasite viability, synchronized ring stage parasites were arrayed in triplicate in U-bottom 96-well plates (BD Falcon) in either 0 nM or 500 nM aTc. Samples were taken at 0 h and 72 h, and parasite biomass quantified using the Renilla-Glo(R) Luciferase Assay System (Promega) using a GloMax® Discover Multimode Microplate Reader (Promega). Growth in the knockdown (0 nM aTc) condition was normalized to luminescence values from cells grown in 500 nM aTc (100% growth) and exposed to 500 nM dihydroartemisinin (DHA) (0% growth). Data analysis and visualization was performed using Prism (GraphPad).

### Susceptibility assays for knockdown parasite lines

To determine the effect of protein knockdown on parasite susceptibility to 4AQS compounds, synchronized parasite lines were arrayed in duplicate at 0.5% parasitemia in the presence (500 nM) and absence of aTc in flat-bottom 384 well plates (Corning) and exposed to a 2-fold dilution series of each compound. After growth at 37°C under 90% N_2_/ 5% CO_2_/ 5% O_2_ atmosphere for 72 h hours, parasite abundance was quantified using luciferase measurements as described above. Luminescence values were normalized to those from parasites exposed to DMSO (100% viability) and 500 nM DHA (0% viability). Dose response curves were fit and estimates for EC_50_ were determined using the R package drc (57).

### Western blotting analysis of eIF2α phosphorylation

Highly synchronous *P. falciparum* 3D7 infected RBCs (30-35 h post-invasion; 2.5% hematocrit, 5-6% parasitemia) were exposed to 4AQS compounds, using borrelidin (Sigma B3061) as a positive control and 0.05% DMSO as a negative control, for 3 h. Infected RBCs were pelleted, washed with ice-cold 1 x PBS + cOmplete™ EDTA-free protease inhibitor cocktail (Roche), and lysed with 0.05% saponin in PBS on ice for 5 min. Washed pellets were resuspended in Bolt LDS sample buffer containing reducing agent and analysed by Western blotting as previously described (58). Membranes were washed three times in PBS + 0.1% Tween 20. Chemiluminescence signal was detected using the Bio-Rad ChemiDoc MP imaging system. Primary antibodies: rabbit anti-phospho-eIF2α (Cell Signaling Technology-119A11; 1:1,000); polyclonal mouse anti-*Pf*BiP, generated using recombinant *Pf*BiP at the WEHI Antibody Services (1:1,000). Secondary antibodies: goat anti-rabbit IgG-peroxidase (Chemicon-A132P; 1:20,000), goat anti-mouse IgG-peroxidase (Chemicon-A181P; 1:50,000).

### Mass spectrometry to identify and quantify the 4AQS-Asn and 4AQS-Thr adducts

Aliquots of late trophozoite stage *P. falciparum* (3D7 strain) culture were exposed to 10 μM 4AQS compounds for 3 h. Following treatment, parasite-infected RBCs were lysed with 0.1% saponin in PBS and the parasite pellet was washed 3 times with ice-cold PBS. Cell pellets were kept on ice and resuspended in water as one volume, followed by the addition of five volumes of cold chloroform-methanol (2:1 [vol/vol]) solution. Samples were incubated on ice for 5 min, subjected to vortex mixing for 1 min and centrifuged at 14,000 x g for 10 min at 4°C to form 2 phases. The top aqueous layer was transferred to a new tube and subjected to LC-MS analysis. Data analysis was performed using Xcalibur (version 4.4).

### High-performance liquid chromatography (HPLC) and mass spectrometric (MS) analyses

Samples were analysed by reversed-phase ultra-high performance liquid chromatography (UHPLC) coupled to tandem mass spectrometry (MS/MS) employing a Vanquish UHPLC linked to an Orbitrap Fusion Lumos mass spectrometer (Thermo Fisher Scientific, San Jose, CA, USA) operated in positive ion mode. Solvent A was 0.1% formic acid in water and solvent B was 0.1% formic acid in acetonitrile. 10 μL of each sample was injected onto an RRHD Eclipse Plus C18 column (2.1 × 1000 mm, 1.8 μm; Agilent Technologies, USA) at 50°C at a flow rate of 350 μL/min. Compounds were separated by gradient elution over 18 minutes with a solvent timetable as follows [time (min), %B]: [0, 0], [6, 0], [13, 25], [13.1, 99], [14, 99], [14.1, 0], [18, 0].

MS experiments were performed using a Heated Electrospray Ionization (HESI) source. The spray voltage, flow rate of sheath, auxiliary and sweep gases were 3.5kV, 25, 5, and 0 ‘arbitrary’ unit(s), respectively. The ion transfer tube and vaporizer temperatures were maintained at 350°C and 150°C, respectively, and the S-Lens RF level was set at 30%. A full-scan mass spectrum and targeted MS/MS spectra for proton adducts of candidate reaction hijacker adducts with 20 possible common amino acids were acquired in cycles throughout the run. The full-scan MS-spectra were acquired in the Orbitrap at a mass resolving power of 120,000 (at *m/z* 200) across an *m/z* range of 200–1500 using quadrupole isolation and the targeted MS/MS were acquired using higher-energy collisional dissociation (HCD)-MS/MS in the Orbitrap at a mass resolving power of 7500 (at *m/z* 200), a normalized collision energy (NCE) of 20% and an *m/z* isolation window of 1.2.

### ATP consumption assay

The consumption of ATP by wildtype *Pf*AsnRS and *Hs*AsnRS was determined using a luciferase-based assay as per the manufacturer’s instructions (Kinase-Glo^®^ Luminescent Kinase Assay, Promega). Reactions were conducted using 25 nM enzyme (*Pf*AsnRS or *Hs*AsnRS) in 50 mM Tris-HCl (pH 7.6), 50 mM KCl, 25 mM MgCl_2_, 0.1 mg/ml BSA, 1 mM TCEP, with 200 μM L-asparagine, 10 μM ATP, 1 unit/mL inorganic pyrophosphatase, and 2.5 mg/mL *E.coli* tRNA (if present). Reactions were incubated with 100 μM free base compound or 1 μM Asn conjugate at 37°C for 1 hour, followed by addition of the Kinase-Glo reagent. Luminescence output was measured using a plate reader (CLARIOstar, BMG LABTECH) and the highest signal within 20 min after addition of reagents was recorded using MARS data analysis software (version 3.32). Assay conditions were optimized to ensure ATP consumption remained in the linear range with respect to AsnRS concentration. The concentration of ATP was quantified by linear regression using an ATP standard curve (Microsoft Excel). Samples with no enzyme served as negative controls. The statistical significance of ATP consumption relative to the AsnRS + *Ec*tRNA positive control was determined by ordinary one-way ANOVA with Dunnett’s multiple comparisons test (GraphPad Prism, version 11.0.0).

### Differential scanning fluorimetry (DSF)

The effect of hijackers compounds and synthetic Asn-hijacker adducts on the thermal stability of AsnRS enzymes was assayed as previously described (Xie et al., 2022), with some modifications. Briefly, 3 μM *Pf*AsnRS or *Hs*AsnRS was incubated with 100 μM compound or Asn-adduct, 40 μM ATP, 80 μM L-asparagine, and 60 μM *Ec*tRNA in 25 mM Tris-HCl (pH 8.0), 150 mM NaCl, 5 mM MgCl_2_, 1 mM TCEP, at 37°C for 3 h. SYPRO Orange (Sigma-Aldrich; 5,000X concentrate in DMSO) was added to the reaction mixture at a final concentration of 10X. 10 μL of the sample was transferred into each well of a 96-well qPCR plate (Applied Biosystems). To facilitate unfolding of the highly stable apo *Pf*AsnRS during the denaturation step, a low concentration of SDS was used as a destabilising agent. The final concentration of SDS (0.001%) was determined empirically by titration. Accordingly, 10 μL of SDS solution (0.002%) was added to each *Pf*AsnRS sample, while 10 μL of 2X Tris buffer was added to each *Hs*AsnRS sample, resulting in a two-fold dilution of each. The plate was sealed and analyzed using StepOnePlus Real-Time PCR system (Applied Biosystems). The samples were heated from 20°C to 90°C with a 1°C per min continuous gradient. The thermal unfolding curve was plotted as the first derivative curve of the raw fluorescence values. The melting temperature (Tm), defined as the peak of the first derivative curve, was used to assess the thermal stability of protein-ligand complexes.

### Modelling of the P. falciparum AsnRS-Asn-tRNA complex

A model of the *Pf*AsnRS-Asn-tRNA complex was generated by combining a modified version of the AlphaFold model for *Pf*AsnRS bound to Asn-AMP with the tRNA from the structure of the *E. coli* aspartyl-tRNA synthase/ tRNA complex, 1C0A, as described previously (11). The structural water molecule incorporated in the revised model was positioned on the basis of HOH1003 in chain A of 1C0A.

## Data Availability

Additional data are available in Supplementary Information. Source data are provided.

## Acknowledgements

OSM-S-106 and Asn-OSM-S-106 were kindly provided by Professor Matthew Todd, University College London, United Kingdom. AMS was kindly provided by Dr Steven Langston, Takeda Pharmaceuticals Inc. Asn-AMS was kindly provided by Derek S. Tan, Sloan Kettering Institute, New York, USA. We thank Elizabeth Winzeler, University of California, San Diego, USA, and Michael Griffin, University of Melbourne, for helpful discussions.

L.T. and S.C.X acknowledge funding from the Australian National Health and Medical Research Council (APP2019492) and the Australian Research Council (DE230101173 to SX; DP250101586 to L.T. and S.C.X). M.C.S.L. and J.N. acknowledge funding from the Gates Foundation (INV-045906, M.C.S.L; INV-026505 and INV-045904, J.N.). LMB acknowledges the Medicines for Malaria Venture (RD-19-0001) for funding. D.A.F. acknowledges funding support from the NIH (R01 AI185559).

## Author Contributions

Conceptualisation: X.Y., L.E., N.K., S.K.N, L.C.G., C.J.M., C-W.T., M.J.D. L-M.B., K.L., G.D., M.C.S.L., D.A.F., J.C.N., M.G.S., L.T., S.C.X.; Investigation: X.Y., L.E., N.K., S.K.N, L.C.G., N.B, C.D., C.J.M., T.R., M.T.F., C-W.T., T.Y., L. H.M.L., M.G.L., M.L.D.S., E.C., M.J.D. L-M.B., J.D., K.L., G.D., D.A.F., J.C.N., M.G.S., L.T., S.C.X.; Analysis: X.Y., L.E., N.K., S.K.N, L.C.G., N.B, C.D., C.J.M., T.R., M.T.F., C-W.T., T.Y., L. H.M.L., M.G.L., M.L.D.S., E.C., M.J.D. L-M.B., J.D., K.L., G.D., M.C.S.L., D.A.F., J.C.N., M.G.S., L.T., S.C.X.; Funding acquisition: M.J.D. L-M.B., K.L., G.D., M.C.S.L., D.A.F., J.C.N., M.S., L.T., S.C.X.; Writing: W.Y., L.E., N.K., S.K.N, L.C.G., N.B, C.D., C.J.M., T.R., C-W.T., T.Y., M.G.L., M.L.D.S., E.C., M.J.D. L-M.B., J.D., M.C.S.L., D.A.F., J.C.N., M.G.S., L.T., S.C.X.

## Competing interests

The authors have no competing interests to declare.

## Notes

### Competing Interest Statement

The authors have declared no competing interest.

## References

1. World_Health_Organisation. 2025. WHO World Malaria Report 2025. Geneva: World Health Organization; Licence: CC BY-NC-SA 30 IGO.

2. van der Pluijm RW, Imwong M, Chau NH, Hoa NT, Thuy-Nhien NT, Thanh NV, Jittamala P, Hanboonkunupakarn B, et al. 2019. Determinants of dihydroartemisinin-piperaquine treatment failure in *Plasmodium falciparum* malaria in Cambodia, Thailand, and Vietnam: a prospective clinical, pharmacological, and genetic study. Lancet Infect Dis 19:952–961.

3. Balikagala B, Fukuda N, Ikeda M, Katuro OT, Tachibana S-I, Yamauchi M, Opio W, Emoto S, et al. 2021. Evidence of artemisinin-resistant malaria in Africa. New England Journal of Medicine 385:1163–1171.

4. Rosenthal PJ, Asua V, Conrad MD. 2024. Emergence, transmission dynamics and mechanisms of artemisinin partial resistance in malaria parasites in Africa. Nature Reviews Microbiology 22:373–384.

5. Björkman A, Benn CS, Aaby P, Schapira A. 2023. RTS,S/AS01 malaria vaccine-proven safe and effective? Lancet Infect Dis 23:e318–e322.

6. Burrows JN, Duparc S, Gutteridge WE, Hooft van Huijsduijnen R, Kaszubska W, Macintyre F, Mazzuri S, Möhrle JJ, et al. 2017. New developments in anti-malarial target candidate and product profiles. Malaria Journal 16:26.

7. Xie SC, Tai C-W, Morton CJ, Ma L, Huang S-C, Wittlin S, Du Y, Hu Y, et al. 2024. A potent and selective reaction hijacking inhibitor of *Plasmodium falciparum* tyrosine tRNA synthetase exhibits single dose oral efficacy in vivo. PLoS Pathog 20:e1012429.

8. Xie SC, Metcalfe RD, Dunn E, Morton CJ, Huang SC, Puhalovich T, Du Y, Wittlin S, et al. 2022. Reaction hijacking of tyrosine tRNA synthetase as a new whole-of-life-cycle antimalarial strategy. Science 376:1074–1079.

9. Xie SC, Shi Y, Kobe B, Morton CJ, Tilley L. 2026. Reaction hijacking of enzymes to generate potent therapeutic modulators in situ. Trends Biochem Sci 51:439–456.

10. Xie SC, Tai C-W, Morton CJ, Ma L, Huang S-C, Wittlin S, Du Y, Hu Y, et al. 2024. A potent and selective reaction hijacking inhibitor of *Plasmodium falciparum* tyrosine tRNA synthetase exhibits single dose oral efficacy in vivo. PLOS Pathogens 20:e1012429.

11. Xie SC, Wang Y, Morton CJ, Metcalfe RD, Dogovski C, Pasaje CFA, Dunn E, Luth MR, et al. 2024. Reaction hijacking inhibition of *Plasmodium falciparum* asparagine tRNA synthetase. Nat Commun 15:937.

12. Luo Z, Qiu H, Peng X, Tan Q, Chen B, Gu Q, Liu H, Zhou H. 2025. Development of potent inhibitors targeting bacterial prolyl-tRNA synthetase through fluorine scanning-directed activity tuning. Eur J Med Chem 291:117647.

13. Baragaña B, Hallyburton I, Lee MC, Norcross NR, Grimaldi R, Otto TD, Proto WR, Blagborough AM, et al. 2015. A novel multiple-stage antimalarial agent that inhibits protein synthesis. Nature 522:315–20.

14. Cowell AN, Istvan ES, Lukens AK, Gomez-Lorenzo MG, Vanaerschot M, Sakata-Kato T, Flannery EL, Magistrado P, et al. 2018. Mapping the malaria parasite druggable genome by using in vitro evolution and chemogenomics. Science 359:191–199.

15. Luth MR, Gupta P, Ottilie S, Winzeler EA. 2018. Using in vitro evolution and whole genome analysis to discover next generation targets for antimalarial drug discovery. ACS Infect Dis 4:301–314.

16. Kümpornsin K, Kochakarn T, Yeo T, Okombo J, Luth MR, Hoshizaki J, Rawat M, Pearson RD, et al. 2023. Generation of a mutator parasite to drive resistome discovery in *Plasmodium falciparum*. Nat Commun 14:3059.

17. Carrasquilla M, Drammeh NF, Rawat M, Sanderson T, Zenonos Z, Rayner JC, Lee MCS. 2022. Barcoding genetically distinct *Plasmodium falciparum* strains for comparative assessment of fitness and antimalarial drug resistance. mBio 13:e0093722.

18. Ferreira LT, Cassiano GC, Alvarez LCS, Okombo J, Calit J, Fontinha D, Gil-Iturbe E, Coyle R, et al. 2024. A novel 4-aminoquinoline chemotype with multistage antimalarial activity and lack of cross-resistance with PfCRT and PfMDR1 mutants. PLoS Pathog 20:e1012627.

19. Fagbami L, Deik AA, Singh K, Santos SA, Herman JD, Bopp SE, Lukens AK, Clish CB, et al. 2019. The adaptive proline response in *P. falciparum* is independent of *Pf*eIK1 and eIF2α signaling. ACS Infect Dis 5:515–520.

20. Castilho BA, Shanmugam R, Silva RC, Ramesh R, Himme BM, Sattlegger E. 2014. Keeping the eIF2 alpha kinase Gcn2 in check. Biochim Biophys Acta 1843:1948–68.

21. Sugawara A, Tanaka T, Hirose T, Ishiyama A, Iwatsuki M, Takahashi Y, Otoguro K, Ōmura S, et al. 2013. Borrelidin analogues with antimalarial activity: design, synthesis and biological evaluation against *Plasmodium falciparum* parasites. Bioorg Med Chem Lett 23:2302–5.

22. Ganesan SM, Falla A, Goldfless SJ, Nasamu AS, Niles JC. 2016. Synthetic RNA-protein modules integrated with native translation mechanisms to control gene expression in malaria parasites. Nat Commun 7:10727.

23. Ibba M, Soll D. 2000. Aminoacyl-tRNA synthesis. Annu Rev Biochem 69:617–50.

24. Mirdita M, Schutze K, Moriwaki Y, Heo L, Ovchinnikov S, Steinegger M. 2022. ColabFold: making protein folding accessible to all. Nat Methods 19:679–682.

25. Jumper J, Evans R, Pritzel A, Green T, Figurnov M, Ronneberger O, Tunyasuvunakool K, Bates R, et al. 2021. Highly accurate protein structure prediction with AlphaFold. Nature 596:583–589.

26. Cusack S, Hartlein M, Leberman R. 1991. Sequence, structural and evolutionary relationships between class 2 aminoacyl-tRNA synthetases. Nucleic Acids Res 19:3489–98.

27. Marck C, Grosjean H. 2002. tRNomics: analysis of tRNA genes from 50 genomes of Eukarya, Archaea, and Bacteria reveals anticodon-sparing strategies and domain-specific features. Rna 8:1189–232.

28. Gardner MJ, Shallom SJ, Carlton JM, Salzberg SL, Nene V, Shoaibi A, Ciecko A, Lynn J, et al. 2002. Sequence of *Plasmodium falciparum* chromosomes 2, 10, 11 and 14. Nature 419:531–4.

29. Xie SC, Griffin MDW, Winzeler EA, Ribas de Pouplana L, Tilley L. 2023. Targeting aminoacyl tRNA synthetases for antimalarial drug development. Annu Rev Microbiol 77:111–129.

30. Kobayashi T, Takimura T, Sekine R, Kelly VP, Kamata K, Sakamoto K, Nishimura S, Yokoyama S. 2005. Structural snapshots of the KMSKS loop rearrangement for amino acid activation by bacterial tyrosyl-tRNA synthetase. J Mol Biol 346:105–17.

31. Vondenhoff GH, Van Aerschot A. 2011. Aminoacyl-tRNA synthetase inhibitors as potential antibiotics. Eur J Med Chem 46:5227–36.

32. Lux MC, Standke LC, Tan DS. 2019. Targeting adenylate-forming enzymes with designed sulfonyladenosine inhibitors. J Antibiot (Tokyo) 72:325–349.

33. Kim S, Lee SW, Choi EC, Choi SY. 2003. Aminoacyl-tRNA synthetases and their inhibitors as a novel family of antibiotics. Applied Microbiology and Biotechnology 61:278–288.

34. Tao J, Schimmel P. 2000. Inhibitors of aminoacyl-tRNA synthetases as novel anti-infectives. Expert Opinion on Investigational Drugs 9:1767–1775.

35. Hurdle Julian G, O’Neill Alexander J, Chopra I. 2005. Prospects for aminoacyl-tRNA synthetase inhibitors as new antimicrobial agents. Antimicrobial Agents and Chemotherapy 49:4821–4833.

36. Teng M, Hilgers MT, Cunningham ML, Borchardt A, Locke JB, Abraham S, Haley G, Kwan BP, et al. 2013. Identification of bacteria-selective threonyl-tRNA synthetase substrate inhibitors by structure-based design. J Med Chem 56:1748–60.

37. Charlton MH, Aleksis R, Saint-Leger A, Gupta A, Loza E, Ribas de Pouplana L, Kaula I, Gustina D, et al. 2018. N-Leucinyl benzenesulfonamides as structurally simplified leucyl-tRNA synthetase inhibitors. ACS Medicinal Chemistry Letters 9:84–88.

38. Zhang F, Du J, Wang Q, Hu Q, Zhang J, Ding D, Zhao Y, Yang F, et al. 2013. Discovery of N-(4-sulfamoylphenyl)thioureas as *Trypanosoma brucei* leucyl-tRNA synthetase inhibitors. Organic & Biomolecular Chemistry 11:5310–5324.

39. Filisetti D, Théobald-Dietrich A, Mahmoudi N, Rudinger-Thirion J, Candolfi E, Frugier M. 2013. Aminoacylation of *Plasmodium falciparum* tRNA(Asn) and insights in the synthesis of asparagine repeats. J Biol Chem 288:36361–71.

40. Nass G, Poralla K, Zahner H. 1969. Effect of the antibiotic Borrelidin on the regulation of threonine biosynthetic enzymes in *E. coli*. Biochem Biophys Res Commun 34:84–91.

41. Meanwell NA. 2018. Fluorine and fluorinated motifs in the design and application of bioisosteres for drug design. Journal of Medicinal Chemistry 61:5822–5880.

42. Gillis EP, Eastman KJ, Hill MD, Donnelly DJ, Meanwell NA. 2015. Applications of fluorine in medicinal chemistry. Journal of Medicinal Chemistry 58:8315–8359.

43. Richardson P. 2021. Applications of fluorine to the construction of bioisosteric elements for the purposes of novel drug discovery. Expert Opinion on Drug Discovery 16:1261–1286.

44. Ketprasit N, Tai C-W, Sharma VK, Manickam Y, Khandokar Y, Ye X, Dogovski C, Hilko DH, et al. 2025. Natural product-mediated reaction hijacking mechanism validates *Plasmodium* aspartyl-tRNA synthetase as an antimalarial drug target. PLoS Pathog 21: e1013057.

45. Duffey M, Blasco B, Burrows JN, Wells TNC, Fidock DA, Leroy D. 2021. Assessing risks of *Plasmodium falciparum* resistance to select next-generation antimalarials. Trends in Parasitology 37:709–721.

46. Frame IJ, Merino EF, Schramm VL, Cassera MB, Akabas MH. 2012. Malaria parasite type 4 equilibrative nucleoside transporters (ENT4) are purine transporters with distinct substrate specificity. Biochem J 446:179–90.

47. Kenthirapalan S, Waters AP, Matuschewski K, Kooij TWA. 2016. Functional profiles of orphan membrane transporters in the life cycle of the malaria parasite. Nature Communications 7:10519.

48. Deveci G, Kamil M, Kina U, Temel BA, Aly ASI. 2023. Genetic disruption of nucleoside transporter 4 reveals its critical roles in malaria parasite sporozoite functions. Pathogens and Global Health 117:284–292.

49. Ruff M, Krishnaswamy S, Boeglin M, Poterszman A, Mitschler A, Podjarny A, Rees B, Thierry JC, et al. 1991. Class II aminoacyl transfer RNA synthetases: crystal structure of yeast aspartyl-tRNA synthetase complexed with tRNA(Asp). Science 252:1682–9.

50. Gamo FJ, Sanz LM, Vidal J, de Cozar C, Alvarez E, Lavandera JL, Vanderwall DE, Green DV, et al. 2010. Thousands of chemical starting points for antimalarial lead identification. Nature 465:305–10.

51. Love MI, Huber W, Anders S. 2014. Moderated estimation of fold change and dispersion for RNA-seq data with DESeq2. Genome Biol 15:550.

52. Reader J, Botha M, Theron A, Lauterbach SB, Rossouw C, Engelbrecht D, Wepener M, Smit A, et al. 2015. Nowhere to hide: interrogating different metabolic parameters of *Plasmodium falciparum* gametocytes in a transmission blocking drug discovery pipeline towards malaria elimination. Malar J 14:213.

53. Reader J, van der Watt ME, Birkholtz LM. 2022. Streamlined and robust stage-specific profiling of gametocytocidal compounds against *Plasmodium falciparum*. Front Cell Infect Microbiol 12:926460.

54. Delves MJ, Straschil U, Ruecker A, Miguel-Blanco C, Marques S, Dufour AC, Baum J, Sinden RE. 2016. Routine in vitro culture of *P. falciparum* gametocytes to evaluate novel transmission-blocking interventions. Nature Protocols 11:1668–1680.

55. Nasamu AS, Falla A, Pasaje CFA, Wall BA, Wagner JC, Ganesan SM, Goldfless SJ, Niles JC. 2021. An integrated platform for genome engineering and gene expression perturbation in *Plasmodium falciparum*. Sci Rep 11:342.

56. Deitsch K, Driskill C, Wellems T. 2001. Transformation of malaria parasites by the spontaneous uptake and expression of DNA from human erythrocytes. Nucleic Acids Res 29:850–3.

57. Ritz C, Baty F, Streibig JC, Gerhard D. 2015. Dose-response analysis using R. PLoS One 10:e0146021.

58. Bridgford JL, Xie SC, Cobbold SA, Pasaje CFA, Herrmann S, Yang T, Gillett DL, Dick LR, et al. 2018. Artemisinin kills malaria parasites by damaging proteins and inhibiting the proteasome. Nature Communications 9:3801.

59. Eiler S, Dock-Bregeon A, Moulinier L, Thierry JC, Moras D. 1999. Synthesis of aspartyl-tRNA(Asp) in *Escherichia coli* - a snapshot of the second step. EMBO J 18:6532–41.

