## Supplementary information for "Potent reaction hijacking inhibitors of *Plasmodium falciparum* asparagine tRNA synthetase"

**MBio**

Xi Ye *et al.*

**The PDF file includes:**

S1-S11 Figures
S1-S9 Tables
Chemistry Methods
Supplementary References

### Supplementary Figures

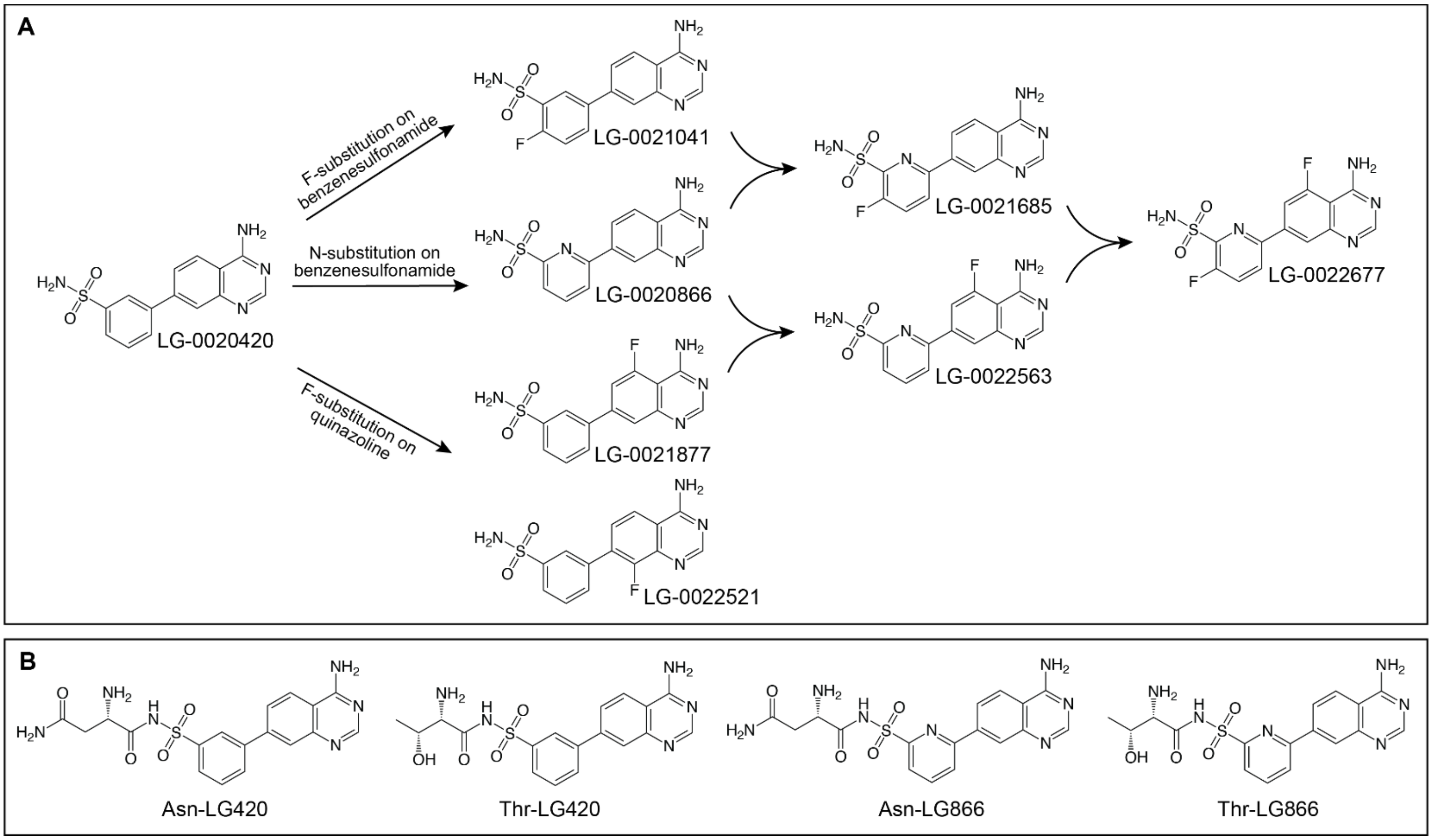

**Supplementary Figure 1. Structures of 4AQS series and adducts examined in this study.**

(A) Structures and logic pathway for synthesis of LG-0020420 (MMV2323511), LG-0021041 (MMV2480986), LG-0020866 (MMV2480572), LG-0021877 (MMV2482899), LG-0022521 (MMV2503069), LG-0021685 (MMV2482269), LG-0022563 (MMV2503071) and LG-0022677 (MMV2503074). (B) Amino acid adducts were generated as controls: Asn-LG420 (LG-0020957; MMV2480808), Thr-LG420 (LG-0020445; MMV2323518), Asn-LG866 (LG-0021200; MMV2481583) and Thr-LG866 (LG-0021186; MMV2481220).

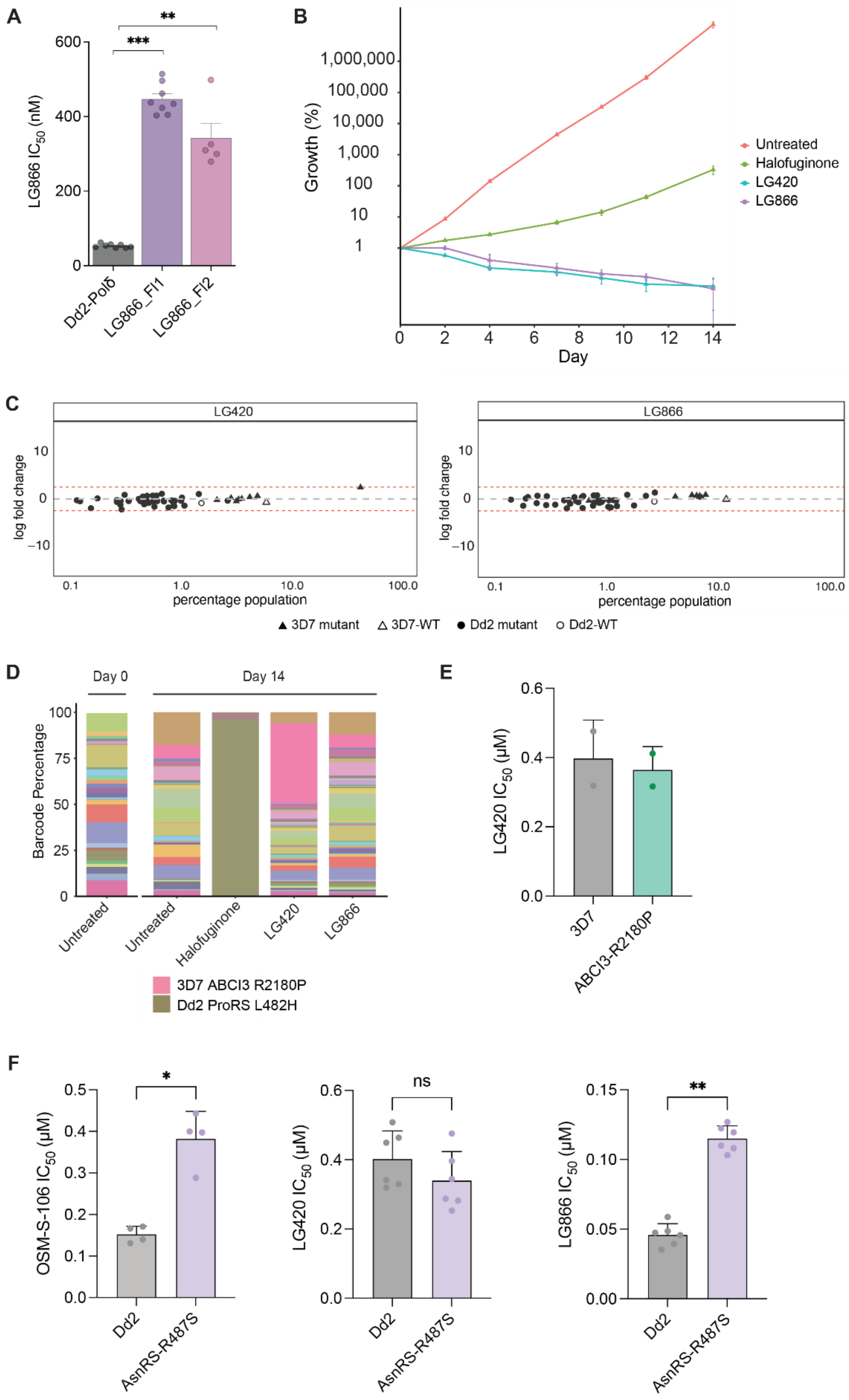

**Supplementary Figure 2. Susceptibility of 4AQS series members to resistance development and cross-resistance**

A) Drug susceptibility of Dd2-Polδ recrudescent parasites exposed to LG866 compared to the parent strain. The $IC_{50}$ values (nM) represent the mean ± SEM from 5-8 independent assays, each performed in duplicate. Statistical significance was determined for all samples using two-tailed Mann-Whitney U tests (GraphPad Prism, version 10). $^{**}p < 0.01$, $^{***}p < 0.001$. See Supplementary Table 3 for data values. (B-E) Cross-resistance profiling of LG420 and LG866 using the AReBar assay. (B) Growth curves for LG420, LG866, the positive control halofuginone, and a no-drug control. (C) $Log_2$ fold change (LFC) values for each barcoded line relative to the no-drug control following treatment with LG420 and LG866. Dotted lines indicate 2.5 LFC. (D) Proportion of each barcoded line in the pool at day 0 and at day 14 after treatment with LG420, LG866 or halofuginone. (E) Bar graph showing the drug response to the $ABCI3^{R2180P}$ mutant cell line (weak signal identified from AReBar analysis) following treatment with LG866. (F) Sensitivity to exposure (72-h) to LG420, LG866 and OSM-S-106 for a wildtype parental line (Dd2) and a CRISPR-edited clone harbouring *Pf*$AsnRS^{R487S}$. Data represent means from 4-6 independent experiments and error bars correspond to SD.

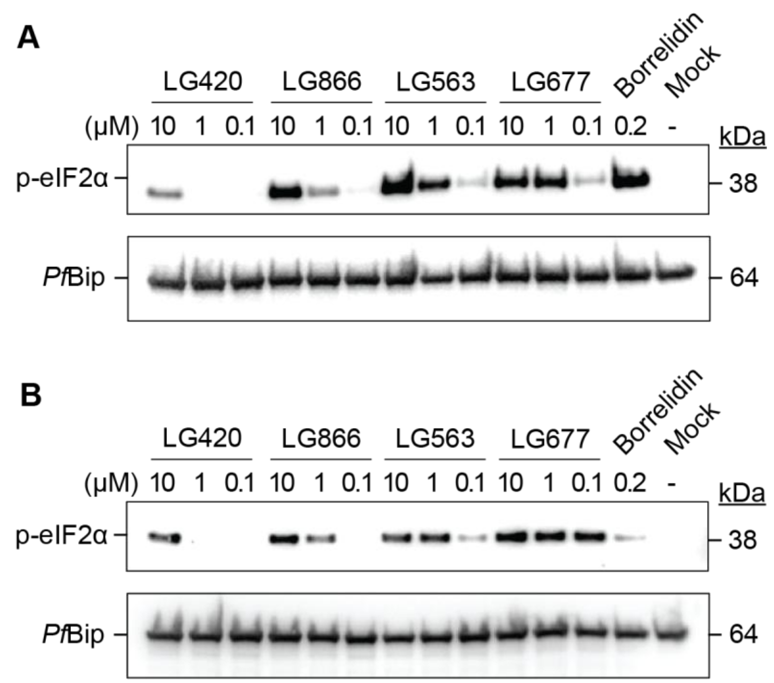

**Supplementary Figure 3. Additional Western blot analysis of eIF2α phosphorylation in 4AQS-treated *P. falciparum***

(A,B) Two additional analyses of trophozoite-stage *P. falciparum* 3D7 cultures that were exposed to 0.2 µM borrelidin or different concentrations of LG420, LG866, LG563, and LG677 for 3 h. Western blot analysis was performed on parasite extracts to detect phosphorylated eIF2α. *Pf*BiP served as the loading control. Blots are related to Figure 1.

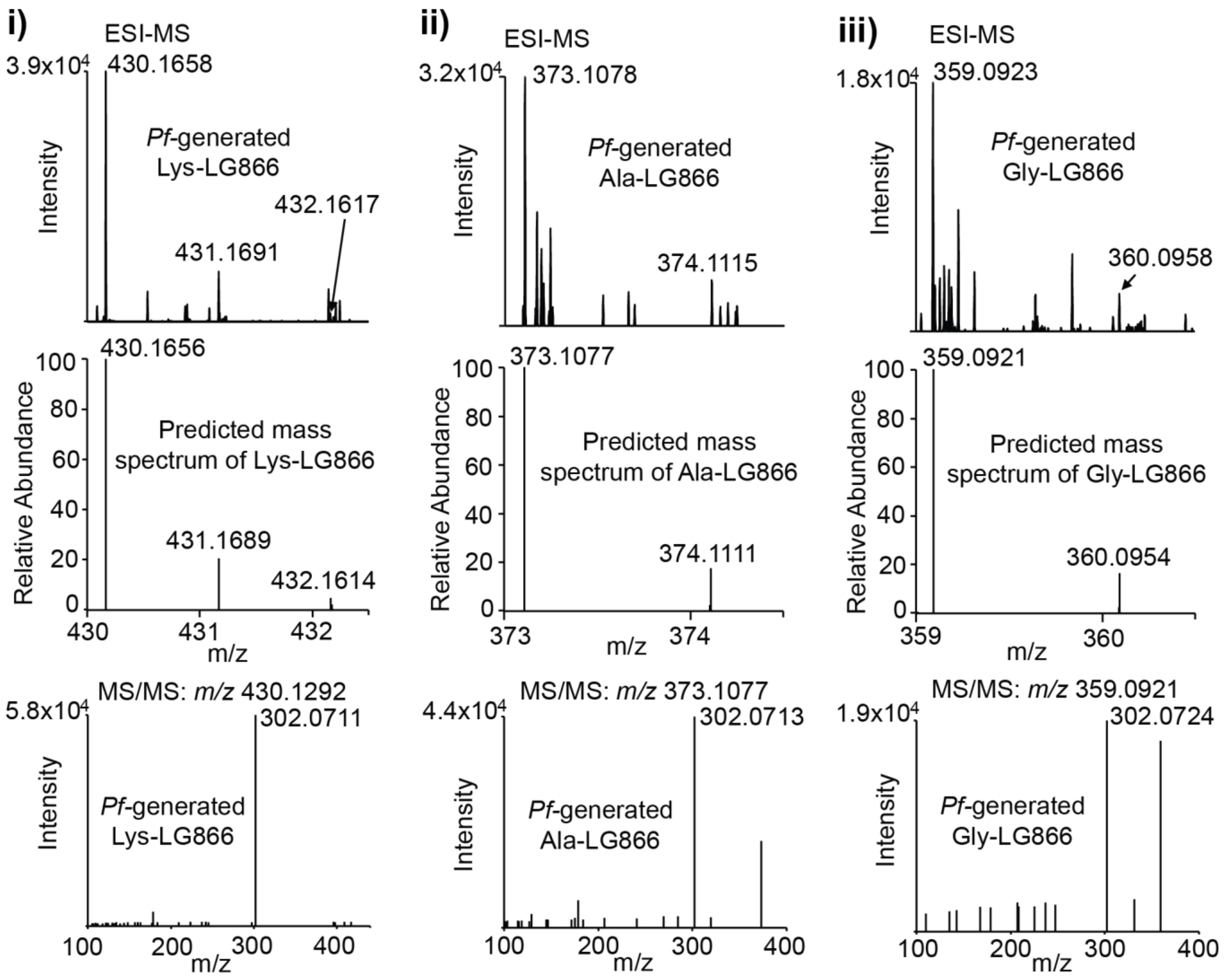

**Supplementary Figure 4. Targeted mass spectrometry analysis of minor amino acid adducts in LG866-treated *P. falciparum***

Late trophozoite stage *P. falciparum* 3D7 cultures were exposed to LG866 at a concentration of 10 µM and extracts were subjected to LC-MS analysis. Detected (top panels) and predicted (middle panels) mass spectra, and MS/MS fragmentation spectra (bottom panels) for the minor amino acid adducts are shown: (i) Lys-LG866, (ii) Ala-LG866 and (iii) Gly-LG866.

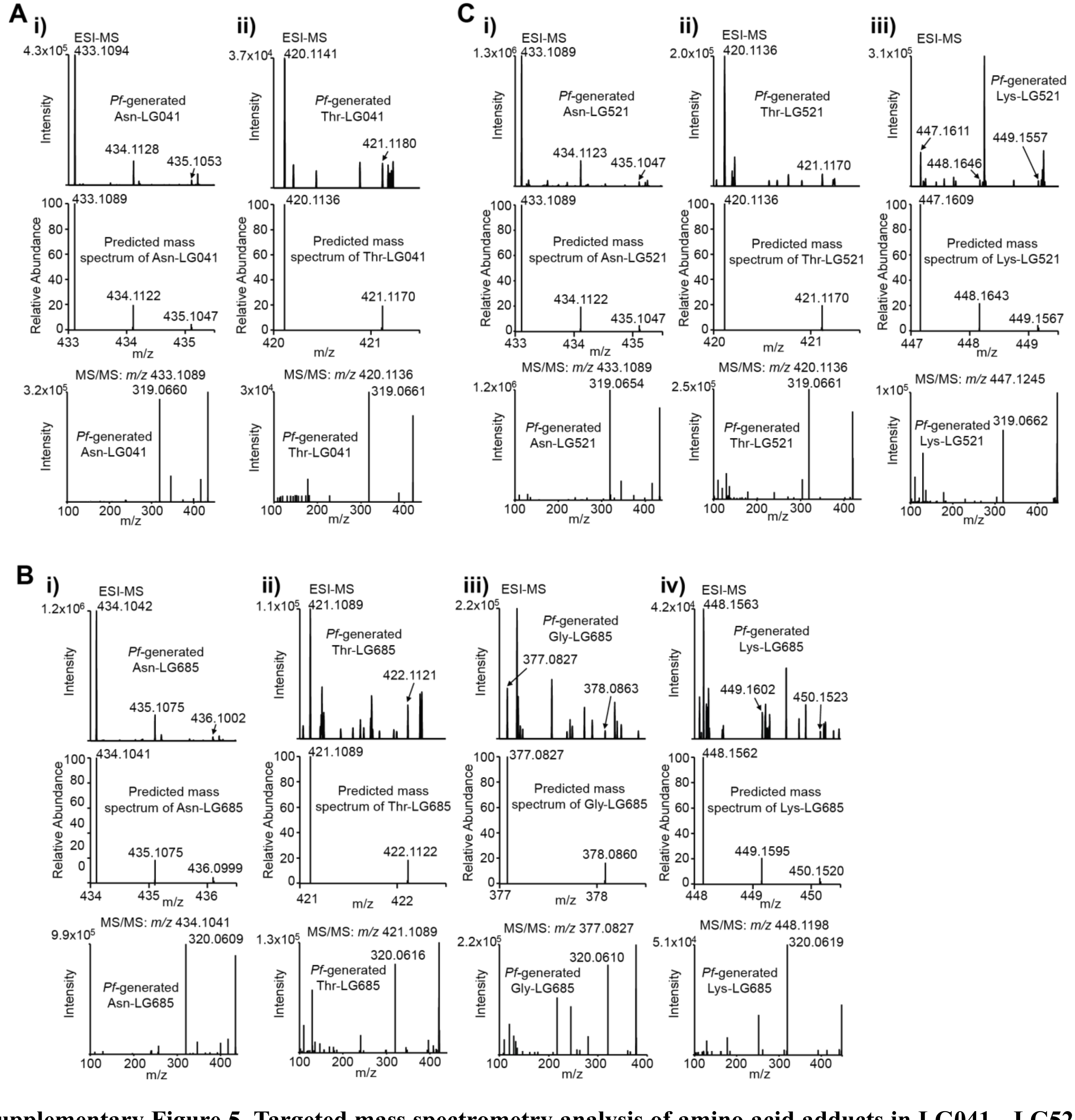

**Supplementary Figure 5. Targeted mass spectrometry analysis of amino acid adducts in LG041-, LG521- and LG685-treated *P. falciparum***

Late trophozoite stage *P. falciparum* 3D7 cultures were exposed to LG041, LG521 and LG685 at a concentration of 10 µM and extracts were subjected to LC-MS analysis. (A) Detected (top panels) and predicted (middle panels) mass spectra, and MS/MS fragmentation spectra (bottom panels), for Asn-LG041 (i) and Thr-LG041 (ii). (B) Detected (left panels) and predicted (middle panels) mass spectra, and MS/MS fragmentation spectra (right panels), for (i) Asn-LG685, (ii) Thr-LG685, (iii) Gly-LG866 and (iv) Lys-LG866. (C) Detected (left panels) and predicted (middle panels) mass spectra, and MS/MS fragmentation spectra (right panels), for (i) Asn-LG521, (ii) Thr-LG521 and (iii) Lys-LG521.

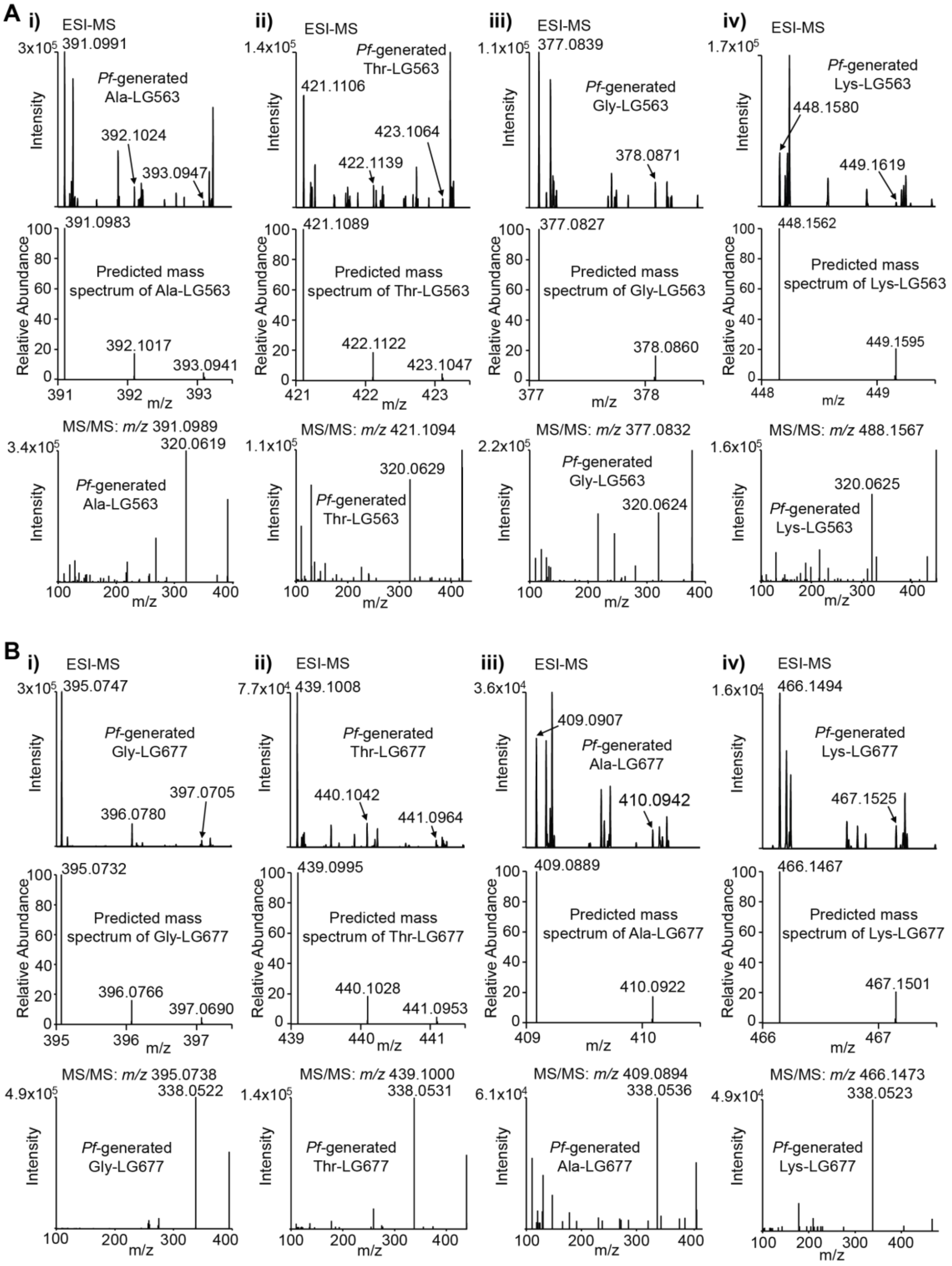

**Supplementary Figure 6. Targeted mass spectrometry analysis of minor amino acid adducts in LG563- and LG677-treated *P. falciparum***

Late trophozoite stage *P. falciparum* 3D7 cultures were exposed to LG563 and LG677 at a concentration of 10 µM and extracts were subjected to LC-MS analysis. (A) Detected (top panels) and predicted (middle panels) mass spectra, and MS/MS fragmentation spectra (bottom panels) for the minor amino acid-LG563 adducts are shown: (i) Ala-LG563, (ii) Thr-LG563, (iii) Gly-L563 and (iv) Lys-LG563. (B) Detected (top panels) and predicted (middle panels) mass spectra, and MS/MS fragmentation spectra (bottom panels) for the minor amino acid-LG677 adducts are shown: (i) Ala-LG563, (ii) Thr-LG563, (iii) Gly-L563 and (iv) Lys-LG563.

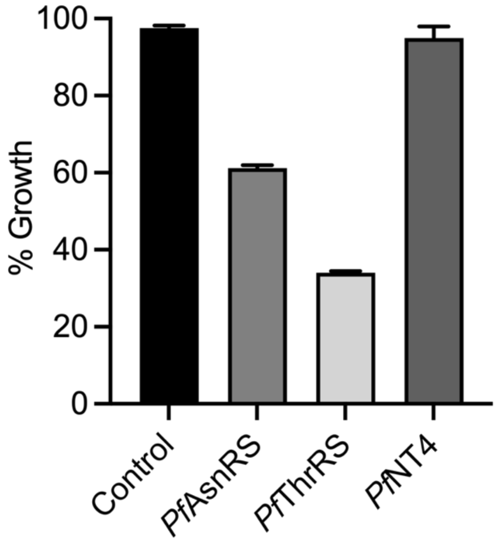

**Supplementary Figure 7. Effect of target knock-down on parasite growth**

Growth of aptamer-regulatable *Pf*AsnRS, *Pf*ThrRS and *Pf*NT4 lines was assessed over 72 h, relative to aTc-treated controls. Error bars correspond to range of 2 independent experiments, with each experiment containing 6 technical replicates.

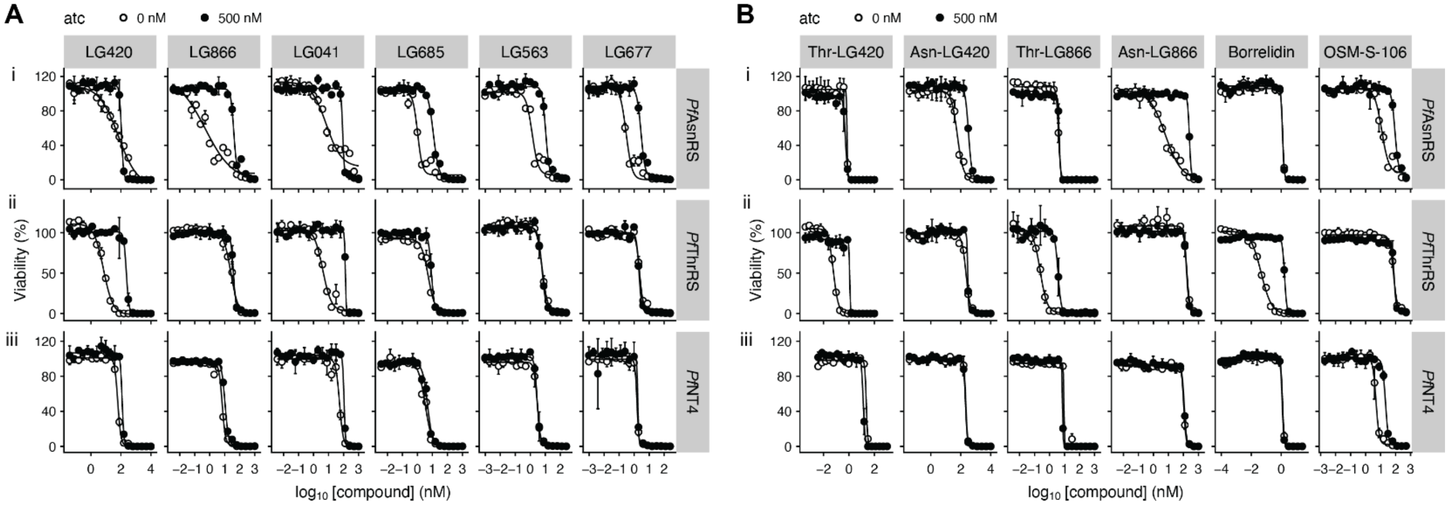

**Supplementary Figure 8. Effect of target knock-down on parasite growth in presence of 4AQS series compounds**

Sensitivity to exposure (72-h) to different hijackers (A) or adducts and controls (B) for aptamer-regulatable *Pf*AsnRS (i) and *Pf*AsnRS (ii) and *Pf*NT4 (iii) lines upon addition of aTc (closed circles) and with the target expression reduced (open circles), with data normalized to a no drug control. Amino acid adducts, Asn-LG420 (LG0020957), Thr-LG420 (LG-0020445), Asn-LG866 (LG-0021200) and Thr-LG866 (LG-0021186), borrelidin and OSM-S-106 are used as controls. Plots show representative experiments from at least three biological replicates. Data represent the mean of two technical replicates and error bars correspond to SD.

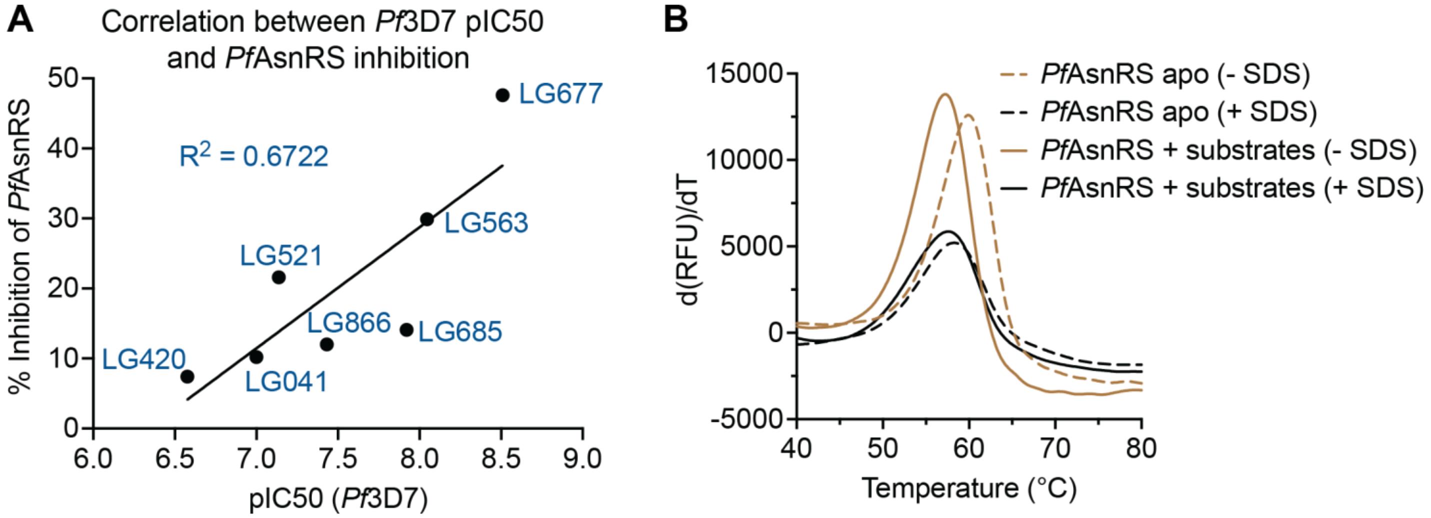

**Supplementary Figure 9. Correlation of cell growth inhibition and enzyme inhibition and effect of SDS of *P. falciparum* AsnRS stability.**

(A) Correlation of percentage inhibition of *Pf*AsnRS enzyme activity in an *in vitro* assay with $pIC_{50}$ value for inhibition of growth of *P. falciparum* cultures. (B) Apo *Pf*AsnRS (1.5 µM) and *Pf*AsnRS (1.5 µM) in the presence of 20 µM ATP, 40 µM Asn, 80 µM *Ec*tRNA were incubated at 37°C for 3 h. First derivatives of melting curves determined in the absence or presence of 0.001% SDS. Data are representative of 3-7 independent experiments.

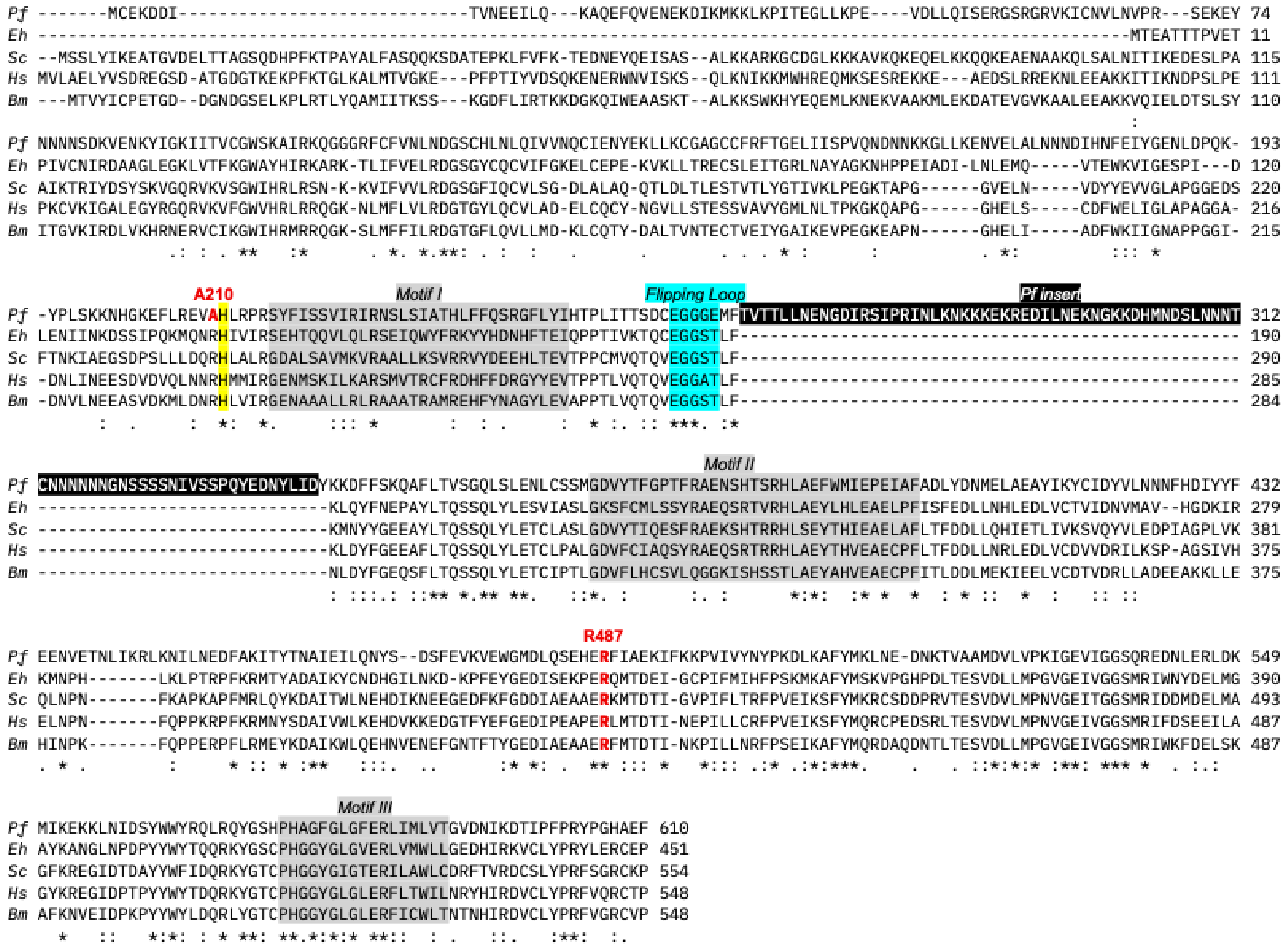

**Supplementary Figure 10. Sequence alignment of AsnRS sequences from different species.**

Type II aaRSs exhibit three motifs (I-III, grey shading) involved in ATP binding and dimerization. Alignment of AsnRS sequences from *Plasmodium falciparum* (*Pf*), *Entamoeba histolytica* (*Eh*), *Saccharomyces cerevisiae* (*Sc*), *Homo sapiens* (*Hs*) and *Brugia malayi* (*Bm*) reveals a high level of conservation of these motifs. *P. falciparum* features a large insert (highlighted in black, white font). The flipping loop that has previously been shown to undergo dynamic motions that facilitate tRNA binding [1] is highlighted in cyan. *Pf*AsnRS Arg487 (red) is mutated to Ser upon selection against OSM-S-106 [2]. *Pf*AsnRS Ala210 (red) is mutated to Thr upon selection against LG866. Ala210 is adjacent to a conserved His (yellow highlight).

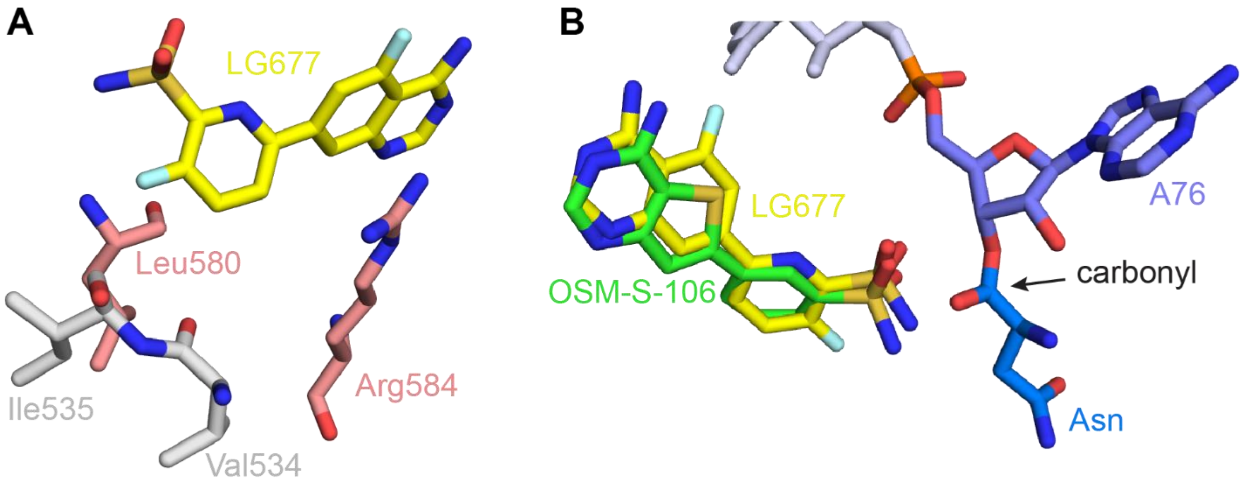

**Supplementary Figure 11. A *Pf*AsnRS-Asn-tRNA complex model with docked OSM-S-106 and LG677**

(A) Active site residues in the LG677-docked pocket. (B) Close-up view of the active site with 3' terminal A76 of the tRNA and representative in silico docks of OSM-S-106 and LG677 to the *Pf*AsnRS-Asn-tRNA model. The 4-amino-thienopyrimidine and 4-amino-quinazoline groups adopt similar poses.

### Supplementary Tables

**Supplementary Table 1. Activity against immature (>90% stage II/III) and mature (>95% stage V) stage gametocytes (Pf3D7*elo1-pfs16*-CBG99) and activity against transmissible gametes.** Data for gametocyte killing are from three independent biological repeats (n = 3), each performed in technical triplicates, mean ± S.E. Data for gamete inactivation are from two independent biological repeats.

| **Compound** | **Immature-stage gametocytes $IC_{50\_48h}$ (nM)** | **Mature-stage gametocytes (% inhibition at 10 µM or $IC_{5\_48h}$ (nM))** |
|---|---|---|
| **LG563** | 224 ± 5 | 18% |
| **LG677** | 199 ± 30 | 32% |
| **MMV390048** | 379 ± 16 | 162 ± 9 |
| **Methylene Blue** | 66 ± 3 | 420 ± 58 |
| **Compound** | **Dual gamete formation assay (male) $IC_{50\_48h}$ (nM)** | **Dual gamete formation assay (female) $IC_{50\_48h}$ (nM)** |
| **LG563** | 654/661 (n = 2) | 3,650/8,690 (n = 2) |
| **LG677** | 382/363 (n = 2) | 1,220/3,800 (n = 2) |
| **Cabamiquine (DDD498)*** | 5.2/3.8 (n = 2) | 1.2/1.1 (n = 2) |
| *Data for the DDD498 control is very similar to a previous report [3], validating the assay. | | |

**Supplementary Table 2. Summary of resistance selection using standard Minimum Inoculum for Resistance (MIR) protocol.** Dd2-B2 and Dd2-Polδ parasites were subjected to pressure with LG866 at 3 x $IC_{90}$. Mean $IC_{50}$ = 48 nM; Mean $IC_{90}$ = 95 nM. Parasites were subjected to a single-step selection at 3 x $IC_{90}$.

| **Parasite line** | **Dd2-B2** | **Dd2-Polδ** |
|---|---|---|
| **Number of parasites in the selection** | $10^9$ | $10^9$ |
| **Day of recrudescence** | No recrudescence | Day 42 |
| **$IC_{50}$ fold shift (clones)** | N.A. | ~6-8-fold |
| **Minimum Inoculum for Resistance (MIR)** | >$10^9$ | ~4x$10^9$ (predicted)* |

* MIR prediction based on Dd2-Polδ having a mutation rate ~8-10 fold higher than Dd2-B2

**Supplementary Table 3. $IC_{50}$ values for LG866-selected Dd2- Polδ lines.** $IC_{50}$ (nM) for LG-0020866 are shown for the drug-sensitive Dd2-Polδ parental strain and the LG866 selected asparagine-tRNA ligase mutant lines. Data represent means ± SEM from 6 independent experiments, each performed in duplicate. Statistical significance was determined using the two-tailed Mann–Whitney *U* test (asterisk (*) indicates $p < 0.05$).

| **Sample** | **N** | **$IC_{50}$ ± SEM (nM)** |
|---|---|---|
| **Dd2-Polδ** | 8 | 53 ± 1.8 |
| **LG866_Fl1** | 8 | 447 ± 14.4 |
| **LG866_Fl2** | 5 | 343 ± 33.6 |

**Supplementary Table 4. List of mutations from whole-genome sequencing**

| Mutation in | Gene ID | Gene Name | Amino acid change | Codon change | Fl1.alt_AB | Fl2.alt_AB | Dd2-PolD.alt_AB |
|---|---|---|---|---|---|---|---|
| **F1, F2** | PF3D7_0211800 | asparagine--tRNA ligase | A210T | Gca/Aca | 1 | 1 | 0 |
| **F2** | PF3D7_0415200 | Conserved Plasmodium protein, unknown function | N762H | Aat/Cat | 0 | 0.9 | 0 |
| **F2** | PF3D7_0418300 | Conserved Plasmodium protein, unknown function | D856V | gAt/gTt | 0 | 0.94 | 0 |
| **F2** | PF3D7_0513900 | Conserved protein, unknown function | L210I | Ctc/Atc | 0 | 1 | 0 |
| **F1** | PF3D7_0526700 | Conserved protein, unknown function | D626N | Gat/Aat | 1 | 0 | 0 |
| **F2** | PF3D7_0919000 | Nucleosome assembly protein | D223Y | Gat/Tat | 0 | 1 | 0 |
| **F2** | PF3D7_1121400 | WD repeat-containing protein, putative | A259T | Gct/Act | 0 | 0.95 | 0 |
| **F1** | PF3D7_1123500 | Golgi protein 2 | Y1099D | Tat/Gat | 1 | 0 | 0 |
| **F2** | PF3D7_1234400 | Microgamete surface protein MiGS, putative | F342C | tTt/tGt | 0 | 1 | 0 |
| **F1** | PF3D7_1329000 | DNA-directed RNA polymerase III | G968C | Ggt/Tgt | 0.3 | 0 | 0 |

| | | | | | | | |
|---|---|---|---|---|---|---|---|
| | | subunit RPC1, putative | | | | | |
| **F1** | PF3D7_1346400 | VPS13 domain-containing protein, putative | M5502I | atG/atT | 1 | 0 | 0 |
| **F1** | PF3D7_1424400 | 60S ribosomal protein L7-3, putative | M264I | atG/atT | 1 | 0 | 0 |

**Supplementary Table 5: Representation of barcoded mutant parasite lines within the AReBar library population prior to and following selection with LG420 and LG866.**

| Parasite line | Gene Description | Gene ID | Day 0(%) | Day 14 (%) | | |
|---|---|---|---|---|---|---|
| | | | Untreated | Untreated | LG-0020420 | LG-0020866 |
| 3D7 | Wild type | | 9.641 | 17.811 | 6.048 | 12.148 |
| 3D7 ABCI3 R2180P | ABC transporter I family member 1 | PF3D7_0319700 | 5.755 | 7.276 | 43.199 | 6.983 |
| 3D7 ACS10 M300I | Acyl CoA synthase | PF3D7_0525100 | 4.375 | 2.497 | 2.157 | 4.230 |
| 3D7 ACS11 D648Y | Acyl CoA synthase | PF3D7_1238800 | 1.381 | 0.245 | 0.458 | 0.708 |
| 3D7 ATP2 CNV2 | Phospholipid-transporting ATP2 | PF3D7_1219600 | 6.318 | 7.170 | 4.281 | 7.015 |
| 3D7 DHFR-TS G378E | Dihydrofolate reductase-thymidylate synthase | PF3D7_0417200 | 4.681 | 9.776 | 3.232 | 7.319 |
| 3D7 DHFR-TS I403L | Dihydrofolate reductase-thymidylate synthase | PF3D7_0417200 | 5.288 | 7.794 | 3.394 | 7.033 |
| 3D7 FTb A515T | Farnesyltransferase subunit beta | PF3D7_1147500 | 6.270 | 7.038 | 3.623 | 7.930 |
| 3D7 MDR2 K840N | Multidrug resistance protein 2 | PF3D7_1447900 | 5.020 | 4.362 | 2.895 | 5.971 |
| 3D7 NCR1 A1108T | Niemann-Pick type C1-related protein | PF3D7_0107500 | 5.946 | 7.353 | 5.026 | 6.575 |
| Dd2 | Wild type | | 3.083 | 2.245 | 1.560 | 2.741 |
| Dd2 AcAS A597V | Acetyl-CoA synthetase | PF3D7_0627800 | 0.490 | 0.051 | 0.341 | 0.181 |
| Dd2 AcAS T648M | Acetyl-CoA synthetase | PF3D7_0627800 | 1.016 | 1.386 | 0.585 | 0.770 |
| Dd2 AsnRS R487S | Asn-tRNA synthetase | PF3D7_0509600 | 1.069 | 0.425 | 0.893 | 2.301 |
| Dd2 ATP4 G358S | Non-SERCA-type Ca2+ -transporting P-ATPase | PF3D7_1211900 | 0.784 | 0.432 | 0.494 | 0.351 |
| Dd2 ATP4 L927V | Non-SERCA-type Ca2+ -transporting P-ATPase | PF3D7_1211900 | 0.993 | 0.205 | 0.437 | 0.568 |
| Dd2 ATP4 Q172H | Non-SERCA-type Ca2+ -transporting P-ATPase | PF3D7_0321900 | 0.630 | 0.359 | 0.567 | 0.479 |
| Dd2 CARL I1139K-a | Cyclic amine resistance locus | PF3D7_0321900 | 0.002 | 0.001 | 0.000 | 0.000 |
| Dd2 CARL I1139K-b | Cyclic amine resistance locus | PF3D7_0321900 | 0.167 | 0.090 | 0.124 | 0.051 |
| Dd2 CARL V1103L | Cyclic amine resistance locus | PF3D7_1114700 | 0.459 | 0.200 | 0.155 | 0.292 |
| Dd2 CLK3 H259P | Cyclin-dependent-like kinase | PF3D7_1438500 | 0.892 | 0.727 | 0.295 | 0.447 |

| | | | | | | |
|---|---|---|---|---|---|---|
| Dd2 CPSF Y408S E | Cleavage and polyadenylation specific factor | PF3D7_143 8500 | 0.633 | 0.135 | 0.501 | 0.522 |
| Dd2 CPSF Y408S S | Cleavage and polyadenylation specific factor | PF3D7_070 9000 | 0.503 | 0.001 | 0.314 | 0.199 |
| Dd2 CRT M343L | Chloroquine resistance transporter | PF3D7_125 0200 | 2.003 | 0.366 | 1.039 | 2.771 |
| Dd2 CSC1 L800P | CSC1-like protein, putative | PF3D7_MI T02300 | 0.668 | 0.082 | 0.459 | 0.628 |
| Dd2 cytBC1 G33V | Cytochrome b | PF3D7_MI T02300 | 1.585 | 0.035 | 0.565 | 0.794 |
| Dd2 cytBC1 V284L | Cytochrome b | PF3D7_041 7200 | 0.597 | 0.142 | 0.267 | 0.423 |
| Dd2 DHFR-TS S216R | Dihydrofolate reductase-thymidylate synthase | PF3D7_060 3300 | 1.247 | 0.228 | 0.619 | 0.847 |
| Dd2 DHODH C276Y | Dihydroorotate dehydrogenase | PF3D7_060 3300 | 0.762 | 0.144 | 0.422 | 0.780 |
| Dd2 DHODH F227I | Dihydroorotate dehydrogenase | PF3D7_060 3300 | 0.363 | 0.003 | 0.116 | 0.243 |
| Dd2 DHODH I263F | Dihydroorotate dehydrogenase | PF3D7_060 3300 | 0.067 | 0.000 | 0.065 | 0.036 |
| Dd2 DHODH L531F | Dihydroorotate dehydrogenase | PF3D7_145 1100 | 0.382 | 0.010 | 0.178 | 0.295 |
| Dd2 eEF2 L755F | Elongation factor 2 | PF3D7_145 1100 | 0.558 | 0.162 | 0.287 | 0.543 |
| Dd2 eEF2 Y186N | Elongation factor 2 | PF3D7_112 8400 | 0.472 | 0.016 | 0.268 | 0.239 |
| Dd2 GGPPS S228T | Geranylgeranyl diphosphate synthase | PF3D7_070 8400 | 0.678 | 0.179 | 0.683 | 0.609 |
| Dd2 HSP90 A41S | Heat shock protein 90 | PF3D7_133 2900 | 1.638 | 2.430 | 0.785 | 1.109 |
| Dd2 IleRS E180D | Ile-tRNA synthetase | PF3D7_133 2900 | 1.313 | 0.401 | 0.833 | 1.070 |
| Dd2 IleRS L810F | Ile-tRNA synthetase | PF3D7_133 2900 | 1.145 | 0.521 | 0.841 | 0.918 |
| Dd2 IleRS V500A | Ile-tRNA synthetase | PF3D7_134 3700 | 1.132 | 0.365 | 0.472 | 0.878 |
| Dd2 kelch13 C580C | Kelch protein K13 | PF3D7_134 3700 | 1.592 | 0.454 | 0.713 | 1.241 |
| Dd2 kelch13 C580Y | Kelch protein K13 | PF3D7_134 3700 | 1.144 | 0.436 | 0.490 | 0.942 |
| Dd2 kelch13 R539T | Kelch protein K13 | PF3D7_090 8800 | 0.506 | 0.074 | 0.288 | 0.463 |
| Dd2 MCP D195N | Mitochondrial carrier protein | PF3D7_090 8800 | 0.241 | 0.010 | 0.095 | 0.140 |
| Dd2 MCP P214T | Mitochondrial carrier protein | PF3D7_050 9800 | 1.335 | 6.699 | 1.093 | 1.264 |
| Dd2 PI4K S1320L+L1418F | Phosphatidylinositol 4-kinase | PF3D7_050 9800 | 1.277 | 0.183 | 0.710 | 0.909 |

| | | | | | | |
|---|---|---|---|---|---|---|
| Dd2 PI4K S743F+H1484Y | Phosphatidylinositol 4-kinase | PF3D7_1213800 | 0.850 | 0.345 | 0.520 | 0.791 |
| Dd2 ProRS L482H | Pro-tRNA synthetase | PF3D7_1011400 | 2.816 | 0.060 | 1.478 | 1.562 |
| Dd2 PROTB5 A20V | Proteasome beta 5 26S (A80V immature) | PF3D7_1011400 | 1.010 | 0.325 | 0.610 | 0.836 |
| Dd2 PROTB5 M45I | Proteasome beta 5 26S (M105I immature) | PF3D7_1359900 | 0.831 | 0.740 | 1.020 | 1.185 |
| Dd2 QRP1 D1863Y | Quinoxaline resistance protein | PF3D7_1117500 | 1.974 | 4.128 | 0.884 | 1.056 |
| Dd2 TyrRS S234C | Tyr-tRNA synthetase | PF3D7_1113300 | 1.580 | 0.840 | 0.872 | 0.620 |
| Dd2 UDP-GT F37V | UDP-galactose transporter | PF3D7_1211900 | 0.354 | 0.006 | 0.048 | 0.191 |
| Dd2 yDHODH | yeast dihydroorotate dehydrogenase | | 4.481 | 3.035 | 2.695 | 1.800 |

**Supplementary Table 6. List of oligonucleotides for gene knockdown donor vector construction.**

| Description | Nucleotide sequence |
|---|---|
| ThrRS cKD LHR forward | TGATGTTGAAGAAAATCCAGGTCCACACCAATGTGGAACCATAC AACTGG |
| ThrRS cKD LHR reverse | GGATGTAGTTGAATTGTTTTAATTGTGCTTCTCTTATTTTTTTGTT TAATGTATTGACGG |
| ThrRS cKD RHR forward | GTACAAACCCGGAATTCGAGCTCGGGCTTTAATGATGACATAAG TACATA |
| ThrRS cKD RHR reverse | TTAGACCTAGGGATAACAGGGTAATCATATAATTCTGAGGTACTT GAAGA |
| ThrRS sgRNA Klenow forward | TAACGGTCCTAAGGTAGCGAATAATACGACTCACTATAGGCAAC GAACACTGTAACTTTA |
| ThrRS sgRNA Klenow reverse | GACTAGCCTTATTTTAACTTGCTATTTCTAGCTCTAAAACTAAAG TTACAGTGTTCGTTG |
| ThrRS recodonized region | TTCAACTACATCCTGGTCGTGGGTGAAAAAGAGCTCACCACAA ACACTGTCACCCTTAGGGATAGAGACGATCAGAACAACCAACA CGTTTATACCATTCAAGAACTCATCAACAAGTTCAACAAGCTTC TCGACGTGAACTCTAAGAAGTTCAATCAGATTAAGGAATTCAAC TCGAACCAGACTATC |

**Supplementary Table 7. Percentage inhibition of ATP consumption by recombinant *Pf*AsnRS and *Hs*AsnRS induced by treatment with 100 µM OSM-S-106 or AMS (free base), or 1 µM Asn conjugates of OSM-S-106, LG420, LG866, and AMS.**

| **% Inhibition ± SEM** | **OSM-S-106** | **AMS** | **Asn-OSM106** | **Asn-LG420** | **Asn-LG866** | **Asn-AMS** |
|---|---|---|---|---|---|---|
| ***Pf*AsnRS** | 31.3 ± 1.6 | 107.9 ± 1.0 | 80.5 ± 1.7 | 87.9 ± 2.1 | 94.5 ± 1.1 | 101.7 ± 4.2 |
| ***Hs*AsnRS** | 0.6 ± 1.3 | 115.7 ± 2.7 | 36.0 ± 1.8 | 5.7 ± 3.2 | 23.2 ± 5.3 | 112.4 ± 1.0 |

**Supplementary Table 8. Thermal melting of recombinant *Pf*AsnRS in the presence and absence of low level SDS.** *Pf*AsnRS (3 µM) was incubated with 40 µM ATP, 80 µM L-asparagine, and 60 µM *Ec*tRNA in 25 mM Tris-HCl (pH 8.0), 150 mM NaCl, 5 mM $MgCl_2$, 1 mM TCEP, at 37°C for 3 h. $Tm_{app}$ was assessed in the presence of SYPRO Orange, with or without 0.001% SDS.

| | ***Pf*AsnRS (with or without 0.001% SDS)** | **$Tm_{app}$ (°C) Mean ± SEM** | **n** |
|---|---|---|---|
| **- SDS** | **Enzyme alone** | 60.0 ± 0.1 | 3 |
| | **+ ATP + Asn + *Ec*tRNA** | 57.2 ± 0.3 | 3 |
| **+ SDS** | **Enzyme alone** | 58.2 ± 0.1 | 7 |
| | **+ ATP + Asn + *Ec*tRNA** | 57.6 ± 0.2 | 7 |

**Supplementary Table 9. Thermal stabilization of recombinant *Pf*AsnRS and *Hs*AsnRS by nucleoside sulfamates.** *Pf*AsnRS or *Hs*AsnRS was incubated with 100 µM LG series free base or Asn-adduct, 40 µM ATP, 80 µM L-asparagine, and 60 µM *Ec*tRNA in 25 mM Tris-HCl (pH 8.0), 150 mM NaCl, 5 mM $MgCl_2$, 1 mM TCEP, at 37°C for 3 h. $Tm_{app}$ was assessed in the presence of 10X SYPRO Orange, with the addition of 0.001% SDS for *Pf*AsnRS, but not for *Hs*AsnRS. $\Delta Tm_{app}$ was calculated as $Tm_{app}$ (AsnRS + ATP + Asn + *Ec*tRNA + pro-inhibitor) − $Tm_{app}$ (AsnRS + ATP + Asn + *Ec*tRNA), or as $Tm_{app}$ (AsnRS + Asn-adduct) − $Tm_{app}$ (AsnRS alone).

| ***Pf*AsnRS** | **$Tm_{app}$ (°C) Mean ± SEM** | **Δ $Tm_{app}$ (°C) Mean ± SEM** | **n** |
|---|---|---|---|
| **Enzyme alone*** | 58.2 ± 0.1 | - | 7 |
| **+ ATP + Asn + *Ec*tRNA*** | 57.6 ± 0.2 | - | 7 |
| **+ ATP + Asn + *Ec*tRNA + 100 µM AMS** | 61.8 ± 0.1 | 4.5 ± 0.1 | 4 |
| **+ ATP + Asn + *Ec*tRNA + 100 µM OSM-S-106** | 59.0 ± 0.1 | 1.9 ± 0.1 | 3 |
| **+ ATP + Asn + *Ec*tRNA + 100 µM LG420** | 58.0 ± 0.2 | 0.8 ± 0.2 | 4 |
| **+ ATP + Asn + *Ec*tRNA + 100 µM LG866** | 58.7 ± 0.1 | 1.5 ± 0.1 | 3 |
| **+ ATP + Asn + *Ec*tRNA + 100 µM LG041** | 58.8 ± 0.1 | 1.6 ± 0.1 | 3 |
| **+ ATP + Asn + *Ec*tRNA + 100 µM LG685** | 59.3 ± 0.1 | 1.8 ± 0.1 | 4 |
| **+ ATP + Asn + *Ec*tRNA + 100 µM LG521** | 59.3 ± 0.1 | 1.8 ± 0.1 | 4 |
| **+ ATP + Asn + *Ec*tRNA + 100 µM LG563** | 58.3 ± 0.1 | 1.2 ± 0.1 | 3 |
| **+ ATP + Asn + *Ec*tRNA + 100 µM LG677** | 59.3 ± 0.1 | 2.1 ± 0.1 | 3 |
| **+ 100 µM Asn-AMS** | 66.0 ± 0.1 | 8.1 ± 0.1 | 3 |
| **+ 100 µM Asn-OSM-S-106** | 63.2 ± 0.0 | 5.3 ± 0.0 | 3 |
| **+ 100 µM Asn-LG420** | 63.7 ± 0.1 | 5.8 ± 0.1 | 3 |
| **+ 100 µM Asn-LG866** | 64.1 ± 0.1 | 6.0 ± 0.2 | 7 |

| ***Hs*AsnRS** | **$Tm_{app}$ (°C) Mean ± SEM** | **Δ $Tm_{app}$ (°C) Mean ± SEM** | **n** |
|---|---|---|---|
| **Enzyme alone** | 51.5 ± 0.1 | - | 3 |
| **+ ATP + Asn + *Ec*tRNA** | 50.8 ± 0.1 | - | 3 |
| **+ ATP + Asn + *Ec*tRNA + 100 µM AMS** | 68.6 ± 0.1 | 17.8 ± 0.2 | 3 |
| **+ ATP + Asn + *Ec*tRNA + 100 µM OSM-S-106** | 53.5 ± 0.2 | 2.7 ± 0.1 | 3 |
| **+ ATP + Asn + *Ec*tRNA + 100 µM LG420** | 50.5 ± 0.1 | -0.3 ± 0.0 | 3 |

| | | | |
|---|---|---|---|
| **+ ATP + Asn + *Ec*tRNA + 100 µM LG866** | 50.6 ± 0.1 | -0.2 ± 0.0 | 3 |
| **+ ATP + Asn + *Ec*tRNA + 100 µM LG041** | 50.4 ± 0.1 | -0.3 ± 0.0 | 3 |
| **+ ATP + Asn + *Ec*tRNA + 100 µM LG685** | 50.8 ± 0.1 | 0.0 ± 0.1 | 3 |
| **+ ATP + Asn + *Ec*tRNA + 100 µM LG521** | 50.7 ± 0.2 | 0.0 ± 0.1 | 3 |
| **+ ATP + Asn + *Ec*tRNA + 100 µM LG563** | 50.5 ± 0.1 | -0.2 ± 0.0 | 3 |
| **+ ATP + Asn + *Ec*tRNA + 100 µM LG677** | 50.5 ± 0.1 | -0.2 ± 0.0 | 3 |
| **+ 100 µM Asn-AMS** | 73.6 ± 0.1** | 22.0 ± 0.1** | 3 |
| **+ 100 µM Asn-OSM-S-106** | 63.5 ± 0.0 | 12.0 ± 0.1 | 3 |
| **+ 100 µM Asn-LG420** | 58.1 ± 0.1 | 6.6 ± 0.1 | 3 |
| **+ 100 µM Asn-LG866** | 62.1 ± 0.1 | 10.6 ± 0.1 | 3 |

* Data repeated from Supplementary Table 7, for ease of comparison.

**Main transition

**Chemistry Methods**

**General Experimental Details:**

Yields reported herein refer to purified products. Analytical TLC was performed on Merck silica gel 60 $F_{254}$ aluminium-backed plates. Compounds were visualised by UV light and/or stained with 5% $H_2SO_4$ in ethanol followed by heating. Flash column chromatography was performed on silica gel (100-200 Mesh). $^1$H-NMR spectra were recorded on a Bruker 400 MHz, Avance II spectrometer with a 5mm DUL (Dual) $^{13}$C probe and Bruker 400 MHz, Avance III HD spectrometer with BBFO (Broad Band Fluorine Observe) probe. Chemical shifts ($\delta$) are expressed in parts per million (ppm) with reference to the deuterated solvent peak in which the sample is prepared.

Abbreviations used:

| | |
|---|---|
| TLC | Thin Layer Chromatography |
| mL | Milliliters |
| mmol | Millimoles |
| h | Hour or hours |
| min | Minute or minutes |
| g | Grams |
| mg | Milligrams |
| rt or RT | Room temperature, ambient, about 25°C |
| MeOH | Methanol |
| EA | Ethyl acetate |
| RM | Reaction mixture |
| THF | Tetrahydrofuran |
| KOAc | Potassium acetate |

*Preparation of LG-0020420 (**3**):*
A mixture of 3-(4,4,5,5-tetramethyl-1,3,2-dioxaborolan-2-yl)benzenesulfonamide (120 mg, 424 µmol), 7-bromoquinazolin-4-amine (100 mg, 446 µmol), Pd(dppf)$Cl_2$ (30.0 mg, 41.0 µmol) and $K_2CO_3$ (180 mg, 1.30 mmol) in dioxane (1.5 mL) and $H_2O$ (0.4 mL) was degassed and purged with $N_2$ for 3 times, and then the mixture was stirred at 80 °C for 12 h under $N_2$ atmosphere. The mixture was filtered and the solid was collected, the filter cake was washed with water (4 mL) and MeOH (3 mL). Then triturated with DMF (added 2 drops TFA) to give 3-(4-aminoquinazolin-7-yl)benzenesulfonamide (38.8 mg, 129 µmol, 28.9% yield) as a pale yellow solid.$^1$H NMR (400 MHz, DMSO-$d_6$) δ 10.03 - 9.68 (m, 2H), 8.86 (s, 1H), 8.63 (d, $J$ = 8.8 Hz, 1H), 8.25 (s, 1H), 8.18 - 8.12 (m, 1H), 8.11 - 8.04 (m, 2H), 7.97 (d, $J$ = 8.0 Hz, 1H), 7.84 - 7.76 (m, 1H), 7.52 (s, 2H); *m/z* ES+ $[M+H]^+$ 301.1.

*Preparation of LG-0020866 (**4**):*
Step 1: Synthesis of 7-(4,4,5,5-tetramethyl-1,3,2-dioxaborolan-2-yl)quinazolin-4-amine
To a stirred solution of 7-bromo-4-quinazolinylamine (0.3 g, 1.34 mmol) and 4,4,5,5-tetramethyl-2-(4,4,5,5-tetramethyl-1,3,2-dioxaborolan-2-yl)-1,3,2-dioxaborolane ($B_2Pin_2$) (374 mg, 1.1 eq., 1.47 mmol) in 1,4-dioxane (6 mL, 70.3 mmol) under $N_2$ gas was added potassium acetate (394 mg, 3 eq., 4.02 mmol) then purge with $N_2$ gas for 5 min. After that PdCl2(dppf).dcm (98 mg, 0.1 eq., 134 µmol) was added at RT. The resulting reaction

mixture was stirred at 100 $^0$C for 3 h. The reaction mixture was monitored by TLC and LCMS. Upon completion, the reaction mixture was quenched with water (20 mL) and extracted with dichloromethane (2 × 50 mL). The combined organic layers were washed with brine, dried over anhydrous sodium sulfate, filtered, and concentrated under reduced pressure to afford the crude title compound as an off-white solid (0.280 g, 77%). Crude compound was forwarded next step without purification.

Step 2: Synthesis of 6-(4-aminoquinazolin-7-yl)pyridine-2-sulfonamide (2)

To a stirred solution of 7-(4,4,5,5-tetramethyl-1,3,2-dioxaborolan-2-yl)-4-quinazolinylamine (280 mg, 1.03 mmol) and 6-bromo-2-pyridinesulfonamide (245 mg, 1.03 mmol) in 1,4-dioxane (5 mL, 58.6 mmol) & water (0.5 mL, 27.8 mmol) under $N_2$ gas was added dipotassium carbonate (357 mg, 2.5 eq., 2.58 mmol) then purge with $N_2$ gas for 5 min and then $PdCl_2$(dppf).dcm (84.3 mg, 0.1 eq., 103 µmol) were added at RT. The resulting reaction mixture was stirred at 100 $^0$C for 16 h. The reaction mixture was monitored by TLC and LCMS. Upon completion, the reaction mixture was quenched with water (20 mL) and extracted with dichloromethane (2 × 50 mL). The combined organic layers were washed with brine, dried over anhydrous sodium sulfate, filtered, and concentrated under reduced pressure to afford the crude product which was further purified by prep-HPLC in ABC buffer to afford the title compound as an off-white solid (0.026 g, 8%). $^1$H NMR (400 MHz, DMSO-*d6*): $\delta$ 8.52 (s, 1H), 8.45-8.41 (m, 2H), 8.36 (s, 2H), 8.20 (t, *J*=7.8 Hz, 1H), 7.95-7.87 (m, 3H), 7.6 (s, 2H). LC/MS: 99%. MS (ESI$^+$): *m/z* 301.95 (M+H), (RT: 3.78 min and Purity 97.44%).

*Preparation of LG-0021041 (**5**):*

Step 1: Synthesis of 5-bromo-2-fluorobenzenesulfonamide: To a stirred solution of 4-bromo-2-(chlorosulfonyl)-1-fluorobenzene 1 (2 g, 7.31 mmol) in Aqueous ammonia solution (40 mL) at 0°C. The reaction mixture was stirred at RT for 3h. The reaction mixture was monitored by TLC and LCMS. Upon completion, the reaction mixture was quenched with water (50 mL) and extracted with dichloromethane (2 × 100 mL). The combined organic layers were washed with brine, dried over anhydrous sodium sulfate, filtered, and concentrated under reduced pressure to afford the crude title compound as an off-white solid (1.2 g, 61%). LC/MS: MS (ESI): *m/z* 251.60 $[M-H]^+$, (retention time:0.98 min; purity 94.49%).

Step-2: Synthesis of 2-fluoro-5-(4, 4, 5, 5-tetramethyl-1, 3, 2-dioxaborolan-2-yl) benzenesulfonamide: To a stirred solution of 5-bromo-2-fluorobenzenesulfonamide 2 (0.1 g, 394 µmol) in 1,4-dioxane (3 mL) and $B_2Pin_2$ (150 mg, 1.5 eq., 590 µmol) was added potassium acetate (77.3 mg, 2 eq., 787 µmol) at RT. The reaction mixture was degassed with nitrogen for 10 minutes. After that mixture was charged with [1, 1′-Bis (diphenylphosphino) ferrocene] dichloropalladium (II), DCM complex with (32.1 mg, 0.1 eq., 39.4 µmol) at RT. The reaction mixture was heated at 100 °C for 3 hours.

The reaction mixture was monitored by TLC and LCMS. Upon completion, the reaction mixture was quenched with water (20 mL) and extracted with ethyl acetate (2 × 50 mL). The combined organic layers were washed with brine, dried over anhydrous sodium sulfate, filtered, and concentrated under reduced pressure to afford the title crude compound as a brown solid, used further without purification. (0.220 g, 19%). LC/MS: MS (ESI): *m/z* 415.05 $[M+H]^+$,(retention time: 4.83 min; purity 99.68%).

Step-3: Synthesis of 5-(4-amino-7-quinazolinyl)-2-fluorobenzenesulfonamide: To a stirred solution of 2-fluoro-5-(4,4,5,5-tetramethyl-1,3,2-dioxaborolan-2-yl) benzenesulfonamide 3 (220 mg, 731 µmol) and 7-bromo-4-quinazolinylamine (164 mg, 731 µmol) in in 1,4-dioxane (8 mL) and water (2 mL) was added dipotassium carbonate (252 mg, 2.5 eq., 1.83 mmol) at RT. The reaction mixture was degassed with nitrogen for 10 minutes. To the reaction mixture charged [1,1′-Bis(diphenylphosphino)ferrocene]dichloropalladium(II), complex with dichloromethane (119 mg, 0.2 eq., 146 µmol) at RT. The reaction mixture was heated at 100 °C for 3 hours. The

reaction was monitored by TLC and LCMS. Upon completion, the reaction mixture was filtered through scintered funnel, washed with dichloromethane, and the filtrate was concentrated under reduced pressure to get crude product which was further purified by prep-HPLC in ABC buffer to afford title compound as white solid (0.087 g, 37%). $^1$H NMR (400 MHz, DMSO-*d*6): δ 8.44 (s, 1H), 8.35 (d, *J*= 8.64 Hz, 1H), 8.18-8.12 (m, 2H), 7.93 (d, *J*=1.6 Hz, 1H), 7.84-7.82 (m, 3H), 7.59 (t, *J* = 8.72 Hz, 1H).
LCMS: MS (ESI$^+$): *m/z* 319.3 (M+H), (RT: 3.16 min and Purity 99.80%).

*Preparation of LG-0021685 (**6**):*
Step 1: Synthesis of 6-bromo-3-fluoro-2-[(p-methoxyphenyl)methylthio]pyridine: To a stirred solution of (p-methoxyphenyl)methanethiol (0.4 g, 2.59 mmol) in tetrahydrofuran (10 mL, 123 mmol) was added potasium tert butoxide (582 mg, 2 eq., 5.19 mmol) at 0°C. The resultant reaction mixture was stirred at 0°C for 10 min, followed by the addition of 6-bromo-2-chloro-3-fluoropyridine (546 mg, 2.59 mmol). The resultant reaction mixture was stirred at RT for 3 h. The reaction mixture was monitored by TLC and LCMS. Upon completion, the reaction mixture was quenched with water (50 mL) and extracted with ethyl acetate (2 × 100 mL). The combined organic layers were washed with brine, dried over anhydrous sodium sulfate, filtered, and concentrated under reduced pressure to afford the crude product. The crude material was purified by flash column chromatography using a 12 g snap column, eluting with 0–5% ethyl acetate in heptane, to afford the title compound as an off-white solid (0.26 g, 26%). LC/MS: MS (ESI): *m/z* 327.60 [M+H]$^+$, (retention time: 2.34 min; purity 86.35%).
Step 2: Synthesis of 6-bromo-2-(chlorosulfonyl)-3-fluoropyridine: To a stirred solution of 6-bromo-3-fluoro-2-[(p-methoxyphenyl)methylthio]pyridine (0.5 g, 1.52 mmol) in dichloromethane (22 mL, 343 mmol) and water (1.1 mL, 60.9 mmol) was added acetic acid (1.1 mL, 13 eq., 19.2 mmol) under inert atmosphere at -10°C. Then 1,3-dichloro-5,5-dimethyl-2,4-imidazolidinedione (0.6 g, 2 eq., 3.05 mmol) was added and resulting reaction mixture was stirred at -10°C for 1 h. The reaction mixture was monitored by TLC and LCMS. Upon completion, the reaction mixture was quenched with $NaHCO_3$ (50 mL) and extracted with dichloromethane (2 × 100 mL). The combined organic layers were washed with brine, dried over anhydrous sodium sulfate, filtered, and concentrated under reduced pressure to afford the crude title compound as a brown solid (0.41 g, 98%) and forwarded to the next step without purification.
Step 3: Synthesis of 6-bromo-3-fluoro-2-pyridinesulfonamide: Procedure: To a 100 mL RBF charged with 6-bromo-2-(chlorosulfonyl)-3-fluoropyridine (410 mg, 1.49 mmol) was added NH3 (7 M in MeOH) (8.54 mL, 40 eq., 59.7 mmol) at 0°C and under inert atmosphere. After addition, the reaction was stirred at the room temperature for 3 h. The reaction mixture was monitored by TLC and LCMS. Upon completion, the reaction mixture was quenched with water (50 mL) and extracted with dichloromethane (2 × 100 mL). The combined organic layers were washed with brine, dried over anhydrous sodium sulfate, filtered, and concentrated under reduced pressure to afford the crude product which was purified by flash column chromatography using a 12 g snap column, eluting with 30–40% ethyl acetate in heptane to afford the title compound as an off-white solid (0.3 g, 36%). LC/MS: MS (ESI): *m/z* 252.60 [M-H]$^+$, (retention time: 0.85 min; purity 45.82%).

Step 4: Synthesis of 6-(4-amino-7-quinazolinyl)-3-fluoro-2-pyridinesulfonamide—trifluoroacetic acid: To a stirred solution of 6-bromo-3-fluoro-2-pyridinesulfonamide (0.3 g, 1.18 mmol) and 7-(4,4,5,5-tetramethyl-1,3,2-dioxaborolan-2-yl)-4-quinazolinylamine (1.38 g, 1.2 eq., 5.09 mmol) in 1,4-dioxane (8 mL, 93.8 mmol) and water (2 mL, 111 mmol) was added $Cs_2CO_3$ (958 mg, 2.5 eq., 2.94 mmol) under inert atmosphere. Reaction mixture was purged with N2 for 5 min. Then CatCxium Pd G3 (85.6 mg, 0.1 eq., 118 μmol) was added and again purged with N2 for 5 min. Resulting reaction mixture was stirred at 100°C for 16 h. The reaction mixture was monitored by TLC and LCMS. Upon completion, the reaction mixture was quenched with water (50 mL) and extracted with dichloromethane (2 × 100 mL). The combined organic layers were washed with brine, dried over anhydrous sodium sulfate, filtered, and concentrated under reduced pressure to afford the crude product which was was purified by prep-HPLC in TFA buffer to afford the title compound as an off-white solid (0.023 g, 4%). $^1$H NMR (400 MHz, DMSO-*d6*): δ 9.55 (s, 1H), 8.90 (s, 1H), 8.55 (m, 3H), 8.47 (s, 1H), 8.22 (t, *J* = 9.08 Hz, 1H), 7.98 (s, 2H). LC/MS: MS (ESI$^+$): *m/z* 320.02 (M+H), (RT: 2.97 min and Purity 99.72%).

*Preparation of LG-0021877 (7):*

Step 1: Synthesis of 7-bromo-5-fluoro-4-quinazolinylamine: To a stirred solution of 4-bromo-2,6-difluorobenzonitrile (2 g, 9.17 mmol) in dimethylacetamide (10 mL, 108 mmol) in a 30 mL glass vial at room temperature was added acetic acid.formamidine (1.91 g, 2 eq., 18.3 mmol) and the reaction was stirred at 150 °C for 10 h. The reaction mixture was monitored by TLC and LCMS. After completion, the reaction mixture was evaporated under reduced pressure to get the crude product which was allowed to suspend in cold water (20 mL) and pH was adjusted to 8.5 by adding aqueous $NH_4OH$ solution. The reaction mass was cooled in ice-bath for 1 h, a brown precipitate was obtained, filtered and washed with chilled water, diethyl ether and finally with pentane. The crude compound was purified by flash column chromatography using a 12 g snap column, eluting with 30–40% ethyl acetate in heptane, to afford the title compound (750 mg, 2.51 mmol, 27%) as an off-white solid.
LC/MS: MS (ESI): *m/z* 244.06 [M+H]$^+$, (retention time: 1.54 min; purity 81.68%).

Step 2: Synthesis of 5-fluoro-7-(4,4,5,5-tetramethyl-1,3,2-dioxaborolan-2-yl)-4-quinazolinylamine: To a stirred solution of 7-bromo-5-fluoro-4-quinazolinylamine (750 mg, 3.1 mmol) and 4,4,5,5-tetramethyl-2-(4,4,5,5-tetramethyl-1,3,2-dioxaborolan-2-yl)-1,3,2-dioxaborolane (1.18 g, 1.5 eq., 4.65 mmol) in 1,4-dioxane (26.7 mL, 313 mmol) in a 30 mL glass vial at room temperature was added KOAc (912 mg, 3 eq., 9.3 mmol) under $N_2$ atmosphere. The reaction mixture was purged with $N_2$ gas for 5 min and then added PdCl2(dppf).dcm (227 mg, 0.1 eq., 310 μmol). The resulting reaction was again purged with $N_2$ gas for 5 min. The reaction was stirred at 85°C for 4 h. The reaction mixture was monitored by TLC and LCMS. Upon completion, the reaction mixture was quenched with water (50 mL) and extracted with dichloromethane (2 × 100 mL). The combined organic layers were washed with brine, dried over anhydrous sodium sulfate, filtered, and concentrated under reduced pressure to afford the crude title compound (0.8 g, 2.77 mmol, 89%) as brown semi solid. Crude compound was forwarded to next step without further purification. LCMS data complied with boronic acid mass.
LC/MS: MS (ESI): *m/z* 208.14 [M+H]$^+$, (retention time: 0.21 and 0.49 min; purity (89.58%)

Step 3: Synthesis of m-(4-amino-5-fluoro-7-quinazolinyl)benzenesulfonamide—trifluoroacetic acid (1/1): To a stirred solution of m-bromobenzenesulfonamide **5** (0.5 g, 2.12 mmol) and 5-fluoro-7-(4,4,5,5-tetramethyl-1,3,2-dioxaborolan-2-yl)-4-quinazolinylamine (735 mg, 1.2 eq., 2.54 mmol) in a mixture of 1,4-dioxane (8 mL, 93.8 mmol) and water (2 mL, 111 mmol) in a 30 mL reaction vial was added cesium carbonate (1.73 g, 2.5 eq., 5.29 mmol) under inert atmosphere at room temperature. The reaction mixture was purged with $N_2$ gas for 5 min and then added cataCXium® A Pd G3 (154 mg, 0.1 eq., 212 μmol). Reaction mixture was again purged with $N_2$ gas for 5 min. The resulting reaction mixture was stirred at 100°C for 3 h. The progress of reaction was monitored by TLC and LCMS. After completion of reaction, the reaction mixture was quenched with water (50 mL and extracted with ethyl acetate (3 x 30 mL). The combined organic layers were washed with brine solution and dried over anhydrous sodium sulfate, filtered and distilled over rotavapor to get crude compound. The crude compound was purified from prep-HPLC in TFA buffer to obtain m-(4-amino-5-fluoro-7-quinazolinyl)benzenesulfonamide.trifluoroacetic acid (22 mg, 49.4 μmol, 2%) as an off white solid. $^1$H NMR (400 MHz, DMSO-$d_6$): *δ* 9.66 (s, 1H), 8.85 (s, 2H), 8.25 (s, 1H), 8.10 (d, *J* = 7.96 Hz, 1H), 7.97-7.95 (m, 2H)

7.85 (d, *J* = 1.04 Hz, 1H), 7.78 (t, *J* = 7.80 Hz, 1H), 7.50 (s, 2H). LC/MS: MS (ESI): *m/z* 319.17 $[M+H]^+$, (retention time: 4.07 min; purity (95.68%).

*Preparation of LG-0022521 (**8**):*

Step 1: Synthesis of 7-bromo-8-fluoro-4-quinazolinylamine: To a stirred solution of 4-bromo-2,3-difluorobenzonitrile (1 g, 1.0 eq, 4.59 mmol) in 5 mL dimethylacetamide_at RT was added acetic acid/formamidine (1/1) (955 mg, 2 eq., 9.17 mmol) and stirred at 150 °C for 16 h. After completion of the reaction, the reaction mixture was evaporated under reduced pressure to afford a dry brown solid. The obtained residue was suspended in cold water, and the pH was adjusted to 8.5 using aqueous ammonium hydroxide. The mixture was further cooled and stirred in water for 1 h, resulting in the formation of a brown precipitate. The precipitate was collected by filtration and washed sequentially with chilled water, diethyl ether, and pentane to afford title compound (0.6 g, 46%) as a yellow powder.

LC/MS**:** MS ($ESI^+$): *m/z* 241.70 $[M+H]^+$, (RT: 1.37 min and Purity 85.10%).

Step 2: Synthesis of 3-(4-amino-8-fluoroquinazolin-7-yl)benzenesulfonamide:

To a stirred solution of 7-bromo-8-fluoro-4-quinazolinylamine (0.4 g, 1.0 eq, 1.65 mmol) and m-(4,4,5,5-tetramethyl-1,3,2-dioxaborolan-2-yl)benzenesulfonamide ( 468 mg, 1.0 eq, 1.65 mmol) in 15 mL 1,4-dioxane and 1.5 mL water were added cesium carbonate (1.35 g, 2.5 eq., 4.13 mmol) under inert atmosphere at RT. Reaction mixture was purged with N2 for 5 min. Then cataCXium® A Pd $G_3$ (124 mg, 0.1 eq., 165 µmol) was added. The reaction mixture was again purged with N2 for 5 min, the resulting reaction mixture was stirred at 100°C for 16 h. The reaction mixture was monitored by TLC and LCMS. Upon completion, the reaction mixture was quenched with water (50 mL) and extracted with dichloromethane (2 × 50 mL). The combined organic layers were washed with brine, dried over anhydrous sodium sulfate, filtered, and concentrated under reduced pressure to afford the crude product. The crude material was purified by flash column chromatography using a 12 g snap column, eluting with 15–20% MeOH in DCM to afford the title compound as an off white solid (0.03 g, 5%). $^1$H NMR (400 MHz, DMSO-d6 ): δ 8.51 (s, 1H), 8.37 (brs, 1H), 8.20 (d, *J*= 8.64 Hz, 2H), 8.12 (s, 1H), 7.93-7.91 (m, 2H), 7.75 (t, *J*=7.8 Hz, 1H), 7.67 (t, *J*=7.64 Hz, 1H) 7.48 (s, 2H). LC/MS**:** MS ($ESI^+$): *m/z* 319.23 $[M+H]^+$, (RT: 3.54 min and Purity 98.39%).

P*reparation of LG-0022563 (**9**):*

Step 1 : Synthesis of 7-bromo-5-fluoroquinazolin-4-amine: To a stirred solution of 4-bromo-2,6-difluorobenzonitrile (2 g, 1.0 eq, 9.17 mmol) in 10 mL dimethylacetamide at RT was added formamidine acetate (1.91 g, 2 eq., 18.3 mmol) and stirred at 150 °C for 10 h. After completion, the reaction mixture was evaporated under reduced pressure to dryness to afford a brown solid. The obtained residue was suspended in cold water, and the pH was adjusted to 8.5 using aqueous ammonium hydroxide. The mixture was further cooled and stirred in water for 1 h, resulting in the formation of a brown precipitate. The precipitate was collected by filtration and washed sequentially with chilled water, diethyl ether, and finally with pentane. The crude material was further purified by flash column chromatography using a 12 g snap column, eluting with ethyl acetate and heptane. The desired product eluted at 30–40% ethyl acetate in heptane. to afford title compound (3) (820 mg, 31%) as a brown solid. LC/MS: MS (ESI): *m/z* 242.0 $[M+H]^+$, (retention time: 1.71 min; purity 84.11%).

Step 2: Synthesis of (4-amino-5-fluoroquinazolin-7-yl)boronic acid: To a stirred solution of 7-bromo-5-fluoroquinazolin-4-amine (0.3 g, 1.0 eq, 1.24 mmol) and 4,4,5,5-tetramethyl-2-(4,4,5,5-tetramethyl-1,3,2-dioxaborolan-2-yl)-1,3,2-dioxaborolane (378 mg, 1.2 eq., 1.49 mmol) in 20 mL 1,4-dioxane under N2 gas was

added KOAc (365 mg, 3 eq., 3.72 mmol) then purge the N2 gas for 5 min at RT. After that PdCl2(dppf).dcm (90.7 mg, 0.1 eq., 124 µmol) was added and the resulting reaction mixture was stirred at 110 °C for 3 h.
The reaction mixture was monitored by TLC and LCMS. Upon completion, the reaction mixture was quenched with water (50 mL) and extracted with ethyl acetate (2 × 100 mL). The combined organic layers were washed with brine, dried over anhydrous sodium sulfate, filtered, and concentrated under reduced pressure to afford the crude title compound as brown semi-solid (0.27 g, 44%). LC/MS: MS (ESI): *m/z* 415.05 $[M+H]^+$, (retention time: 4.83 min; purity 99.68%).
Step 3: Synthesis of 6-(4-amino-5-fluoroquinazolin-7-yl)pyridine-2-sulfonamide: To a stirred solution of 4-amino-5-fluoroquinazolin-7-yl)boronic acid (268 mg, 928 µmol) and 6-bromo-2-pyridinesulfonamide (220 mg, 1.0 eq, 928 µmol) in 9 mL 1,4-dioxane and 1 mL water were added cesium carbonate (756 mg, 2.5 eq., 2.32 mmol) under inert atmosphere at room temperature. The reaction mixture was purged with N2 for 5 min. Then cataCXium® A Pd G3 (69.7 mg, 0.1 eq., 92.8 µmol) was added and again purged with $N_2$ atmosphere for 5 min at room temperature. The resulting reaction mixture was stirred at 100°C for 16 h. The progress of the reaction was monitored by TLC and LCMS. Upon completion, the reaction mixture was quenched with water (20 mL), resulting in the formation of a brown precipitate. The precipitate was collected by filtration and washed sequentially with chilled water, diethyl ether, and finally with pentane. The solid was dried under vacuum to afford the crude material which was purified by preparative HPLC using ABC buffer to afford the title compound as an off-white solid (33 mg, 11%). $^1$H NMR (400 MHz, DMSO-*d6*): δ 8.47-8.49 (m, 2H), 8.39 (d, *J*= 1.2 Hz, 1H), 8.23-8.19 (m, 2H), 8.10 (brs, 1H), 7.96 (d, *J*= 7.6 Hz, 1H), 7.63 (s, 1H), 7.40 (brs, 1H). LC/MS: MS ($ESI^+$): *m/z* 320.18 $[M+H]^+$, (RT: 3.42 min and Purity 97.56%).

*Preparation of LG-0022677 (**10**):*
Step 1A: Synthesis of 6-bromo-3-fluoro-2-((4-methoxybenzyl)thio)pyridine: To a stirred solution of (4-methoxyphenyl)methanethiol (1.2 g, 1.0 eq, 7.78 mmol) in 20 mL tetrahydrofuran was added pottasium tert butoxide (1.75 g, 2 eq., 15.6 mmol) at 0°C. The resultant reaction mixture was stirred at 0°C for 10 min, followed by 6-bromo-2-chloro-3-fluoropyridine (1.47 g, 0.9 eq., 7 mmol) in reaction mixture. The resultant reaction mixture was stirred at RT for 3 h. The reaction mixture was monitored by TLC and LCMS. Upon completion, the reaction mixture was quenched with water (50 mL) and extracted with ethyl acetate (2 × 100 mL). The combined organic layers were washed with brine, dried over anhydrous sodium sulfate, filtered, and concentrated under reduced pressure to afford the crude product. The crude material was purified by flash column chromatography using a 12 g snap column, eluting with 10–20% ethyl acetate in heptane to afford the title compound as colorless oil (0.65 g, 38%). LC/MS: MS (ESI): *m/z* 325.65 $[M-H]^+$, (retention time: 2.43 min; purity 76.35%).
Step 2A: Synthesis of 6-bromo-3-fluoropyridine-2-sulfonyl chloride: To a stirred solution of 6-bromo-3-fluoro-2-((4-methoxybenzyl)thio)pyridine (650 mg, 1.0 eq, 1.98 mmol) in 20 mL dichloromethane and 1.4 mL water was added 1.4 mL acetic acid under inert atmosphere at -10°C. Then 1,3-dichloro-5,5-dimethyl-2,4-imidazolidinedione (780 mg, 2 eq., 3.96 mmol) was added and resulting reaction mixture was stirred at -10°C for 1 h. The reaction mixture was monitored by TLC and LCMS. Upon completion, the reaction mixture was quenched with $NaHCO_3$ (30 mL) and extracted with dichloromethane (2 × 60 mL). The combined organic layers were washed with brine, dried over anhydrous sodium sulfate, filtered, and concentrated under reduced pressure to afford the crude title compound as an brown solid 0.53 g, 97%) and forward to next step without purification.
Step 3A: Synthesis of 6-bromo-3-fluoropyridine-2-sulfonamide: To a stirred solution of 6-bromo-3-fluoropyridine-2-sulfonyl chloride (530 mg, 1.0 eq, 1.93 mmol) in 20 mL of 7.0 M NH3 in methanol was stirred at RT for 2 h. The progress of reaction was monitored by TLC. After completion, reaction mixture was directly

concentrated under reduced pressure to get crude. The crude was purified by Si-gel coloumn chromatography using 30% EtOAc in Heptane as elutant. Appropriate column fractions were concentrated under vacuum to get the desired 6-bromo-3-fluoropyridine-2-sulfonamide (130 mg, 18%) as off white solid.
LC/MS: MS (ESI): *m/z* 252.90 $[M\text{-}H]^+$, (retention time: 0.92 min; purity 68.46%).
Step 1 : Synthesis of 7-bromo-5-fluoroquinazolin-4-amine: To a stirred solution of 4-bromo-2,6-difluorobenzonitrile (2 g, 1.0 eq, 9.17 mmol) in 10 mL dimethylacetamide at RT was added formamidine acetate (1.91 g, 2 eq., 18.3 mmol) and stirred at 150 °C for 10 h. After completion, the reaction mixture was evaporated under reduced pressure up to dryness and brown solid was obtained. The brown residual product was allowed to suspend in cold water and pH was adjusted up to 8.5 using NH4OH. Again, cooled the product for 1 h in H2O, a brown precipitate obtained which was filtered and washed with chilled H2O, diethyl ether and finally with pentane. The crude product was further purified by Si-gel column chromatography using 30-40% EtOAc in heptane as elutant. Appropriate column fractions were concentrated under vacuum to get 7-bromo-5-fluoroquinazolin-4-amine (1.2 g, 38%) as brown solid. LC/MS: MS (ESI): *m/z* 242.05 $[M+H]^+$, (retention time: 1.51 min; purity 69.68%).
Step 2: Synthesis of (4-amino-5-fluoroquinazolin-7-yl)boronic acid: To a stirred solution of 7-bromo-5-fluoroquinazolin-4-amine (0.7 g, 1.0 eq, 2.89 mmol) and 4,4,5,5-tetramethyl-2-(4,4,5,5-tetramethyl-1,3,2-dioxaborolan-2-yl)-1,3,2-dioxaborolane (881 mg, 1.2 eq., 3.47 mmol) in 20 mL 1,4-dioxane under N2 gas was added KOAc (851 mg, 3 eq., 8.68 mmol) then purge the N2 gas for 5 min at RT. After that PdCl2(dppf).dcm (212 mg, 0.1 eq., 289 µmol) was added and the resulting reaction mixture was stirred at 110 °C for 3 h. The reaction mixture was monitored by TLC and LCMS. Upon completion, the reaction mixture was quenched with water (50 mL) and extracted with dichloromethane (2 × 100 mL). The combined organic layers were washed with brine, dried over anhydrous sodium sulfate, filtered, and concentrated under reduced pressure to afford the crude title compound as brown semi-solid (0.31 g, 64%) and was forwarded next step without further purification.
Step 3: Synthesis of 6-(4-amino-5-fluoroquinazolin-7-yl)-3-fluoropyridine-2-sulfonamide trifluoroacetic acid**:** To a stirred solution of (4-amino-5-fluoroquinazolin-7-yl)boronic acid (147 mg, 1.0 eq, 510 µmol) and 6-bromo-3-fluoro-2-pyridinesulfonamide (130 mg, 1.0 eq, 510 µmol) in 4.5 mL 1,4-dioxane and 0.5 mL water were added cesium carbonate (415 mg, 2.5 eq., 1.27 mmol) under inert atmosphere at RT. Reaction mixture was purged with N2 for 5 min. Then cataCXium® A Pd G3 (38.3 mg, 0.1 eq., 51.5 µmol) was added. Reaction mixture was again purged with N2 for 5 min. Resulting reaction mixture was stirred at 100°C for 16 h. The progress of reaction was monitored by TLC and LCMS analysis. After completion of reaction, the reaction mixture was quenched with water (50 mL), a brown precipitate obtained which was filtered and washed with chilled H2O, diethyl ether and finally with pentane to get the crude compound which was further purified by prep-HPLC in TFA buffer to get to afford the title compound as an off white solid (14.5 mg, 6%). $^1$H NMR (400 MHz, DMSO-*d6*):$\delta$ 9.27 (brs, 1H), 8.68 (s, 1H), 8.55 (dd, *J1*= 8.8 Hz, *J2*= 3.2 Hz, 1H), 8.35-8.32 (m, 2H), 8.22 (t, *J*= 9.2 Hz, 1H), 7.99 (s, 2H).
LC/MS: MS ($ESI^+$): *m/z* 338.17 $[M+H]^+$, (retention time: 3.48 min and Purity 97.97%).

Preparation of the Asn adduct of LG-0020420:
Step-1: Synthesis of tert-butyl (7-bromoquinazolin-4-yl)(tert-butoxycarbonyl)carbamate: To a stirred solution of 7-bromo-4-quinazolinylamine (2 g, 8.93 mmol) in dichloromethane (20 mL, 312 mmol) in a 50 mL glass vial were added triethylamine (3.76 mL, 3 eq., 26.8 mmol), DMAP (218 mg, 0.2 eq., 1.79 mmol), followed by dropwise addition of di-tert-butyl dicarbonate (6.15 mL, 3 eq., 26.8 mmol) at 0°C under $N_2$ atmosphere. The resulting reaction mixture was stirred at room temperature for 16 h. The reaction mixture was monitored by TLC and LCMS. Upon completion, the reaction mixture was quenched with water (20 mL) and extracted with

dichloromethane (2 × 50 mL). The combined organic layers were washed with brine, dried over anhydrous sodium sulfate, filtered, and concentrated under reduced pressure to afford the crude product. The crude material was purified by flash column chromatography using a 12 g snap column, eluting with 20–30% ethyl acetate in heptane, to afford the title compound as an off-white solid (1.2 g, 2.63 mmol, 30%). LC/MS: MS (ESI): *m/z* 423.85.05 $[M+H]^+$, (retention time: 2.37 min; purity 93.41%).

Step-2: Synthesis of tert-butyl (tert-butoxycarbonyl)(7-(3-sulfamoylphenyl)quinazolin-4-yl)carbamate**:** To a stirred solution of 7-bromo-4-[tert-butoxycarbonyl-tert-butyl(oxycarbonylamino)]quinazoline (1.2 g, 2.83 mmol) and m-(4,4,5,5-tetramethyl-1,3,2-dioxaborolan-2-yl)benzenesulfonamide (881 mg, 1.1 eq., 3.11 mmol) in a 30 mL reaction vial in 1,4-dioxane (15 mL, 176 mmol) & water (1.5 mL, 83.3 mmol) was added dipotassium carbonate (977 mg, 2.5 eq., 7.07 mmol) under $N_2$ atmosphere at room temperature. The reaction mixture was purgged with $N_2$ gas for 5 min and then added PdCl2(dppf).DCM (231 mg, 0.1 eq., 283 µmol). The resulting reaction mixture was again purgged with $N_2$ gas for 15 min., and then stirred at 100 °C for 3 h.

The reaction mixture was monitored by TLC and LCMS. Upon completion, the reaction mixture was quenched with water (20 mL) and extracted with dichloromethane (2 × 50 mL). The combined organic layers were washed with brine, dried over anhydrous sodium sulfate, filtered, and concentrated under reduced pressure to afford the crude product. The crude material was purified by flash column chromatography using a 12 g snap column, eluting with 15–20% ethyl acetate in heptane, to afford the title compound as an off-white solid (1.0 g, 1.5 mmol, 53%). LC/MS: MS (ESI): *m/z* 501.15 $[M+H]^+$, (retention time: 2.29 min; purity 98.28%).

Step-3: Synthesis of tert-butyl (tert-butoxycarbonyl)(7-(3-(N-(N2-(tert-butoxycarbonyl)-N4-trityl-L-asparaginyl)sulfamoyl)phenyl)quinazolin-4-yl)carbamate: To a stirred solution of (*S*)-2-[tert-butyl(oxycarbonylamino)]-3-(N-tritylcarbamoyl)propionic acid (1.71 g, 1.5 eq., 3.6 mmol) in a 50 mL RBF in dichloromethane (15 mL, 234 mmol) under $N_2$ atmosphere were added EDC.HCl (689 mg, 1.5 eq., 3.6 mmol) & HOAt (489 mg, 1.5 eq., 3.6 mmol) at RT and stirred for 30 min. After 30 min., added *m*-{4-[tert-butoxycarbonyl-tert-butyl(oxycarbonylamino)]-7-quinazolinyl}benzenesulfonamide (1.2 g, 2.4 mmol) and DBU (1.07 mL, 3 eq., 7.19 mmol). The resulting reaction mixture was stirred at room temperature for 16 h. The reaction mixture was monitored by TLC and LCMS. Upon completion, the reaction mixture was quenched with water (20 mL) and extracted with dichloromethane (2 × 50 mL). The combined organic layers were washed with brine, dried over anhydrous sodium sulfate, filtered, and concentrated under reduced pressure to afford the crude product. The crude material was purified by flash column chromatography using a 12 g snap column, eluting with 60–70% ethyl acetate in heptane, to afford the title compound as an off-white solid (1.0 g, 1.5 mmol, 34%). LC/MS: MS (ESI): *m/z* 956.85 $[M+H]^+$, (retention time: 2.52 min; purity 98.74%).

Step-4: Synthesis of (S)-2-amino-N1-((3-(4-aminoquinazolin-7-yl)phenyl)sulfonyl)succinamide-trifluoroacetic acid: To a stirred solution of tert-butyl-tert-butyl(7-{m-[(*S*)-2-[tert-butyl(oxycarbonylamino)]-3-(N-tritylcarbamoyl)propionylaminosulfonyl]phenyl}-4-quinazolinyl)(oxycarbonylamino)formylate (250 mg, 261 µmol) in dichloromethane (2 mL, 31.2 mmol) under $N_2$ atmosphere was added trifluoroacetic acid (2 mL) at 0 ºC. The resulting reaction mixture was stirred at RT for 16 h. The reaction mixture was monitored by TLC and LCMS. Upon completion, After completion, reaction mixture was concentrated under reduced pressure to get crude compound which was further purified by prep-HPLC in TFA buffer to get (*S*)-2-amino-N1-((3-(4-aminoquinazolin-7-yl)phenyl)sulfonyl)succinamide- trifluoroacetic acid (1/1) (89 mg, 165 µmol, 63%) as a white solid. $^1H$ NMR (400 MHz, DMSO-*d*6): δ 9.81 (brs, 2H), 8.88 (s, 1H), 8.53 (d, *J* = 8.7 Hz, 1H), 8.23 (t, *J* = 1.6 Hz, 1H), 8.13 (dd, *J* = 8.7, 1.6 Hz, 1H), 8.02 (d, *J* = 1.6 Hz, 1H), 7.97-7.92 (m, 3H), 7.79 (s, 3H), 7.66 (t, *J* = 15 Hz, 1H), 7.59 (brs, 1H), 7.14 (s, 1H), 3.72 (brs, 1H), 2.76 (dd, *J* = 17, 3.5 Hz, 1H), 2.44 (dd, *J* = 15.3, 8.7 Hz, 1H). LC/MS: MS ($ESI^+$): *m/z* 415.05 $[M+H]^+$, (retention time: 4.83 min; purity 99.68%).

Preparation of the Thr adduct of LG-0020866:

Step-1: Synthesis of tert-butyl ((2*S*,3*R*)-1-((6-bromopyridine)-2-sulfonamido)-3-(tert-butoxy)-1-oxobutan-2-yl)carbamate: To a stirred solution of (*N*-(tert-butoxycarbonyl)-O-(tert-butyl)-L-threonine (348 mg, 1.0 eq, 1.27 mmol) in 5 mL dichloromethane in a 30 mL glass vial were added EDCI.HCl (364 mg, 1.5 eq., 1.9 mmol) and HOAt (258 mg, 1.5 eq., 1.9 mmol) at room temperature under N2 atmosphere. The reaction mixture was stirred for 30 minutes at the same temperature and then added 6-bromo-2-pyridinesulfonamide **1** (0.3 g, 1.0 eq, 1.27 mmol) and DBU (567 µL, 3 eq., 3.8 mmol) at room temperature. The resulting reaction mixture was stirred at room temperature for 16 h.The reaction mixture was monitored by TLC and LCMS. Upon completion, the reaction mixture was quenched with water (20 mL) and extracted with dichloromethane (2 × 30 mL). The combined organic layers were washed with brine, dried over anhydrous sodium sulfate, filtered, and concentrated under reduced pressure to afford the crude product which was purified by flash column chromatography using a 12 g snap column, eluting with 40–50% ethyl acetate in heptane to afford the title compound as an off-white solid (0.330 g, 1.5 mmol, 49%). LC/MS: MS (ESI): *m/z* 49175 $[M-H]^+$, (retention time: 1.56 min; purity 93.57%).

Step-2: Synthesis of tert-butyl ((2*S*,3*R*)-1-((6-(4-aminoquinazolin-7-yl)pyridine)-2-sulfonamido)-3-(tert-butoxy)-1-oxobutan-2-yl)carbamate: To a stirred the solution of (tert-butyl ((2*S*, 3*R*)-1-((6-bromopyridine)-2-sulfonamido)-3-(tert-butoxy)-1-oxobutan-2-yl)carbamate (0.3 g, 1.0 eq, 625 µmol) and 7-(4,4,5,5-tetramethyl-1,3,2-dioxaborolan-2-yl)-4-quinazolinylamine (254 mg, 1.5 eq., 937 µmol) in a 30 mL reaction vial in 1,4-dioxane (15 mL, 176 mmol) & water (1.5 mL, 83.3 mmol) was added dipotassium carbonate (259 mg, 3 eq., 1.87 mmol) under $N_2$ atmosphere at room temperature. The reaction mixture was purgged with $N_2$ gas for 5 min and then added $Pd(PPh_3)_4$ (72.2 mg, 0.1 eq., 62.5 µmol)) The resulting reaction mixture was again purgged with $N_2$ gas for 15 min., and then stirred at 100 °C for 3 h. The reaction mixture was monitored by TLC and LCMS. Upon completion, the reaction mixture was quenched with water (50 mL) and extracted with dichloromethane (2 × 100 mL). The combined organic layers were washed with brine, dried over anhydrous sodium sulfate, filtered, and concentrated under reduced pressure to afford the crude product. The crude material was purified by flash column chromatography using a 12 g snap column, eluting with 10–15% ethyMeOH in DCM, to afford the title compound as an off-white solid (0.070 g, 19%). LC/MS: MS (ESI): *m/z* 559.24 $[M+H]^+$, (retention time: 1.85 min; purity 97.75%).

Step-3: Synthesis of (2*S*, 3*R*)-2-amino-N-((6-(4-aminoquinazolin-7-yl)pyridin-2-yl)sulfonyl)-3-hydroxybutanamide.TFA: To a stirred solution of tert-butyl ((2S,3R)-1-((6-(4-aminoquinazolin-7-yl)pyridine)-2-sulfonamido)-3-(tert-butoxy)-1-oxobutan-2-yl)carbamate (70 mg, 1.0 eq, 125 µmol) in a 30 mL reaction vial in dichloromethane (15 mL, 176 mmol) was added trifluoroacetic acid (192 µL, 20 eq., 2.51 mmol) at 0 °C under $N_2$ atmosphere. The resulting reaction mixture was stirred at room temperature for 16 h. The reaction was monitored by TLC and LCMS. After completion, the reaction mixture was evaporated under reduced pressure to get crude compound which was triturated with n-pentane (3X 10 mL) and purified by prep-HPLC using TFA buffer to afford the title compound as an off white solid (0.032 g, 62%). $^1H$ NMR (400 MHz, DMSO-$d_6$): δ 9.57 (brs, 2H), 8.82 (s, 1H), 8.54-8.51 (m, 2H), 8.44-8.42 (m, 1H), 8.27 (d, $J$ = 8.0 Hz, 1H), 8.13 (t, $J$ = 7.6 Hz, 1H), 8.02 (d, $J$ = 7.6 Hz, 1H), 7.73 (bs, 2H), 5.15 (brs, 1H), 3.92-3.86 (m, 1H), 3.22 (d, $J$ = 5.2 Hz, 1H ), 1.12 (d, $J$ = 6.4 Hz, 3H). LC/MS: MS ($ESI^+$): *m/z* 403.10 (M+H), (RT: 4.52 min and Purity 99.88%).

Preparation of the Asn adduct of LG-0020866

Step-1: Synthesis of (S)-1, 2-bis (N-6-bromo-2-pyridylsulfonylcarbamoyl) ethyl 2-methyl-2-propanecarbamate: To a stirred solution of (*S*)-2-[tert-butyl(oxycarbonylamino)]-3-(N-tritylcarbamoyl) propionic acid (1.5 g, 1.5 eq.,

3.16 mmol) in 5 mL dichloromethane in a 30 mL glass vial were added EDC.HCl (606 mg, 1.5 eq., and 3.16 mmol) and HOAt (431 mg, 1.5 eq., 3.16 mmol) at room temperature under $N_2$ atmosphere. The reaction mixture was stirred for 30 minutes at the same temperature and then added 6-bromo-2-pyridinesulfonamide (0.5 g, 1 eq., 2.11 mmol) and DBU (963 mg, 3 eq., and 6.33 mmol) at room temperature. The resulting reaction mixture was stirred at room temperature for 16 h. The reaction mixture was monitored by TLC and LCMS. Upon completion, the reaction mixture was quenched with water (50 mL) and extracted with dichloromethane (2 × 100 mL). The combined organic layers were washed with brine, dried over anhydrous sodium sulfate, filtered, and concentrated under reduced pressure to afford the crude product. The crude material was purified by flash column chromatography using a 12 g snap column, eluting with 80–100% ethyl acetate in heptane to afford the title compound as an off-white solid (0.7 g, 35%). LC/MS: MS (ESI): *m/z* 692.85 $[M+H]^+$, (retention time: 4.83 min; purity 72.20%).

Step-2: Synthesis of (S)-1, 2-bis [N-6-(4-amino-7-quinazolinyl)-2-pyridylsulfonylcarbamoyl] ethyl 2-methyl-2-propanecarbamate: To a stirred the solution of (*S*)-1,2-bis(N-6-bromo-2-pyridylsulfonylcarbamoyl) ethyl 2-methyl-2-propanecarbamate (1 g, 1.44 mmol) and 7-(4,4,5,5-tetramethyl-1,3,2-dioxaborolan-2-yl)-4-quinazolinylamine (586 mg, 1.5 eq., 2.16 mmol) in a 30 mL reaction vial in 1,4-dioxane (15 mL, 176 mmol) & water (1.5 mL, 83.3 mmol) was added dipotassium carbonate (598 mg, 3 eq., 4.33 mmol) under $N_2$ atmosphere at room temperature. The reaction mixture was purged with $N_2$ gas for 5 min and then added $PdCl_2$dppf.DCM (118 mg, 0.1 eq., 1.44 mmol). The resulting reaction mixture was again purged with $N_2$ gas for 15 min., and then stirred at 100 °C for 3 h. The reaction mixture was monitored by TLC and LCMS. Upon completion, the reaction mixture was quenched with water (50 mL) and extracted with dichloromethane (2 × 100 mL). The combined organic layers were washed with brine, dried over anhydrous sodium sulfate, filtered, and concentrated under reduced pressure to afford the crude product. The crude material was purified by flash column chromatography using a 12 g snap column, eluting with 10–10% ethyl acetate in heptane, to afford the title compound as an off-white solid (0.38 g, 28%).

LC/MS: MS (ESI): *m/z* 758.05 $[M+H]^+$, (retention time: 1.96 min; purity 79%).

Step-3: Synthesis of (*S*)-2-amino-N1-((6-(4-aminoquinazolin-7-yl) pyridin-2-yl) sulfonyl) succinamide-trifluoroacetic acid: To a solution of (*S*)-1,2-bis[N-6-(4-amino-7-quinazolinyl)-2-pyridylsulfonylcarbamoyl] ethyl 2-methyl-2 propanecarbamate (380 mg, 1 eq., 5 mmol) in 4M dioxane in HCl (5.54 g, 2 eq., 152 mmol) (5 mL) at 0 °C in a 30 mL reaction vial and stirred the reaction mixture at room temperature for 16h. Reaction was monitored by TLC and LCMS data. After completion of reaction, reaction mixture was evaporated under reduced pressure to get crude compound which was triturated with diethyl ether (5 X 10 mL) and was purified by prep-HPLC in TFA buffer to get N-[6-(4-amino-7-quinazolinyl)-2-pyridylsulfonyl] (*S*)-2-aminosuccinamide (0.035 g, 17%) as an off white solid. $^1H$ NMR (400 MHz, DMSO-$d_6$): $\delta$ 9.57 (brs, 2H), 8.81 (s, 1H), 8.55-8.53 (m, 2H), 8.43 (d, $J$ = 8.72 Hz, 1H), 8.26 (d, $J$ = 7.84, 1H), 8.14-8.10 (m, 1H), 7.98 (d, $J$ = 7.72 Hz, 1H), 7.79 (brs, 2H), 7.62 (s, 1H), 7.16 (s, 1H), 3.76-3.74 (m, 1H), 2.90-2.86 (dd, $J$ = 8.72, 2.5 Hz, 1H), 2.46-2.42 (m, 1H). LC/MS: MS ($ESI^+$): *m/z* 416.05 (M+H), (RT: 4.75 min and Purity 98.18%).

### Supplementary References

1. Schmitt E, Moulinier L, Fujiwara S, Imanaka T, Thierry JC, Moras D. Crystal structure of aspartyl-tRNA synthetase from *Pyrococcus kodakaraensis* KOD: archaeon specificity and catalytic mechanism of adenylate formation. Embo J. 1998;17(17):5227-37

2. Xie SC, Wang Y, Morton CJ, Metcalfe RD, Dogovski C, Pasaje CFA, et al. Reaction hijacking inhibition of *Plasmodium falciparum* asparagine tRNA synthetase. Nat Commun. 2024;15(1):937
3. Baragaña B, Hallyburton I, Lee MCS, Norcross NR, Grimaldi R, Otto TD, et al. A novel multiple-stage antimalarial agent that inhibits protein synthesis. Nature. 2015;522(7556):315-20
